# Assembly of a segrosome by a CTP-independent ParB-like protein

**DOI:** 10.1101/2024.05.08.592561

**Authors:** Kirill V. Sukhoverkov, Francisco Balaguer-Perez, Clara Aicart-Ramos, Abbas Maqbool, Govind Chandra, Fernando Moreno-Herrero, Tung B. K. Le

**Affiliations:** Department of Molecular Microbiology John Innes Centre, Norwich, NR4 7UH, United Kingdom; Department of Macromolecular Structures Centro Nacional de Biotecnología, Consejo Superior de Investigaciones Científicas, Madrid, Spain; Department of Biochemistry and Metabolism John Innes Centre, Norwich, NR4 7UH, United Kingdom

## Abstract

The ATP– and CTP-dependent ParA-ParB-*parS* segrosome is a macromolecular complex that segregates chromosomes/plasmids in most bacterial species. CTP binding and hydrolysis enable ParB to slide on DNA and to bridge and condense DNA, thereby dictating the size and dynamics of the tripartite ParAB*S* complex. Several other evolutionarily distinct systems can also segregate DNA, although the full diversity of bacterial DNA partition systems is not yet known. Here, we identify a CTP-independent ParAB*S* system that maintains a conjugative plasmid SCP2 in the filamentous bacterium *Streptomyces coelicolor*. We demonstrate that an SCP2 ParB-like protein, ParT, loads onto DNA at an 18-bp *parS* site and diffuses away to the adjacent DNA despite lacking an apparent CTPase domain and detectable NTPase activity. We further show that *parS* DNA stimulates ParT transition from loading to a diffusing state to accumulate on DNA, and ParT activates the ATPase activity of its cognate partner protein ParA. We also identify numerous structural homologs of ParT, suggesting that CTP-independent diffusion on DNA might be widespread in bacteria despite being previously unappreciated. Overall, our findings uncover a CTP-independent DNA translocation as an alternative and unanticipated mechanism for the assembly of a bacterial DNA segregation complex and suggest that CTP binding and hydrolysis is not a fundamental feature of ParAB*S*-like systems.

## INTRODUCTION

DNA segregation is a fundamental process in biology that ensures faithful inheritance of chromosomes and plasmids. In bacteria, chromosomes and low-copy plasmids are actively transported to daughter cells by several evolutionarily distinct DNA partition systems (1–4). However, the full diversity of bacterial DNA partition systems is not yet known.

Among known bacterial DNA partition systems, the ATP and CTP-dependent type-I ParA-ParB-*parS* segrosome is the most widespread (1, 5). Type-I ParAB*S* system consists of a centromere-like *parS* DNA sequence on the cargo DNA, a *parS*-binding CTPase protein ParB, and a force-generating ATPase protein ParA (1, 2). Type-I ParAB*S* segregates DNA by transporting a ParB-coated *parS* DNA cargo along a gradient of ParA-ATP via a Brownian ratchet mechanism (2, 6–11). Briefly, ParB binds ParA and stimulates the ATPase activity of ParA, thereby dissociating ParA-ATP dimer into individual ParA-ADP/apo monomers that no longer bind the nucleoid, creating a local gradient of ParA-ATP with the least nucleoid-bound ParA-ATP near the ParB-*parS* DNA complex. The ParB-*parS* DNA complex then diffuses up the gradient by a Brownian-ratchet mechanism to rebind ParA-ATP, resulting in the net transportation of the ParB-*parS* DNA. Local depletion of nucleoid-bound ParA-ATP creates a ‘no-return’ zone which enforces the unidirectional movement of the ParB-*parS* DNA complex. Repeating cycles of ParB-ParA-DNA interactions result in an overall long-range directional transportation of the ParB-*parS* complex, and subsequently of the whole DNA cargo (12–20).

Canonical type-I ParB is a self-loading DNA clamp that binds a 16-bp *parS* sequence on the bacterial chromosome or plasmid (12–14). CTP-binding induces ParB self-dimerization at its N-terminal domain to close the clamp (12, 21, 22). Clamp closure allows *parS* DNA to transit from the ParB DNA-binding domain to a DNA-storing lumen, essentially enabling ParB to escape high-affinity binding to the *parS* site allowing it to slide along the DNA to neighboring regions (12, 13, 19, 21, 22). Repeated ParB loading onto *parS*, followed by escaping and sliding away, results in multiple ParB-CTP clamps decorating the vicinity of the *parS* locus. Slow CTP hydrolysis reverses the switch to release the clamp from DNA, thus preventing ParB from diffusing too far from *parS* (19, 21, 22). ParB-CTP has also been shown to phase-separate (17, 23) and recruit other ParB molecules to bridge and condense DNA surrounding *parS* sites (15, 20, 24). Altogether, this creates a high local concentration of ParB at the *parS* locus to activate the ATPase activity of ParA, thereby releasing ParA from the nucleoid to create the ‘no return’ zone (12–20) to segregate replicated DNA to daughter cells (6, 25).

The established picture is that CTP binding and hydrolysis allow ParB to switch between multiple modes (loading at *parS* vs. sliding on DNA vs. DNA bridging/condensing/phase separation vs. releasing from DNA), which influence the size and activity of the ParA-ParB-*parS* segrosome. The discovery of a CTP cofactor and the distinct CTP-dependent conformations adopted by ParB raised unsolved questions. For example, what are evolutionary principles that have led to the preference for regulation by CTP versus other nucleotide triphosphates-dependent switches? Are there alternative DNA clamping and ParB-like spreading mechanisms that do not utilize CTP?

Here, we uncover a CTP-independent ParB-like protein, ParT (previously known as SCP2.04c), which is encoded on a low-copy circular plasmid SCP2 in *Streptomyces coelicolor* (26). The *parT* gene and an immediate upstream *parA* gene were previously shown to contribute to the maintenance of SCP2 in *S. coelicolor* (26), however, how ParT and ParA mechanistically segregate SCP2 is unknown. By combining biochemistry, single-molecule *in vitro* reconstitution, chromatin immunoprecipitation with deep sequencing, and AlphaFold2-based structure prediction, we reveal that the N-terminal peptide of ParT binds and activates the ATPase activity of its ParA partner. We further show that ParT is a clamp-like protein that diffuses and accumulates on DNA independently of nucleotide triphosphates. We map the 18-bp ParT-loading *parS* site on SCP2 and show that ParT diffusion and accumulation on DNA are dependent on, and stimulated by, the presence of its cognate *parS* site. Additionally, we identify ∼100 structural homologs of ParT that are widespread among bacterial species, however, those associated with a ParA-like ATPase partner are rare with only nine found in sequenced genomes. Altogether, our findings provide unanticipated insights into a new but rare mode of bacterial DNA segregation that employs a CTP-independent DNA translocase.

## RESULTS

### ParT is a type-I ParB-like protein that lacks a CTPase domain

To search for ParB-like proteins with novel features, we compiled a list of genetically validated ParAB*S* systems that have been previously described in the literature and inspected their predicted structures using AlphaFold2. Among the candidates, *S. coelicolor* SCP2.04c stands out as having the common features of a type-I ParB such as a predicted flexible N-terminal peptide, a middle (M) domain with a helix-turn-helix motif, a C-terminal dimerization (C) domain, and a flexible linker connecting M and C domain. However, SCP2.04c lacks a CTPase fold at the N-terminal region (**Fig. 1A** and **Fig. S1**), leading us to rename this truncated ParB-like protein as ParT, to distinguish it from the canonical type-I ParB protein.

**Fig. 1.**
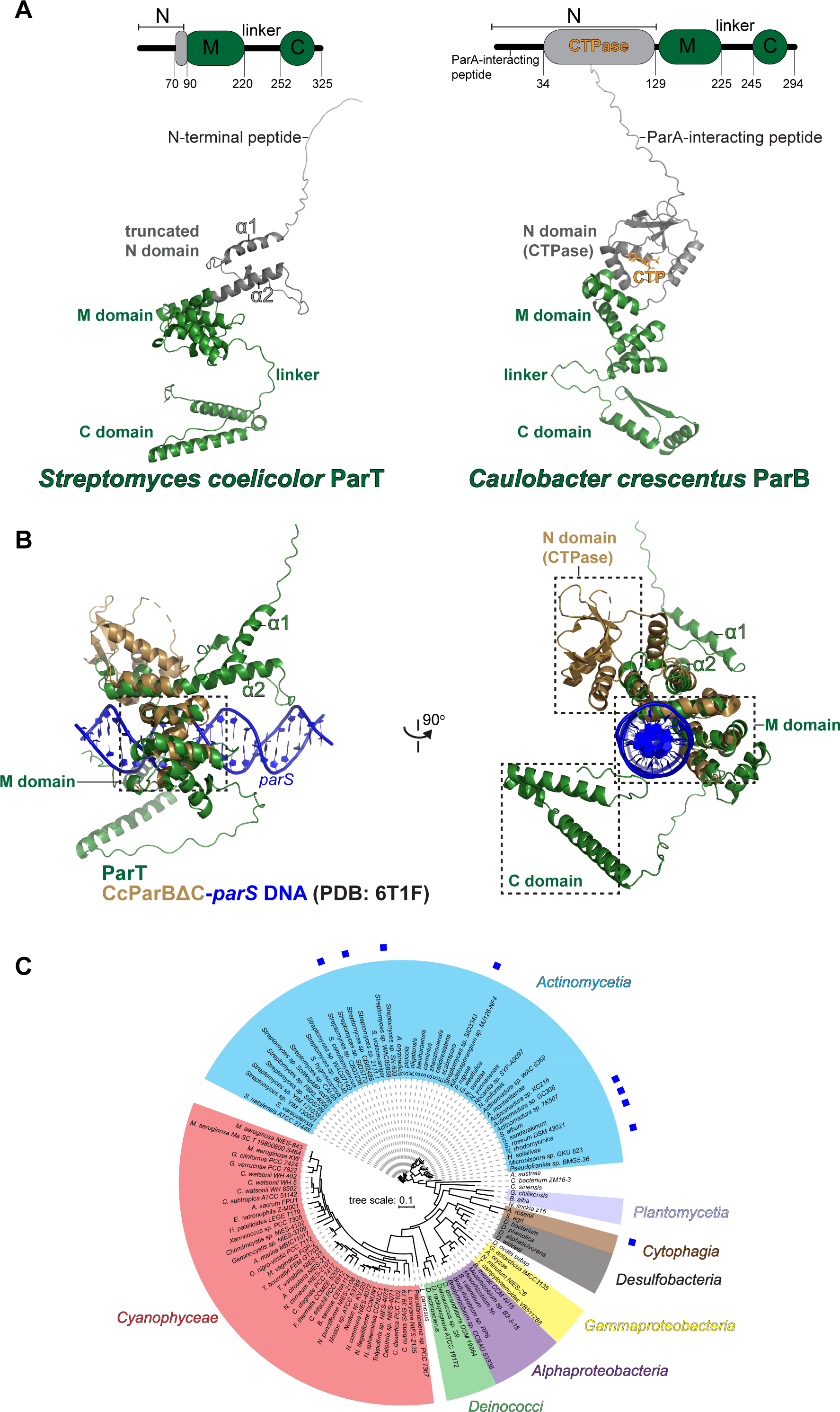
*S. coelicolor* ParT is a type-I ParB-like protein that lacks a CTPase domain. (**A**) The domain architecture of *S. coelicolor* ParT (left panel, UniProt ID: Q8VWE5) and *Caulobacter crescentus* ParB (right panel, UniProt ID: B8GW30) shows the N-terminal domain (N, grey), the middle centromere-binding domain (M, green), a C-terminal domain (C, green), and a linker connecting M and C domains. A ParA-interacting motif resides at the N domain of *C. crescentus* ParB. Below the schematic diagrams are AlphaFold2-predicted structures of ParT and *C. crescentus* ParB. Individual N and M domains of an AlphaFold2-predicted structure of *C. crescentus* ParB align well with co-complex X-ray crystallography structures of the *C. crescentus* ParBΔC (lacking a C domain) with either CTPɣS (PDB: 7BM8) or *parS* (PDB: 6T1F). **(B)** Alignment of an AlphaFold2-predicted structure of ParT to *C. crescentus* ParBΔC-*parS* DNA complex (PDB: 6T1F) suggests that the ParT M domain is likely a DNA-binding domain. **(C)** A 16S rRNA phylogenetic tree that shows the distribution of ParT structural homolog discovered by FoldSeek. Nine cases where a ParA-encoding gene was found in the genomic vicinity surrounding *parT* homologs are indicated by blue squares.

ParT and a *parS*-bound *Caulobacter crescentus* ParBΔCTD (lacking a C domain, PDB: 6T1F) (22) align closely at the M domain, suggesting that ParT M domain is also likely responsible for DNA binding (**Fig. 1B**). The N-terminal (N) domain of ParT is short, consisting of a predicted flexible N-terminal peptide, a helix α1, and part of a long helix α2 (**Fig. 1A-B**). Notably, the predicted α1 has a low model confidence score (pLDDT < 50) (**Fig. S1**), while the predicted α2 is of very high confidence (pLDDT > 90) and is a common feature in structural homologs of ParT (**Fig. S1** and **Fig. 1C**).

To assess the prevalence of ParT, we employed FoldSeek to search for structural homologs of ParT in the AlphaFold Protein Structure Database (27). We retrieved 142 unique ParT-like structures where high-quality genomes of the associated bacteria are also available (**Supp. Dataset 1**). These ParT structural homologs were found in many bacterial species, mostly in the *Actinomycetia* (47 species) and *Cyanophyceae* phyla (38 species) (**Fig. 1C**). Next, to determine whether ParT structural homologs might be part of a DNA partition system, we searched for ParA-encoding gene in the vicinity of the *parT* homologs, finding nine such cases, with eight belonging to the *Actinomycetia* phylum (**Fig. 1C**, blue squares). Overall, these data suggest that *S. coelicolor* ParT is an atypical type-I ParB protein that lacks an apparent CTPase domain, and that ParT homologs that might constitute DNA partition systems are rare and confined to actinomycetes.

### ParT interacts with ParA in the presence of ATP and non-specific DNA to activate the ATPase activity of ParA

To investigate whether ParT, despite lacking the CTPase domain, binds ParA (**Fig. 2A**), we examined the interaction of ParT with ParA in the presence or absence of ATP and non-specific DNA substrate in real-time using bio-layer interferometry (BLI). Biotinylated ParT was immobilized on a streptavidin-coated probe, and BLI monitored the shift in wavelength resulting from the change in probe optical thickness during the association and dissociation of ParA from immobilized ParT (**Fig. 2B**). In the presence of 1 µM of purified ParA alone or with 1 mM ATP or 0.05 mg/mL non-specific DNA, no BLI signal was observed (**Fig. 2B**), suggesting that ParT did not bind to apo-ParA or ParA-ATP complex. However, premixing 1 µM ParA with ATP and non-specific DNA increased the BLI response significantly (**Fig. 2B**), indicating that ParT binds to a ParA-ATP-DNA complex, consistent with the proposed role of ParT in DNA segregation (26). ParB proteins bind their partner ParA via a short positively charged N-terminal peptide (6, 28, 29). We found that the ParTΔN3 variant lacking the first three amino acids (SRR) did not show a detectable response to ParA in the presence of ATP and non-specific DNA (**Fig. 2C**), indicating that similar to ParB, the N-terminal peptide of ParT is also essential for interaction with ParA.

**Fig. 2.**
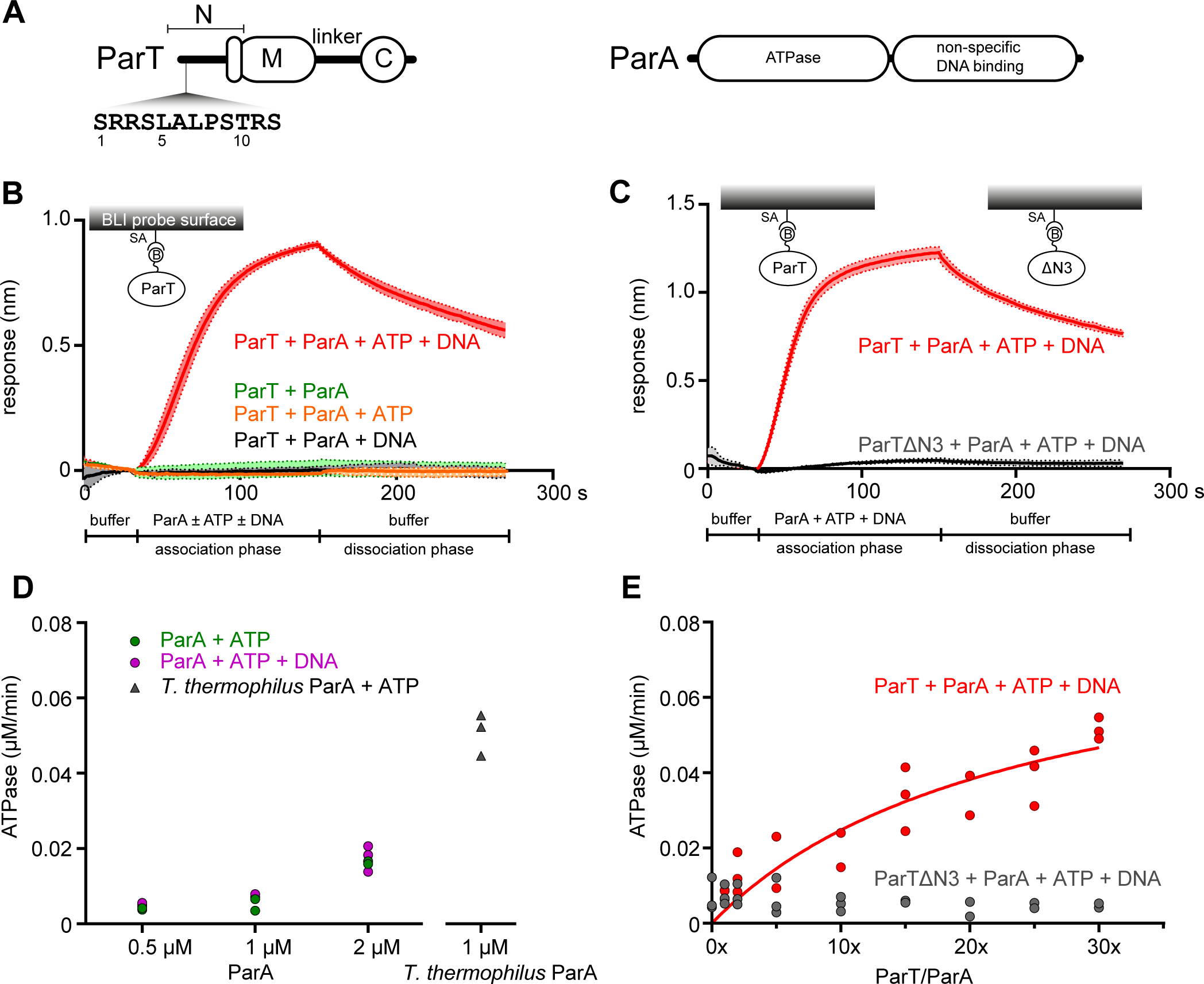
ParT interacts with ParA in the presence of ATP and non-specific DNA to activate the ATPase activity of ParA. (**A**) The domain architecture of ParT (same as Fig. 1A) and ParA. The sequence of the first 12 amino acids of ParT is also shown. The predicted ATPase and non-specific DNA-binding domain of ParA are also indicated on the diagram. **(B)** ParT binds ParA in the presence of ATP and non-specific DNA. Bio-layer interferometry (BLI) analysis of the interaction between 1 µM ParA ± 1 mM ATP ± 0.05 mg/mL salmon sperm DNA and biotinylated ParT that were immobilized on streptavidin (SA) BLI probes. Mean and standard deviation (shading) from three replicates are shown. **(C)** The first three amino acids are required for ParA-ParT interaction. BLI analysis of the interaction between 1 µM ParA + 1 mM ATP + 0.05 mg/mL salmon sperm DNA and biotinylated ParT (WT) or a ParTΔN3 lacking the first three amino acids. Biotinylated ParT and ParTΔN3 were immobilized on streptavidin (SA) BLI probes. Mean and standard deviation (shading) from three replicates are shown. **(D-E)** ParT stimulates the ATPase activity of ParA. Compared to a robust ATPase activity from *Thermus thermophilus* chromosomal ParA, SCP2 ParA by itself hydrolyzed ATP weakly regardless of the presence of DNA. **(E)** A molar excess of ParT, but not ParTΔN3, activated the ATPase activity of SCP2 ParA. All experiments were triplicated.

Next, to investigate whether ParA-ParT interaction promotes ATP hydrolysis by ParA, we measured the ATPase activities of ParA in the presence of ATP, non-specific DNA, and an increasing concentration of ParT. In the absence of ParT, 1 µM ParA did not detectably hydrolyze ATP regardless of the presence of DNA, while a positive control *Thermus thermophilus* chromosomal ParA at the same concentration showed a robust ATPase activity, consistent with a previous report (**Fig. 2D**) (6). However, with increasing molar excess of ParT over 1 µM ParA, ATPase activity was robustly detected and reached a maximal rate of ∼0.05 µM ATP/min (**Fig. 2E**). The addition of ParTΔN3, however, did not result in the activation of ParA ATPase activity (**Fig. 2E**), likely owing to its inability to interact with ParA (**Fig. 2C**). Altogether, these data show that ParT, similar to a type-I ParB, interacts with ParA-ATP-DNA to activate the ATPase activity of its cognate partner ParA.

### ParT binds a specific *parS* site on SCP2

Next, to determine the *parS* site on SCP2, we performed chromatin immunoprecipitation with deep sequencing (ChIP-seq) to map DNA bound by a FLAG-tagged ParT which was expressed ectopically from a =BT1 phage integration site on the *S. coelicolor* A3(2) chromosome (**Fig. 3A**).

**Fig. 3.**
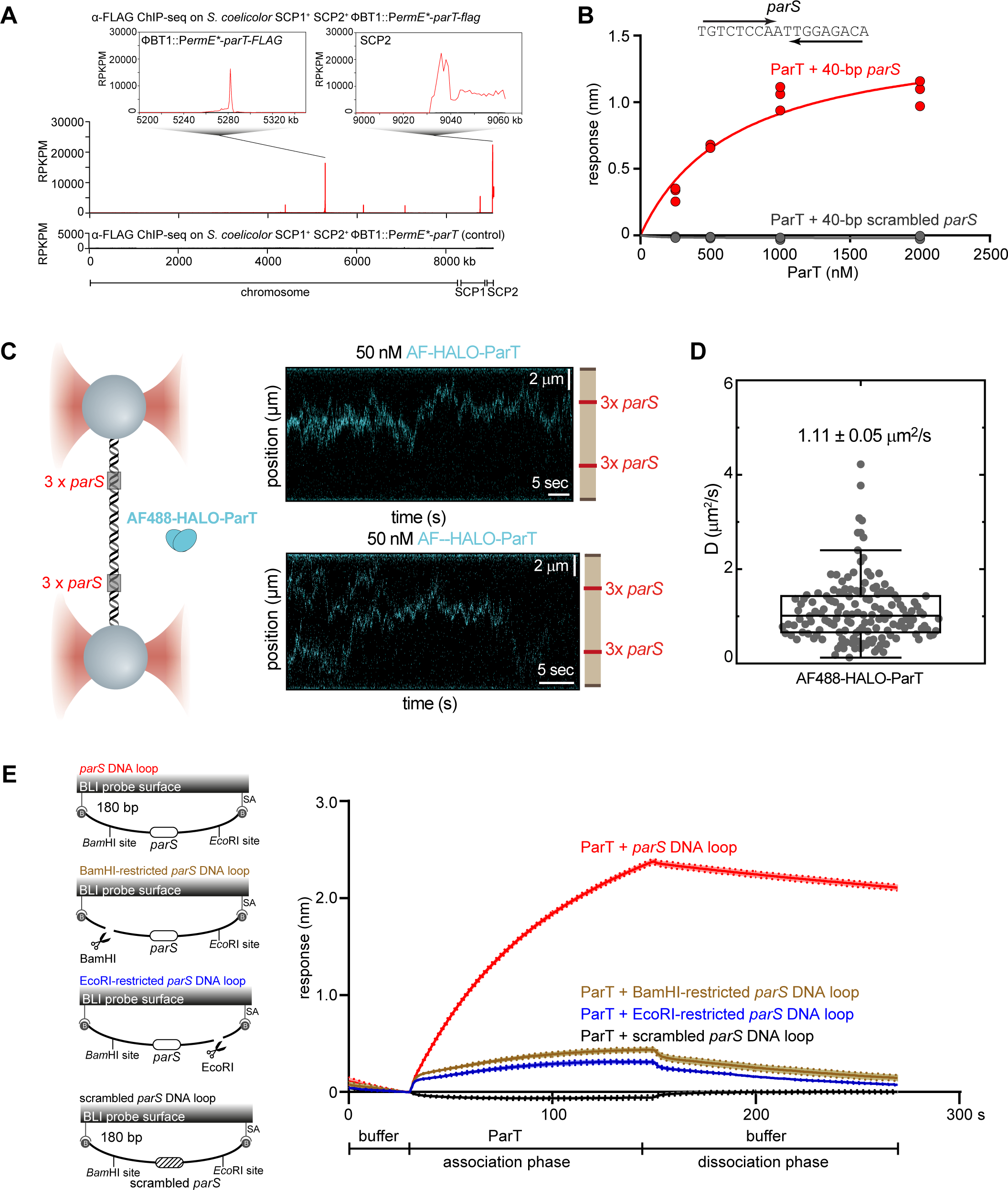
ParT diffuses on a *parS*-containing DNA. (**A**) ParT binds a specific 18-bp *parS* site on SCP2. α-FLAG ChIP-seq profiles show the enrichment of FLAG-tagged ParT on the SCP2 plasmid (inset) and at the ϕBT1 phage integration site on the chromosome (inset). Profiles were plotted with the x-axis representing genomic positions and the y-axis representing the number of reads per kilobase pair per million mapped reads (RPKPM). ChIP-seq experiments were performed twice using biological replicates, and a representative profile is shown. **(B)** BLI analysis of the interaction between increasing concentrations of ParT and 40-bp DNA duplexes containing either an 18-bp *parS* site or a scrambled *parS* site. BLI responses (nm) were defined as the maximal BLI signal during the association phase. A nucleotide sequence of *parS* is also shown, and convergent arrows indicate that *parS* is an inverted repeat. **(C)** ParT diffuses on *parS* DNA without an additional NTP co-factor. (left panel) Schematic of the C-trap optical tweezers experiments where a ∼40-kb DNA containing two clusters of 3x *parS* were tethered between two beads and scanned with a confocal microscope using 488 nm illumination. (right panel) Two representative kymographs showing AlexaFluor AF488)-HALO-ParT binding and diffusing on *parS*-containing DNA. **(D)** Boxplot showing the diffusion constant of individual ParT diffusion on *parS* DNA (*D* = 1.11 ± 0.05 µm^2^/s, mean ± standard error of measurement (SEM), n = 154) (see also **Fig. S2A-B**). **(E)** ParT can accumulate on a closed *parS* DNA loop but not on a *parS* DNA with an open end. BLI analysis of the interaction between 1 µM ParT with a 180-bp dual biotin-labeled DNA that contains either a *parS* or a scrambled *parS* site. Interactions between a dual biotinylated DNA and a streptavidin (SA)-coated probe created a closed DNA loop where both ends were blocked (see the schematic diagram of the BLI probes on the left panel). 180-bp *parS* DNA loops were subsequently restricted by EcoRI or BamHI to generate an open free end. Mean and standard deviation (shading) from three replicates are shown.

*S. coelicolor* with a non-tagged ParT expressed from the same integration site was also employed as a negative control to eliminate false signals that might arise from cross-reaction with the α-FLAG antibody (**Fig. 3A**). ChIP-seq revealed enrichment over the length of SCP2 with a summit on the *parT* gene on this plasmid, and another prominent asymmetrical peak at the ectopic *parT-flag* allele engineered at the =BT1 phage integration site on the chromosome (**Fig. 3A**). Given that the ChIP-seq data suggested that the *parS* site might reside within the coding sequence of ParT, we closely inspected the nucleotide sequence directly underneath the ChIP-seq summits, revealing an 18-bp inverted repeat that is likely to be the core *parS* sequence (**Fig. 3B**). To verify the *parS* sequence, we assessed the binding of purified ParT to a 40-bp linear DNA duplex containing the 18-bp inverted repeat using BLI. Our data showed ParT binding specifically to *parS*-containing DNA duplexes (K_D_ = 680 ± 200 nM), but not to scrambled *parS* DNA duplexes (**Fig. 3B**), consistent with the ChIP-seq data.

### ParT diffuses and accumulates on a *parS* DNA loop independently of CTP

Next, we further investigated ParT-DNA interaction by employing a longer 44.8-kb *parS*-containing DNA and dual optical tweezers combined with confocal fluorescence microscopy (30). Here, individual *parS*-or non-*parS*-containing DNA molecules were immobilized between two polystyrene beads and extended almost to their contour length under force (**Fig. 3C**). To amplify the signal of ParT DNA-binding, three *parS* sites were engineered at two clusters on the DNA (**Fig. 3C**). We incubated DNA with 50 nM of AlexaFluor (AF) 488-labeled HALO-tagged ParT and took confocal images over time to construct kymographs. When a non-*parS* DNA was employed, no fluorescence signal was observed on DNA (not shown), consistent with the requirement of *parS* for the loading of ParT on DNA. When a *parS* DNA was employed, kymographs (over 40 to 80 sec) showed fluorescence signal from AF488-HALO-ParT outside of the *parS* clusters and along the length of the DNA (**Fig. 3C**), suggesting that ParT was now distributed non-specifically along the *parS* DNA. Next, we measured the position of individual AF488-HALO-ParT along the DNA over time to determine a diffusion constant (*D*) for ParT of 1.11 ± 0.05 µm^2^/sec (**Fig. 3D** and **Fig. S2A-B**). This rate is comparable to the diffusion constants previously determined for *B. subtilis* ParB-CTP and a ParB-like KorB-CTP diffusion on DNA (30, 31). These data indicate that ParT can diffuse along a *parS* DNA substrate *in vitro* without additional nucleotide triphosphate (NTP) co-factors.

To investigate whether ParT diffusion leads to its accumulation on *parS* DNA, we followed the interaction of ParT with *parS* DNA in real-time using BLI, which allowed us to use a higher concentration of purified ParT than with our optical tweezers setup. We employed a 180-bp dual biotin-labeled *parS* or non-*parS* DNA tethered at both ends to a streptavidin-coated probe to form a closed DNA loop (14, 19, 21, 22, 32) (**Fig. 3E**). With a non-*parS* DNA substrate, no BLI signal was detectable when the probe was incubated with 1 µM ParT, consistent with a *parS*-specific loading of ParT (**Fig. 3E**). With a cognate *parS* DNA substrate, an incubation with 1µM ParT resulted in a significant BLI response (**Fig. 3E**). Previous reports demonstrated that CTP strongly promotes the association of type-I ParB proteins onto closed *parS* DNA substrates (14, 21, 30, 33). Here, premixing ParT with ATP, CTP, GTP, or UTP did not change the BLI profile markedly (**Fig. S2C**), consistent with ParT lacking an apparent CTPase domain as well as any detectable NTPase activity (**Fig. S2D**).

Next, we investigated whether a DNA substrate with a free end could support high ParT association. The 180-bp dual biotin-labeled DNA loop was designed with a BamHI and an EcoRI recognition site flanking the *parS* site. The DNA-loop probe was immersed in either a BamHI-or an EcoRI-containing buffer to enable DNA restriction to generate a free end (**Fig. 3E**). We found that incubation of a restricted DNA probe with purified ParT reduced the BLI signal by ∼fivefold, suggesting that ParT diffuses on the DNA and, similar to canonical ParB (14), slides off the free DNA end.

### ParT does not condense *parS* DNA *in vitro*

The canonical type-I ParB was previously reported to bridge and condense *parS* DNA *in vitro* in the presence of CTP (20, 24, 30). We wondered whether ParT could also condense DNA, despite lacking a CTPase domain. To investigate this, we employed magnetic tweezers which allow simultaneous measurement of multiple DNA molecules and application of a lower force than optical tweezers. Single DNA molecules harboring a cluster of 5x *parS* sites were immobilized between a glass surface and magnetic beads (**Fig. S3A-B**). Forces in the 1-5 pN range were applied to the beads to stretch *parS* DNA between a pair of magnets before the addition of buffer only or 500 nM of purified ParT. The force was subsequently lowered gradually to 0.002 pN i.e., permissive to DNA condensation, and the extension of the tethered DNA was monitored in real-time. We observed that, while *B. subtilis* ParB robustly condenses DNA in the presence of 2mM CTP, there was no difference in the DNA extension over a range of force in the presence or absence of ParT (**Fig. S3C**), indicating that ParT did not condense DNA under the tested condition.

### ParT self-dimerization is promoted by *parS* DNA *in vitro*

In the presence of CTP, ParB self-dimerizes at the N-terminal CTPase domain to create a clamp-like molecule that diffuses on DNA (18, 21, 22, 33). We wondered whether ParT could form a functionally similar clamp, despite lacking a CTPase domain. To investigate this possibility, we attempted to crystallize apo-ParT or a ParT-*parS* co-complex but were unsuccessful. Given that apo-ParT exists as a dimer in solution (**Fig. S4A**), we instead employed AlphaFold2 Multimer to predict a structure of a ParT dimer to shed light on their possible conformations (**Fig. 4A**). The top-ranking prediction from AlphaFold2 Multimer showed a high-confidence dimerization interface at the N domain of ParT, which is mediated by helices α2 (pTM = 0.71, ipTM = 0.71) and a second dimerization interface at the C domain (pTM = 0.9, ipTM = 0.89) (**Fig. S4B**). To validate whether the opposing ParT N domains dimerize in solution, we tested for site-specific crosslinking of a purified ParT variant using the sulfhydryl-to-sulfhydryl crosslinker bis-maleimidoethane (BMOE) which covalently crosslinks cysteine residues within 8 Å of each other. Based on the AlphaFold2-Multimer-predicted structure, residue S48 at the N domain was selected and substituted by cysteine (**Fig. 4A**). The S48C substitution did not impact the loading and accumulation of ParT (S48C) on a closed DNA loop (**Fig. S5A**), while wild-type ParT, despite having an native cysteine 81 residue, crosslinked minimally under tested conditions (**Fig. S5B**). We observed that, in the absence of *parS* DNA, ∼25% ParT (S48C) could be crosslinked (lane 2, **Fig. 4B**), and the crosslinking efficiency did not change in the presence of a 40-bp scrambled *parS* DNA (Lane 4, **Fig. 4B**). However, the crosslinking efficiency increased to ∼52% (lane 3, **Fig. 4B**) when a cognate *parS* DNA was included. Furthermore, and consistent with the lack of a CTPase domain, the addition of NTP did not impact the crosslinking of ParT (S48C) (**Fig. S5C**). Altogether, these data suggest that a cognate *parS* DNA promotes the engagement of opposing ParT N domains.

**Fig. 4.**
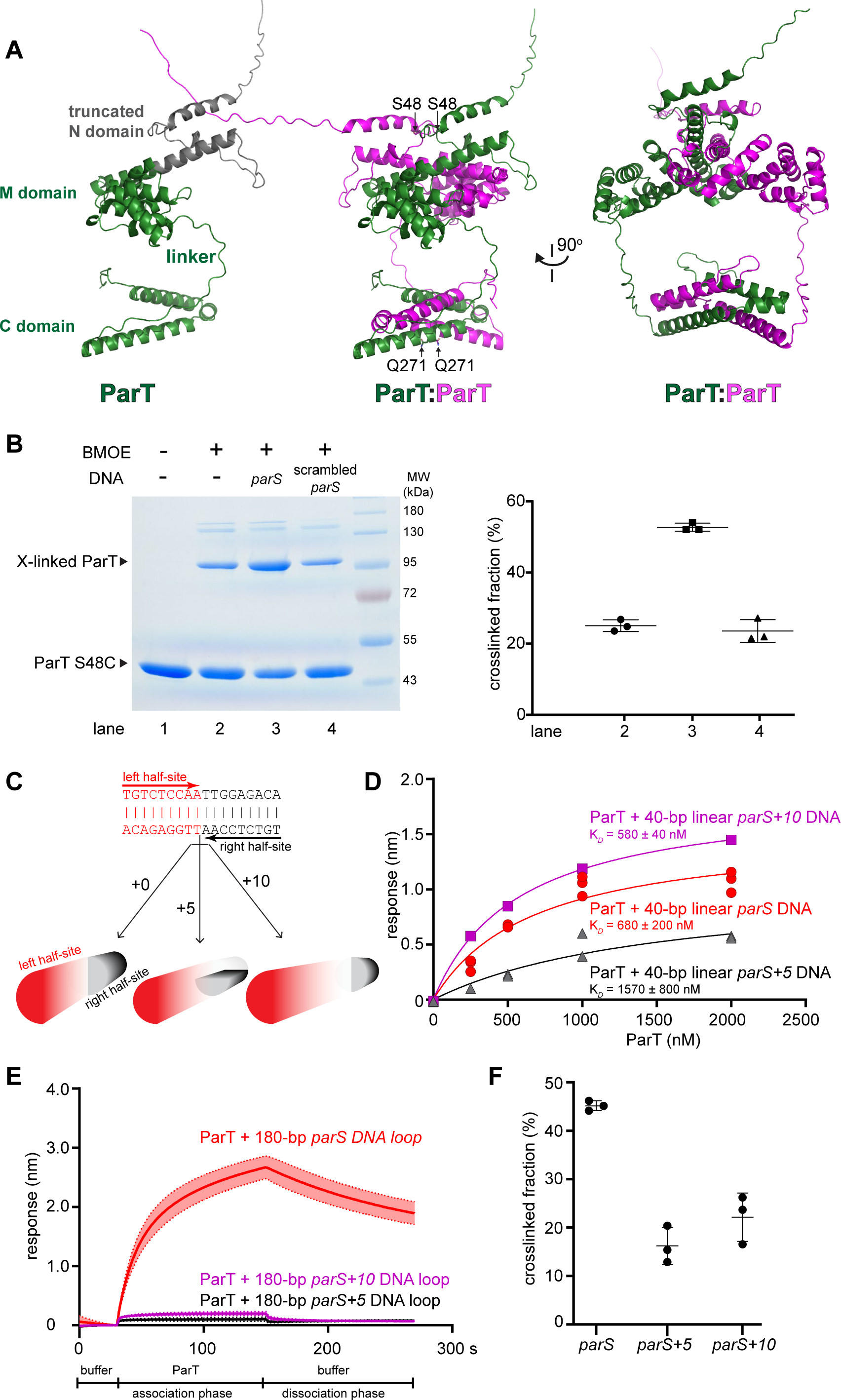
ParT self-dimerization is promoted by *parS* DNA *in vitro*. (**A**) AlphaFold2-predicted structure of ParT monomer (left panel) and ParT dimer (right panel) (see also **Fig. S4B**). Residues S48 at the N domain and Q271 at the C domain of ParT were individually substituted by cysteine for use in BMOE crosslinking assays. **(B)** SDS-PAGE analysis of BMOE crosslinking products of 4 µM ParT S48C with or without 4 μM of 40-bp *parS* DNA duplex or scrambled *parS* DNA duplex. The quantification of the major crosslinked fraction is shown on the right panel. Experiments were triplicated and mean ± standard deviation (SD) were quantified. **(C-F)** The distance between two *parS* half-sites is important for ParT accumulation but not ParT loading on a DNA loop **(C)** An insertion of 5 or 10 bp in between two *parS* half-sites (red and black arrows) rotated these half-sites by a half or a full helical turn, respectively. **(D)** BLI analysis of the interaction between increasing concentrations of ParT and 40-bp DNA duplexes harboring either a *parS* or a *parS+5* or a *parS+10* site. **(E)** BLI analysis of the interaction between 1 µM ParT with a 180-bp dual biotin-labeled DNA that contains either a *parS* or a p*arS+5* or a *parS+10* site. Experiments were triplicated and mean ± SD were quantified. **(F)** Quantification of the major BMOE-mediated crosslinked fraction of ParT (S48C) incubated with 40-bp *parS* or *parS+5* or *parS+10* DNA duplexes.

Next, to investigate the dimerization of opposing C domains of ParT, we constructed a ParT (Q271C) variant. Again, the symmetry-related Q271C residues from opposing ParT subunits were predicted to be crosslinkable based on the AlphaFold2-Multimer predicted structure of a ParT dimer (**Fig. 4A**). ParT (Q271C) showed ∼37% crosslinking efficiency, however, the presence of *parS* DNA did not change the crosslinked fraction (**Fig. S5D**). Overall, our data suggest that ParT N dimerization, but not the C dimerization, is responsive to cognate *parS* DNA.

### The distance between *parS* half-sites is important for ParT accumulation but not for ParT loading on a DNA loop

We hypothesized that *parS* binding promotes ParT(Q48C) crosslinking by optimally orienting the opposing N-M domains for self-dimerization. To test this, we engineered *parS* DNA variants with additional five or ten ‘spacer’ nucleotides in between two halves of the *parS* inverted repeats to rotate *parS* half-sites, and thus of their bound ParT subunits, by a half or a full helical turn, respectively (**Fig. 4C**). We observed that ParT binding to a 40-bp *parS*+5 or *parS*+10 DNA substrate with a similar or ∼twofold higher K_D_ than to a wild-type *parS* DNA (**Fig. 4D**), indicating that the loading of ParT onto DNA substrates was only mildly affected by the extra spacer. By contrast, when a 180-bp dual-biotin *parS* DNA was employed (**Fig. 4E**), a low BLI signal was observed when 1 µM ParT was incubated with probes carrying a 180-bp dual-biotin *parS*+5 or *parS*+10 DNA loop instead (**Fig. 4E**), indicating that ParT could no longer accumulate on such closed DNA loops. Consistent with the inability to accumulate on these DNA loops, the crosslinking efficiency of ParT (Q48C) reduced to ∼22% in the presence of *parS*+5 or *parS*+10 DNA duplexes (**Fig. 4F**). We reasoned that re-orienting *parS* half-sites uncoupled the ParT loading from ParT accumulation, likely by interfering with the ability of ParT to self-dimerize at the N domain.

## DISCUSSION

In this work, we report an atypical DNA segregation protein, ParT, that self-loads onto DNA at a specific *parS* site to diffuse and accumulate on a DNA loop independently of NTP. Results from optical tweezers experiments show no evidence of a filament-like structure of ParT on DNA, indicating that ParT is unlikely to polymerize along the DNA. Furthermore, we did not observe ParT-mediated DNA condensation, suggesting that it does not bridge distal DNA together under the tested conditions. The observation of diffusive single ParT molecules in an optical tweezers experiment and the requirement of a closed DNA loop for ParT accumulation is more consistent with a model of DNA-entrapped ParT that diffuses/slides on DNA (**Fig. 5**).

**Fig. 5.**
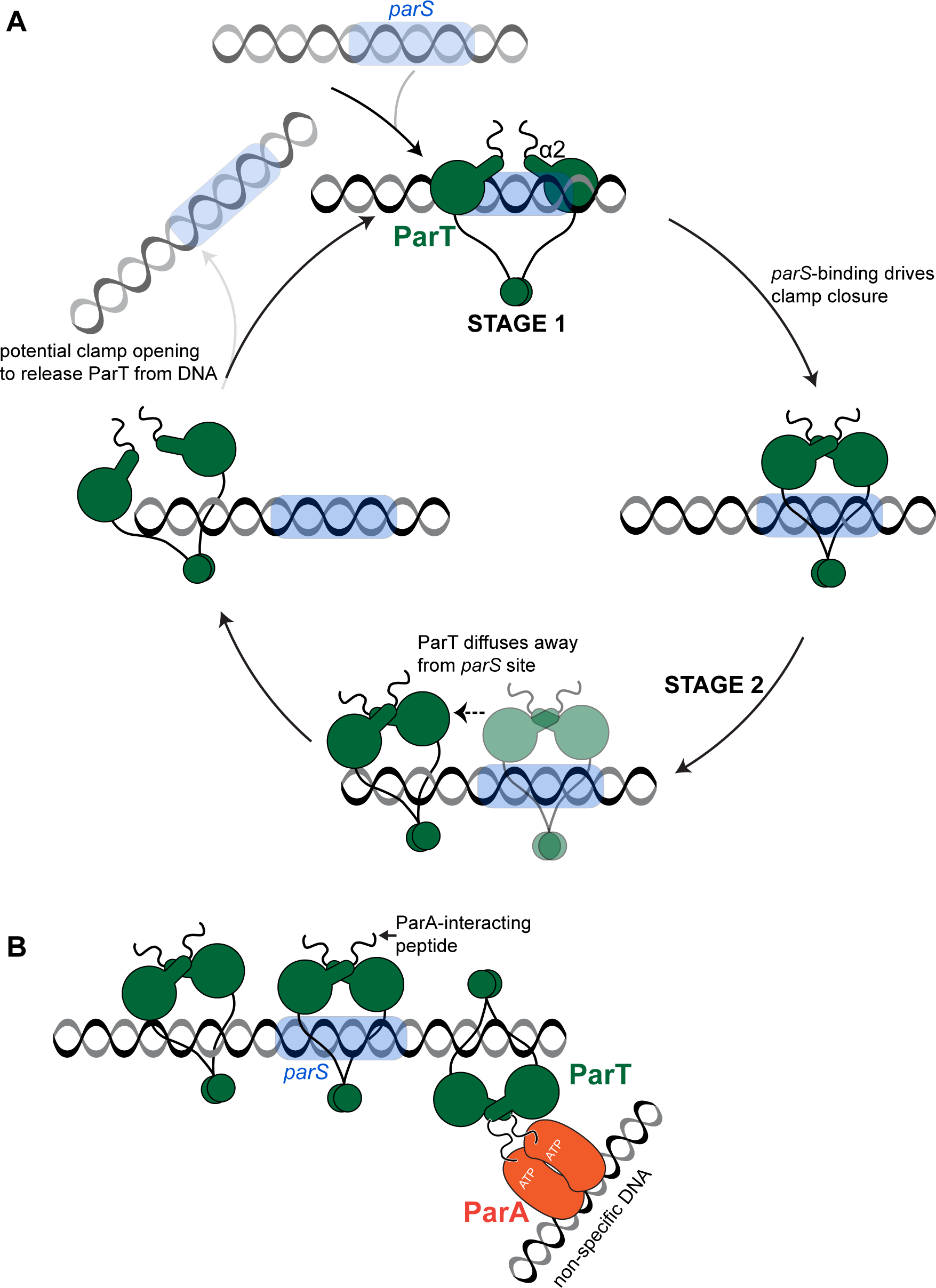
A proposed model of ParT loading and accumulation on *parS* DNA and its interaction with ParA. (**A**) A proposed model for ParT (green) loading, diffusing, and release cycle. Loading ParT is likely to be an open clamp so that the M domain is accessible to bind the *parS* (cyan) DNA site. Binding to *parS* also promotes self-dimerization of α2 at the N domain of opposing ParT subunits to close a clamp that enables ParT to escape to the neighboring DNA. A now free *parS* site is available for the next ParT to load on and subsequently diffuses away. The repeating cycles of ParT loading and diffusing away likely enable multiple ParT to accumulate surrounding *parS* site, this creates a high local concentration of ParT to activate the ATPase activity of the partner ParA protein (orange) (see also panel **B**). ParT clamp can potentially open to release from the DNA but the mechanism of release is not yet known. **(B)** ParT binds to its partner ParA (orange), via a positively-charged N-terminal peptide, in the presence of ATP and non-specific DNA. A high concentration of ParT is required to activate the ATPase activity of ParA, this is potentially achieved by the accumulation of ParT specifically in the vicinity of the *parS* site on SCP2.

The overall behavior of ParT is reminiscent of the ParB CTP-dependent DNA-sliding clamp (18, 34). Yet how ParT diffuses on DNA in the absence of a CTPase domain remains to be demonstrated. Our data are consistent with a model in which ParT can switch between an open– and a closed-clamp conformation in a *parS*-dependent manner to (i) load site-specifically at *parS* (stage 1, **Fig. 5A**) and then (ii) to escape a high-affinity *parS* site to diffuse/slide to neighboring DNA (stage 2, **Fig. 5A**). In stage 1, ParT likely adopts an open conformation so that the DNA-binding (M) domains from opposing ParT subunits are accessible to the *parS* site (**Fig. 5A**). In stage 2, ParT likely switches conformation to enable its escape from *parS* to diffuse away. In ParB, CTP enables the *parS*-loaded ParB subunits to self-dimerize their opposing CTPase (N) domains to form a closed-clamp conformation (19, 21, 22, 33). In this closed conformation, the DNA-binding (M) domains of ParB were re-orientated and became closer together, thus are no longer compatible with binding two consecutive major grooves of *parS* (22, 33). This CTP-dependent re-orientation of the *parS* DNA-binding domain enables ParB to escape from a high-affinity *parS* site to diffuse away to the adjacent DNA (14, 21, 30, 33). Despite lacking the CTPase domain, ParT might similarly form a closed-clamp conformation to escape its cognate *parS* site. AlphaFold2-Multimer predicted a structure of ParT dimer in a closed-clamp conformation in which both the N and C domains from opposing subunits self-engaged (**Fig. 4A**). The clamp-closed conformation of ParT was further supported by the crosslinking experiment that showed a twofold increase in the crosslinking efficiency of ParT S48C in the presence of *parS* DNA. We reason that *parS*-binding alone serves to stimulate the transition from an open-to a closed-clamp conformation by orienting the two opposing N-M domains of ParT, especially helices α2 and α2’, for self-dimerization to close the clamp (**Fig. S4B-C**). Indeed, altering the relative orientation of opposing ParT N-M domains by inserting five or ten nucleotides in between the two *parS* half-sites did not impact the initial ParT loading but eliminated ParT self-dimerization and the subsequent diffusion and accumulation of ParT on a DNA loop (**Fig. 4C-F**). ParT escaping from the loading site to the neighboring DNA also vacates *parS* for the next ParT loading event. Similar to the CTPase ParB, we envision that the repeated ParT loading on *parS*, followed by escape and diffusion to the neighboring DNA, results in multiple ParT molecules decorating the vicinity of *parS*.

We observed that accumulated ParT on a DNA loop could be slowly released when the BLI probe was returned to a buffer-only solution (**Fig. 3E**, dissociation phase). In the case of ParB, CTP hydrolysis regulates the switching from a closed-clamp conformation back to an open-clamp conformation, enabling ParB release from DNA (14, 19–21). But how ParT, lacking a CTPase domain, can be released from DNA, and whether the release is spontaneous or involves a dedicated *in vivo* factor, is not yet known. Future studies, especially the isolation and characterization of diffusion-competent but release-defective ParT variants, will hope to shed light on this aspect of the ParT mechanism.

The accumulation of ParT surrounding *parS* results in a high local concentration that is essential for the activation of the ATPase activity of its partner ParA protein (**Fig. 2E** and **Fig. 5B**). These are essential features, that are shared with type-I ParAB*S* systems, to ensure the ‘no return’ zone only forms where the ParAB*S* segresome is. It would be detrimental to the DNA segregation process if ParB was to activate the ATPase activity of ParA at an equimolar ratio because cytoplasmic DNA-unbound ParB would unproductively release ParA from the nucleoid at spatially random locations inside the cells. In a bigger picture, it has been noted that, despite the diversity of evolutionarily distinct DNA partition systems (type I, II, and III) with vastly different types of centromere-binding proteins (CBPs) and associated NTPases (1), there is a common feature in all systems: the requirement for a high local concentration of CBPs decorating the centromere-like *parS* region. The high local concentration of CBPs can either be functionally achieved by diffusion or oligomerization of CBP from a single *parS* site or by site-specific binding to an array of *parS* sites adjacent to each other (reviewed in (1)). In this context, ParT diffuses to accumulate on DNA independently of CTP represents another case of convergent evolution to segregate bacterial DNA.

Lastly, we noted that, while both ParT and canonical ParB employ a positively charged N-terminal peptide to interact with their cognate ParA partners, the amino acid identity of their ParA-interacting motifs are distinct (ParT: SRR motif vs. ParB: LG-R/K-GL motif). This might contribute to ensuring the SCP2 ParAT*S* segrosome does not cross-interact, thus interfering with the several coexisting ParAB*S* systems in the same cell such as the single ParAB*S* system on the linear chromosome and two other ParABS systems on the linear plasmid SCP1 (35–38).

### Final perspectives

In this work, we further reveal the diversity of bacterial DNA partition systems by demonstrating that a CTP-independent mechanism for the assembly of segrosome exits and a *parS* site alone can promote both self-loading and diffusion of an atypical ParB-like protein ParT. We reason that CTP binding and hydrolysis is not a fundamental feature of ParAB*S*-like systems, but were left wondering what had selected for dependency on CTP in some systems but not in others. It is possible that the binding and breaking down of a small molecule co-factor i.e. CTP potentially offer more regulation for the formation and dynamics of the ParAB*S* segrosome than a CTP-independent segrosome that relies solely on a spontaneous clamp opening/closing. This could be an evolutionary principle that had led to the ubiquity of the canonical CTP-dependent ParABS system in bacteria. Lastly, besides the nine ParT structural homologs that are likely to form functional ParAT*S* modules, we identified other 133 ParT structural homologs that are not associated with neighboring ParA-encoding genes, suggesting a possibility that evolution has adapted a CTP-independent DNA-sliding domain/motif to diversify proteins to perform new biological functions beyond DNA segregation.

## MATERIALS AND METHODS

### Construction of plasmids and strains

#### Construction of pET21b::parT-(his)6 (WT and mutants)

Two overlapping DNA fragments *ParT_WT_f1* and *ParT_WT_f2* containing an N-terminal or a C-terminal half of a codon-optimized version of *parT* (SCP2.04c), respectively, were chemically synthesized (gBlocks, IDT) (**Supp. Dataset 1**). The two gBlocks fragments were assembled into an NdeI-HindIII-cut pET21b backbone using a 2x NEB Gibson assembly master mix (NEB; cat# E2611). To enable such assembly, pET21b was digested with NdeI (NEB, cat# R0111S) and HindIII-HF (NEB, cat# R3104S) and purified by gel extraction. A 10 µL reaction mixture was created with 5 µL 2x Gibson master mix and 5 µL of combined equimolar concentration of purified backbone and gBlock(s). This mixture was incubated at 50°C for 60 min. Gibson assembly was possible owing to a 37-bp sequence shared between the NdeI-HindIII-cut pET21b backbone (**Supp. Table 2**) and each of the two gBlocks fragments. The mixture was introduced into *E. coli* DH5α (**Supp. Table 3**) by heat-shock transformation and carbenicillin-resistant colonies were selected. Subsequently, plasmids were isolated and verified by Sanger sequencing (Genewiz, UK). A correct plasmid was introduced into *E. coli* Rosetta (DE3) pLys (**Supp. Table 3**) to create an overexpression strain. The overexpression strains for ParT (S48C)-His_6_ and ParT (Q271C)-His_6_ were constructed similarly but using *ParT_S48C_f1* + *ParT_WT_f2*, and *ParT_WT_f1* + *ParT_Q271C_f2* gBlocks, respectively (**Supp. Dataset 1**). An overexpression strain for ParA-His_6_ was also constructed similarly but using *ParA_WT_f1* + *ParA_WT_f2* pair of gBlocks (**Supp. Dataset 1**).

#### Introducing a FLAG-tagged parT allele to Streptomyces coelicolor A3(2) genome

To introduce a FLAG tag to the C-terminus of *parT*, DNA containing *parT* was amplified by PCR from *S. coelicolor* A3(2) genomic DNA using primers *ParT_CFLAG_Fw* and *ParT_CFLAG_Rv* (**Supp. Table 1**), and subsequently purified by gel extraction. The purified PCR product was assembled with an NdeI-HindIII-cut pIJ10257 backbone using a 2x Gibson master mix (NEB). The resulting plasmids were verified by Sanger sequencing (Genewiz, UK). A plasmid carrying a non-tagged version of *parT* was constructed similarly, but primers *ParT_Fw* and *ParT_Rv* (**Supp. Table 1**) were used instead.

To integrate *parT*-FLAG or the non-tagged version onto the *S. coelicolor* A3(2) ΦBT1 phage integration site, *E. coli* ET12567 + pUZ8002 (**Supp. Table 3**) were transformed with plasmid pIJ10257::*parT-flag* or pIJ10257::*parT* (**Supp. Table 2**), and subsequently conjugated to *S. coelicolor* A3(2) (**Supp. Table 3**) as previously described (39). Conjugants were purified by re-streaking to single colonies twice on SFM medium containing 50 μg/mL of hygromycin (Merck, cat# 10843555001). Spore stocks of resulting strains were eventually prepared and stored at –80°C.

### Overexpression and purification of ParT (WT/variants)-His_6_ and ParA-His_6_

The C-terminally His-tagged ParT (WT and mutants) were expressed from the plasmid pET21b in *E. coli* Rosetta (DE3) pLys cells (Merck, UK) (**Supp. Table 3**). Overnight culture (80 mL) was used to inoculate 4 L of LB broth supplemented with carbenicillin and chloramphenicol. Cells were grown at 37°C with shaking at 200 rpm till OD_600nm_ reached ∼0.4, then cultures were cooled down to 16°C before isopropyl-β-D-thiogalactopyranoside was added to the final concentration of 0.5 mM. The culture was incubated overnight at 16°C with shaking at 200 rpm before the cells were collected by centrifugation at 5400 g at 16°C.

Pelleted cells were resuspended in 50 mL of buffer containing 100 mM tris(hydroxymethyl)aminomethane hydrochloride (Tris-HCl) pH 7.4, 300 mM sodium chloride, 10 mM imidazole and 5% (v/v) glycerol (buffer A), supplemented with 1 µL of Benzonase (Merck; cat# E1014), 5 mg of lysozyme (Merck; cat# 4403), and an EDTA-free protease inhibitor tablet (Merck; cat# 11873580001). The cell suspension was incubated on a wheel rotator at room temperature for 45 min before cell lysis by sonification (10 cycles of 15 s on/off on ice at an amplitude of 20 microns).

The cell debris was pelleted by centrifugation at 42000 g for 40 min at 4°C. The supernatant was filtered through a 0.22 µm sterile filter (Starlab; cat# E4780-1226) and loaded onto a 1 mL His-Trap column (Cytiva; cat# 17524701) that had been pre-equilibrated with buffer A (100 mM Tris-HCl pH 7.4, 300 mM sodium chloride, 10 mM imidazole, and 5% (v/v) glycerol). The column was washed with buffer A until A_280nm_ plateaued. ParT was eluted from the column by an increasing gradient of buffer B (100 mM Tris-HCl pH 7.4, 300 mM sodium chloride, 500 mM imidazole, and 5% (v/v) glycerol). ParT-containing fractions were pooled and concentrated using Amicon Ultra-15 centrifugal filter units (Merck; cat# UFC901024), and buffer exchanged to a low-salt buffer (100 mM Tris-HCl pH 8, 25 mM sodium chloride and 10% (v/v) glycerol) using PD-10 desalting columns (Cytiva; cat# 17085101).

The low-salt ParT solution was loaded onto a 1 mL Heparin HP column (Cytiva; cat# 17040601) that had been pre-equilibrated with the low-salt buffer. The column was then washed with the low-salt buffer until A_280nm_ plateaued. ParT was eluted from the column by an increasing gradient of a high-salt buffer (100 mM Tris-HCl pH 7.4, 1 M sodium chloride, 10% (v/v) glycerol). ParT-containing fractions were pooled, concentrated using Amicon Ultra-15 centrifugal filter units, and buffer exchanged to an EDTA-containing buffer (100 mM Tris-HCl pH 8, 250 mM sodium chloride, 10 mM EDTA and 10% (v/v) glycerol) using PD-10 desalting columns. Afterward, to remove traces of non-specific DNA, metal ions, or nucleotide triphosphates, the ParT solution in EDTA-containing buffer was concentrated to ∼0.5 mL and dialyzed overnight against 2 L of the EDTA-containing buffer using a dialysis tubing with a molecular weight cut-off of 10 kDa (Thermo Fisher Scientific; cat# 68100). The dialyzed protein was centrifuged at 17000 g at 4°C to remove the precipitated protein, then the supernatant was loaded onto a gel-filtration HiLoad 16/600 Superdex 200 pg column (Cytiva; cat# 28989335) pre-equilibrated with a buffer containing 10 mM Tris-HCl pH 8.0 and 250 mM sodium chloride. Eluted fractions corresponding to non-aggregated ParT were pooled, concentrated, and mixed with glycerol to the final concentration of 5% (v/v). The protein was then aliquoted, flash-frozen in liquid nitrogen, and stored at –80°C.

The ParT (S48C)-His_6_, ParT (Q271C)-His_6_, and ParA-His_6_ were prepared following a similar procedure, however, the purification on heparin columns was omitted. Additionally, to ensure that cysteine residues were reduced, ParT (S48C)-His_6_ and ParT (Q271C)-His_6_ aliquots were supplemented with 1 mM tris (2-carboxyethyl) phosphine hydrochloride (Merck; cat# C4706) before flash-freezing in liquid nitrogen.

### Biotinylation of ParT

NHS-PEG_4_-Biotin (Thermo Scientific; cat# A39259) reagent was to biotinylate ParT-His_6_ for bio-layer interferometry (BLI) assays. A 10 µL of NHS-PEG_4_-Biotin (stock: 40 mM, freshly dissolved in DMSO) was added to 0.5 mL of 40 µM ParT solution in a reaction buffer containing 100 mM HEPES pH 7.4, 300 mM sodium chloride, and 10% (v/v) glycerol. The solution was mixed by inverting the tube several times and incubated at room temperature for 30 min. Then, to remove residual unincorporated biotinylating reagent, the reaction mixture was buffer exchanged to a storage buffer containing 100 mM Tris-HCl pH 7.4, 300 mM sodium chloride, and 10% (v/v) glycerol using PD-10 desalting columns. Afterward, the protein solution was concentrated to the final concentration of ∼60 µM using Amicon Ultra-15 centrifugal filter units, aliquoted, flash-frozen in liquid nitrogen, and stored at –80°C. The same protocol was used to prepare the biotinylated ΔN3ParT variant.

### Measurement of ParA-ParT interaction by bio-layer interferometry (BLI) assay

Streptavidin (SA)-coated probes (Sartorius; cat# 18-5136) were first hydrated for 10 min in a binding buffer containing 100 mM Tris-HCl at pH 8, 150 mM sodium chloride, 1 mM magnesium chloride, and 0.005% (v/v) Tween-20. Biotinylated ParT was diluted in the binding buffer to the final concentration of 1 µM so that they could be immobilized onto the surface of the SA probe. Briefly, SA probes were attached to the BLI instrument (Fortebio) and incubated with shaking at 2200 rpm sequentially in the binding buffer for 30 s, then in 1 µM ParT solution in the binding buffer for 120 s, and lastly in the binding buffer for 120 s. To remove traces of non-specifically bound ParT, probes were incubated for another 5 min in a high-salt buffer containing 100 mM Tris-HCl at pH 8, 1 M sodium chloride, 5 mM EDTA, and 0.005% (v/v) Tween-20, followed by a 10 min incubation in the binding buffer.

To assay for ParA-ParT binding, ParA-His_6_ was first diluted in the binding buffer to the final concentration of 1 µM, and preincubated on ice for 20 min with 1 mM ATP, or 0.05 mg/mL of salmon sperm DNA (Merck; cat# D1626), or combinations thereof. Afterward, ParT-coated probes were incubated in the binding buffer for 30 s, with shaking at 2200 rpm on the BLI instrument, to establish the baseline. The probes were subsequently transferred to a solution containing apo-ParA, ParA + ATP, or ParA + ATP + DNA and incubated with shaking for 120 s. Lastly, the probe was transferred to the binding buffer only and incubated for another 120 s. All assays were performed in triplicates, and the probe was regenerated in between each replicate by a 5-min incubation in the high-salt buffer.

### Preparation of biotinylated linear *parS* DNA

A pair of 40-bp single-stranded DNA oligonucleotides: a 5’-biotinylated *parS_Fw* and a non-biotinylated *parS_Rv* (**Supp. Table 1**) were dissolved in an annealing buffer (1 mM Tris-HCl pH 8.0, 5 mM sodium chloride) to the final concentration of 100 µM. An equal volume of each oligonucleotide solution was mixed and heated at 98°C for 5 min before being left to cool down to room temperature overnight to form 50 µM double-stranded biotinylated *parS* duplex. A biotinylated scrambled *parS* duplex was also prepared similarly but using a 5’-biotinylated *scrambled parS_Fw* and a non-biotinylated *scrambled parS_Rv* oligonucleotides instead (**Supp. Table 1**).

### Measurement of protein-DNA interaction by BLI assay

A modified version of a BLI-based ParA-ParT binding assay was used to assay ParT-DNA interaction. Briefly, 40-bp biotinylated *parS* duplexes were immobilized on streptavidin-coated probes as described above, and then ParT-*parS* binding was measured in increasing ParT concentration from 0.25 to 2 µM, in three technical replicates per concentration. A 40-bp biotinylated scrambled *parS* duplex was also assayed in the same condition to assess non-specific DNA binding by ParT. To calculate the binding constants (K_D_), the maximal values of BLI signal in the association phase were plotted as a function of ParT concentration. The resulting curve was approximated using GraphPad Prism 10 non-linear regression model “One site –-Specific binding” with the default parameters using the following equation:

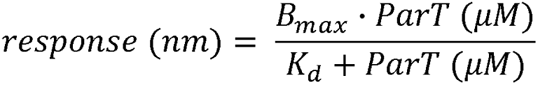

Where *response (nm)* is the maximal values of BLI signal in the association phase measured in nm, *ParT (µM)* is ParT concentration measured in µM*, K_d_* is an apparent ParT-*parS* dissociation constant measured in µM, and *B_max_*is the estimated maximum specific binding possible under the given conditions, measured in nm.

### Preparation of double-biotinylated DNA substrate

A DNA construct containing BamHI and EcoRI restriction sites, a *parS* sequence, a *tetO* sequence, *M13For* and *M13Rev* homologous regions at each end was chemically synthesized (gBlocks, IDT) (**Supp. Dataset 1**). To generate a dual biotin-labeled DNA substrate, PCR reactions were performed using a 2x GoTaq PCR master mix (Promega; cat# M7122), a 5’-biotinylated *M13For* and a 5’-biotinylated *M13Rev* primers (**Supp. Table 1**), and the gBlock fragment as a template. The PCR product from eight reactions was pooled together and purified by agarose gel extraction.

### Measurement of ParT-DNA loop interaction by BLI assay

A double biotinylated 180-bp DNA was diluted in the binding buffer (100 mM Tris-HCl at pH 8, 150 mM sodium chloride, 1 mM magnesium chloride, and 0.005% (v/v) Tween-20) to the final concentration of 1 µM and immobilized on streptavidin-coated probes by sequential incubation with shaking at 2200 rpm in the binding buffer for 30 s, then in the DNA substrate solution for 120 s, and finally in the binding buffer for 120 s. To remove the traces of non-specifically bound DNA, the probes were incubated for another 5 min in a high-salt buffer (100 mM Tris-HCl at pH 8, 1 M sodium chloride, 5 mM EDTA, and 0.005% (v/v) Tween-20), followed by a 10 min incubation in the binding buffer.

To measure ParT binding onto the 180-bp DNA loop, DNA-coated probes were incubated in the binding buffer for 30 s, with shaking at 2200 rpm on the BLI instrument, to establish the baseline and subsequently transferred to a solution containing 1 µM ParT in the binding buffer and incubated with shaking for 120 s. Lastly, the probes were transferred to the binding buffer and incubated for another 120 s. The assay was repeated three times, each time using a new probe.

To generate an open end on the left or right handside of the 180-bp DNA loop, the tips of DNA-coated probes were immersed in 1x rCutSmart buffer (NEB;cat# B6004S) containing either 400 U/mL of BamHI-HF (NEB; cat# R3136S) or EcoRI-HF (NEB; cat# R3101S) and incubated for 2.5 hrs at 37°C. A control probe was also incubated in 1x rCutSmart buffer only to account for the possible non-enzymatic DNA degradation. For each digestion, three DNA-coated probes were used.

### Assessing ParT N-terminus and C-terminus dimerization by *in vitro* BMOE cross-linking assay

ParT (S48C), ParT (Q271C), or ParT (WT), at 4 µM dimer concentration, was incubated on ice either alone or with 4 μM of 40-bp *parS* duplex or scrambled *parS* duplex in a crosslinking buffer (100 mM Tris-HCl pH 7.4, 150 mM sodium chloride, 5 mM magnesium chloride) for 5 min. To determine the crosslinking efficiency of ParT (WT/variants) in the presence of nucleotide triphosphates, the reaction crosslinking reactions were supplemented with either 1 mM of ATP (Thermo Fisher Scientific; cat# R0441), CTP (Thermo Fisher Scientific; cat# R0451), GTP (Thermo Fisher Scientific; cat# R0461) or UTP (Thermo Fisher Scientific; cat# R0471). Afterward, 20 mM DMSO solution of bismaleimidoethane (ThermoFisher; cat# 22323) was added to the reaction mixture to the final concentration of 2 mM, and the mixture was incubated at room temperature for 5 min before the crosslinking reactions were quenched by adding SDS-PAGE loading dye containing β-mercaptoethanol. The quenched samples were heated to 96°C for 3 min to denature the protein, cooled down, and loaded on 12% Tris-glycine polyacrylamide denaturing gels (ThermoFisher; cat# XP00122BOX). The crosslinked and non-crosslinked species were then resolved by electrophoresis at 150 V for 55 min. Upon completion of electrophoresis, gels were stained using an InstantBlue Coomassie solution (Abcam; cat# ab119211), and band intensity was quantified using ImageJ (NIH) (40). All experiments were performed in triplicates.

### Measurement of ParA ATPase activity by EnzChek phosphate release assay

ATP hydrolysis was monitored using an EnzCheck phosphate assay kit (Thermo Fisher Scientific; cat# E6646). ParA, ParT, and salmon-sperm DNA were mixed and pre-incubated on ice for 10 min at the following final concentrations of 5 µM, 5-150 µM, and 2 mg/mL, respectively, in a reaction buffer containing 100 mM Tris-HCl pH 7.4, 150 mM sodium chloride, and 5 mM magnesium chloride. In parallel, an NTPase assay working solution was prepared by diluting freshly defrosted aliquots of aqueous 1 mM 2-amino-6-mercapto-7-methylpurine riboside and 100 IU/mL of purine nucleoside phosphorylase in the reaction buffer to the final concentrations of 0.2 mM and 1 IU/mL, respectively. Afterward, 10 µL of the ParA-ParT-DNA mixtures were mixed with 80 µL of the assay working solution in separate wells of a 96-well plate (Cellstar; cat# 655180) and incubated for 5 min at room temperature. During this incubation, a fresh working solution of ATP was prepared by diluting 100 mM ATP (Thermo Fisher Scientific; cat# R0441) to 10 mM in the reaction buffer. ATP hydrolysis reactions were initiated by adding 10 µL of 10 mM ATP to the wells containing ParA, ParT, DNA in the working solution. Then, the plates were incubated at 25°C with constant shaking and the A_360nm_ was monitored using BioTek Eon plate reader (Agilent). Samples containing only ParA, ParT, or DNA were monitored similarly and served as controls. To convert A_360nm_ values to concentrations of inorganic phosphate, a calibration curve was generated using the same working solution. Briefly, potassium dihydrogen phosphate was added to the final concentration of 5 to 150 µM into wells containing the working solution, and A_360nm_ was monitored as described above. The results were analyzed using Excel and plotted in GraphPad Prism 9.

### Measurement of ParT NTPase activity by EnzChek phosphate release assay

NTP hydrolysis by ParT was monitored by following a similar procedure used for the measurement of ParA ATPase activity. ParT-His_6_ was diluted to the final concentration of 10 µM in either a reaction buffer or in a reaction buffer supplemented with 10 µM of 40-bp *parS* duplex, and incubated on ice for 10 min. Afterward, 10 µL of ParT or ParT-*parS* mixture was mixed with 80 µL of the assay working solution in separate wells of a 96-well plate and incubated for 5 min at room temperature. Then, an NTPase reaction was initiated by adding 10 µL of 10 mM ATP, CTP, GTP, or UTP to wells containing the ParT-*parS* mixture in the working solution and the hydrolysis kinetics was monitored using BioTek Eon plate reader. The positive control Noc-His_6_ was assayed similarly, but only CTPase activity in the presence of an equimolar amount of 22-bp *NBS* duplex was monitored.

### Chromatin immunoprecipitation with deep sequencing (ChIP-seq) and data analysis

*S. coelicolor* A3(2) strains harboring a FLAG-tagged or non-tagged *parT* alleles under the control of an *ermE\** promoter (**Supp. Table 3**) were grown as liquid cultures as follows. Firstly, the spore stocks were diluted to the final OD_450 nm_ of 0.25 in 5 mL of 0.05 M TES buffer and were incubated standing at 50°C for 10 min before adding 5 mL of 2x PG medium (1% (w/v) of yeast extract, 1% (w/v) of casamino acids and 10 mM of CaCl_2_). Then, the heat-shocked spores suspensions were incubated with shaking at 37°C for 3 hours and subsequently pelleted by centrifugation at 3000 rpm for 10 min at room temperature. The pelleted spores were resuspended in 0.5 mL of growth media comprising 40% (v/v) of YEME and 60% (v/v) TSB and then transferred to a 50 mL of the same media in a conical flask where a metal spring was fit in the bottom to facilitate a disperse growth of *Streptomyces* mycelium. The cultures were then grown overnight at 30°C with shaking. Formaldehyde (Merck; cat# F8775) was added to the cultures to the final concentration of 1% (v/v) and the cultures were incubated for 30 min at 30°C with shaking to fix the cells. The crosslinking reaction was quenched by adding glycine (Fisher Scientific; cat# 10467963) to the final concentration of 0.1 M, followed by a 10-min incubation at room temperature on a wheel rotator. Afterward, cells were collected at 8800 g for seven minutes at 4°C, washed twice with 25 mL of phosphate-buffered saline (pH 7.4), and resuspended in 1.5 mL of the same buffer before final centrifugation at 17000 g for 5 min at 4°C. The supernatant was then removed, and the pelleted cells were resuspended in 0.75 mL of the lysis buffer (20 mM of potassium 4-(2-hydroxyethyl)-1-piperazineethanesulfonic acid salt pH 7.9, 50 mM potassium chloride, 10% glycerol, 15 mg/mL lysozyme, and an EDTA-free protease inhibitor tablet).

The cell suspension was incubated in a 37°C water bath for 25 min, and then placed on ice before addition of 0.75 mL of the lysis buffer without lysozyme. Cells were lysed by sonication at 8 µm amplitude in 15 s pulses followed by 15 s pauses for eleven cycles in total using a Soniprep 150 ultrasonic sonicator (Sanyo). Cell debris was then pelleted by centrifugation at 17000 g for 20 min at 4°C, and the supernatant was transferred to a Lo-bind Eppendorf tube (Eppendorf; cat# 022431021). To prepare the supernatant for further processing, 1 M Tris-HCl (pH 8), 5 M sodium chloride, and 10% (v/v) Nonidet P-40 octyl phenoxypolyethoxylethanol (NP-40) were added to the supernatant to the final concentrations of 10 mM, 100 mM, and 0.1 % (v/v), respectively.

To immune-precipitate protein-DNA crosslinking products, 100 µL of anti-FLAG beads were pre-equilibrated in the IPP-150 buffer (10 mM Tris-HCl pH 8, 150 mM sodium chloride, 0.1% NP-40), added to the supernatant, and incubated overnight on a wheel rotator at 4°C. Subsequently, the beads were pelleted by centrifugation at 13000 g for 30 s at 4°C, and the beads were washed five times with 1 mL of IPP-150 each. Lastly, the beads were washed with 1 mL of TE buffer (10 mM Tris-HCl pH 7.4 and 1 mM EDTA).

Beads were resuspended in 150 µL of elution buffer (50 mM Tris-HCl pH 8, 10 mM EDTA, and 1% (w/v) sodium dodecyl sulfate), and incubated at 65°C for 15 min, to reverse the crosslinking. Afterward, beads were pelleted by centrifugation at 17000 g for 5 min at room temperature, and the supernatant was transferred to a 2 mL Lo-bind Eppendorf tube. The remaining beads were resuspended in 100 µL of TE buffer + 1% (w/w) of sodium dodecyl sulfate, and incubated for 5 min at 65°C. Beads were then pelleted as described above, and the supernatant was pooled together with the supernatant from the previous step, and incubated overnight at 65°C to completely reverse protein-DNA crosslinking.

Once the incubation was completed, the sample was centrifuged at 13000 g for three min at room temperature. Afterward, the supernatant was recovered and mixed with five volumes of PB buffer from QIAquick PCR purification kit (Qiagen; cat# 28104) and passed through a QIAquick PCR purification column to allow DNA binding to the membrane. The column was washed twice with 520 µL of PE buffer and finally centrifuged at 17000 g for two min to remove residual buffer. To elute DNA, 20 µL of distilled water was placed on the membrane, and the column was centrifuged at 13000 g for two min to collect the eluate in a 1.5 mL Lo-bind Eppendorf tube. The elution was repeated once more, and the eluate was pooled together and used to construct DNA libraries for Illumina deep sequencing using an NEB DNA Library Prep Kit (NEB; cat # E7645S). The prepared DNA libraries were stored at –80°C before being sequenced at the Tufts University Core Facilities (US).

For analysis of ChIP-seq data, a reference genome of *S. coelicolor* A3(2) SCP1^+^ SCP2^+^ =BT1::*parT-flag* was first constructed by concatenating the sequences of a linear chromosome (Genbank ID: AL645882.2) (with the sequence of pIJ10257::*parT-flag* plasmid inserted at the =BT1 locus), a linear plasmid SCP1 (Genbank ID: AL589148.1), and a cicular plasmid SCP2 (Genbank ID: AL645771.1) together. Subsequently, Hiseq 2500 (50 bp) or NextSeq 550 Illumina (75 bp) short reads were mapped back to this reference genome using Bowtie 1 (41) and the following command: bowtie –M 1 –n 1 –best –strata –p 4 –chunkmbs 512 A32-bowtie –sam *.fastq > output.sam. Subsequently, the sequencing coverage at each nucleotide position was computed using BEDTools (42) using the following command: bedtools genomecov –d –ibam output.sorted.bam –g A32.fna > coverage_output.txt. When necessary, MACS2 were employed to call peaks (43), for example, using the following command: macs2 callpeak –t./exp/output.sorted.bam –c./control/output.sorted.bam –f BAM –g 8e+6 –-nomodel –n expvscontrol. ChIP-seq profiles were plotted with the x-axis representing genomic positions and the y-axis is the number of reads per base pair per million mapped reads (RPBPM) or number of reads per kb per million mapped reads (RPKPM) using custom R scripts. For the list of ChIP-seq datasets in this study, see **Supp. Table 4**.

### Identification of ParT structural homologues by FoldSeek

The amino acid sequence of ParT (UniProt database entry: Q8VWE5) was used to build an AlphaFold2 model of a ParT dimer, which was then used to query FoldSeek for structurally similar proteins in the AlphaFold Database (44, 45). Afterward, upon removal of duplicate entries, we used a bespoke Perl script utilizing the Net::FTP module to download associated genomic assemblies where these were available in the NCBI bacterial genome assemblies collection. To search for structural homologs of ParT that possibly constitute a ParAB*S* system, we created a consensus ParA sequence based on the collection of experimentally confirmed ParA enzymes (**Supp. Dataset 2**) and used BLAST2.13.0 search (46) with an expectation value E = 1×10^−9^; followed by enforcing the percentage of identity to 30% or more and percentage of coverage of queries and subjects to 50% or more to identify ParA-like enzymes localized on the same contigs with ParT structural homologs. Lastly, to establish an evolutionary relationship between ParT structural homologs, we selected the genome assemblies that have 16S rRNA gene included and built a phylogenetic tree using FastTree2 with default parameters (47).

### Mass photometry measurements of ParT oligomeric state

All mass photometric measurements were recorded at 25°C using the Refeyn OneMP mass photometer (Refeyn Ltd, UK). The instrument was calibrated with a set of calibrants, β-amylase (56, 112, and 224 kDa), BSA (66 kDa), and urease (90.7, 272, and 545 kDa). A stock solution of 1 µM ParT-His_6_ in a buffer containing 10 mM Tris-HCl pH 8.0 and 250 mM sodium chloride was diluted in PBS buffer on a coverslip to the final concentrations of 100 nM, 50 nM, and 25 nM. Movies were recorded by using AcquireMP software (version R1) for 60 sec with a frame rate of 60 per sec and using a large field of view. The data were processed using DiscoverMP software (version R1.2). The mass of ParT-His_6_ was estimated by fitting a Gaussian distribution into mass histograms and taking the value at the mode of the distribution.

### Design and construction of a large plasmid with 3x *parS* sites for confocal optical tweezers (C-trap) experiments

The large DNA plasmid containing 3 copies of the 22 bp-inverted repeat sequence *parS* (CGTGTCTCCAATTGGAGACATC) was produced as follows. First, a large DNA plasmid with a single *parS* site was fabricated by ligating a dsDNA duplex containing a single copy of the *parS* site into a large plasmid previously prepared in our laboratory that did not contain this site. The dsDNA duplex with the *parS* site was obtained by annealing two oligonucleotides (**Supp. Table 1**) by heating at 95°C for 5 min and cooling down to 20°C at a rate of –1 °C min^−1^ in hybridization buffer (10 mM Tris-HCl pH 8.0, 1 mM EDTA, 200 mM NaCl, and 5 mM MgCl_2_) followed by a phosphorylation step of the 5’-terminal ends by the T4 PNK (NEB). This dsDNA duplex was ligated into the large plasmid digested with KpnI (NEB) and dephosphorylated with rSAP (NEB). This resulted in a large plasmid with a single *parS* site. To fabricate the plasmid with 3x *parS* sites, the previous large plasmid containing 1x *parS* site was digested with NruI (NEB) and dephosphorylated. A new dsDNA duplex was prepared by annealing two oligonucleotides containing 2x *parS* sites separated by 40 bp. These oligonucleotides were designed to contain several internal restriction sites flanking the *parS* sites. These restriction sites were used to digest the duplex with different pairs of restriction enzymes generating different ends. The diverse dsDNA duplexes were used in different ligation steps. For this specific cloning, the dsDNA duplex once annealed was digested with SfoI and HpaI (NEB). The digested dsDNA duplex was ligated into the linearized large plasmid, resulting in a large plasmid with 3x *parS* sites (20622 bp). Plasmids were introduced into *E. coli* DH5α competent cells and potentially positive colonies were then selected by colony PCR (48). Plasmids were purified from the cultures using a QIAprep Spin Miniprep Kit (QIAGEN), analyzed by restriction enzyme digestion, and finally verified by Sanger sequencing. This plasmid was used to produce a C-trap dsDNA construct (see below).

### Design and construction of a DNA plasmid with 5x *parS* sites for magnetic tweezers (MT) experiments

A DNA plasmid containing 5x *parS* sites was produced by following several cloning steps. First, a plasmid derived from pUC19 plasmid previously prepared in our laboratory was digested with KpnI and PshAI (NEB) followed by dephosphorylation, and the vector fragment of 6190 bp was gel extracted (QIAGEN). On the other hand, a small fragment containing a single *parS* site was produced by PCR amplifying a region of the large plasmid containing 1x *parS* sites described above with Phusion High-Fidelity DNA Polymerase (Thermo Scientific) (**Supp. Table 1**). The PCR fragment was then digested with KpnI in one end, followed by phosphorylation of the blunt end with T4 PNK and ligated into the 6190 bp vector fragment. This resulted in a plasmid of 6417 bp with 1x *parS* site.

To increase the number of *parS* sites the annealed two long oligonucleotides containing 2x *parS* sites described above for the C-trap large plasmid (**Supp. Table 1**) were employed. To fabricate a plasmid with 3x *parS* sites, the dsDNA duplex with 2x *parS* sites digested with SfoI and HpaI was ligated into the plasmid with 1x *parS* site digested with SfoI and dephosphorylated. This resulted in a plasmid with 3x *parS* sites (6507 bp). To fabricate a plasmid with 5x *parS* sites, the dsDNA duplex with 2x *parS* sites digested with XhoI and SalI (NEB) was ligated into the plasmid with 3x *parS* site digested with XhoI and dephosphorylated. This resulted in the final plasmid with 5x *parS* sites (6614 bp). The plasmids were cloned and analyzed as described for C-trap plasmids. This plasmid with 5x *parS* sites was used to prepare a C-magnetic tweezers dsDNA construct (see below).

### C-trap dsDNA construct with 3x *parS* sites

C-trap experiments were performed on dsDNA molecules of 20622 bp containing 3x *parS* sites. The central part of the dsDNA construct was obtained by digestion of the large C-trap plasmid (see above) with NotI (NEB) produced following published protocols (48). Without further purification, the fragment was ligated to highly biotinylated handles of ∼1 kb ending in PspOMI. Handles for C-trap constructs were prepared by PCR (**Supp. Table 1**) as described for biotin-labeled MT handles. These handles were highly biotinylated to facilitate the capture of DNA molecules in the C-trap experiments. As both sides of the DNA fragment end in NotI, it is possible to generate tandem (double-length) tethers flanked by the labeled handles. The sample was ready for use in C-trap experiments without further purification. The DNAs were not exposed to intercalating dyes or UV radiation during their production and were stored at 4°C. The sequence of the central part of the C-trap construct is included in **Supp. Table 5**.

A control dsDNA construct of 20482 bp without *parS* sites was similarly prepared but using as central part the fragment corresponding to the linearization of the large plasmid previously prepared in our laboratory that did not contain any *parS* site that was employed to fabricate the large plasmids with *parS* sites. The sequence of the central part of this control C-trap construct is included in **Supp. Table 5**.

### Magnetic tweezers dsDNA construct with 5x *parS* sites

The dsDNA construct for magnetic tweezers experiments consisted of a central dsDNA fragment of 6602 bp containing 5x *parS* sites, obtained by digestion with NotI and ApaI (NEB) of the final MT plasmid described above, flanked by two highly labeled DNA fragments, one with digoxigenins and the other with biotins, of 997 bp and 140 bp, respectively, used as immobilization handles. The biotinylated handle was shorter to minimize the attachment of two beads per DNA tether. Handles for MT constructs were prepared by PCR (**Supp. Table 1**) with 200 µM final concentration of each dNTP (dGTP, dCTP, dATP), 140 µM dTTP, and 66 µM Bio-16-dUTP or Dig-11 dUTP (Roche) using plasmid pSP73-JY0 (49) as template, followed by digestion with the restriction enzyme ApaI or PspOMI (NEB), respectively. The Dig-PspOMI handle was dephosphorylated with rSAP to fabricate non-torsionally constrained tethers. Labeled handles were ligated to the central part overnight using T4 DNA Ligase (NEB). The sample was then ready for use in MT experiments without further purification. The DNAs were never exposed to intercalating dyes or UV radiation during their production and were stored at 4°C. The sequence of the central part of the MT construct is included in **Supp. Table 5**.

### Confocal optical tweezers experiments

Confocal-optical tweezers experiments were carried out using a dual optical tweezers setup combined with confocal microscopy and microfluidics (C-Trap; Lumicks). A computer-controlled stage allowed rapid displacement of the optical traps within a five-channel fluid cell, allowing the transfer of the tethered DNA between different channels separated by laminar flow. Channel 1 contained 4.38 µm streptavidin-coated polystyrene beads (Spherotech). Channel 2 contained the DNA substrate harboring 3x *parS* sequences or the control DNA without *parS* both of them labeled with multiple biotins at both ends. DNA and beads were diluted in 20 mM HEPES pH 7.8, 100 mM KCl, and 5 mM MgCl_2_. A single DNA tether was assembled by first capturing two beads in channel 1, one in each optical trap, and fishing for a DNA molecule in channel 2. The tether was then transferred to channel 3 filled with reaction buffer (100 mM Tris pH 8, 100 mM NaCl, 1 mM MgCl_2,_ and 1 mM DTT) to verify the length of the correct length of the DNA by force-extension curves. The DNA was then incubated in channel 4 filled with 1 μM ParT. To reduce the fluorescence background in single ParT diffusion measurements, imaging was occasionally performed in channel 3 after protein incubation in channel 4.

To visualize the AF488-HALO-ParT, A 488 nm laser was used to excite AF488-HALO-ParT and the emission was detected with a 500-525 nm filter. Protein-containing channels were passivated with BSA (0.1% w/v in PBS) for 30 min before the experiment. Kymographs were generated by single line scans between the two beads using a pixel size of 100 nm and a pixel time of 0.1 ms, resulting in a typical time per line of 22.4 ms. The confocal laser intensity at the sample was 2.2 µW. Experiments were performed in constant-force mode at 15 pN.

For the calculation of the diffusion coefficient, we used custom Python scripts to access, visualize, and export confocal data from Bluelake (Lumicks) HDF5 files obtained from C-Trap experiments. The quantification of the individual ParT trajectories was done by using a custom LabVIEW software that provides the position of individual proteins along the DNA for a given time (t). The length of the time courses was restricted to 2.5 sec to increase the statistical sample. The mean square displacement (MSD) was then calculated for a given time interval (Δt) and the diffusion coefficient (*D*) was obtained as described in (30, 50, 51). A total of 154 trajectories were used for the diffusion coefficient calculation.

### Magnetic tweezers experiments

Magnetic tweezers experiments were performed using a homemade setup that was previously described (30, 52). Briefly, optical images of micron-sized superparamagnetic beads tethered to a glass surface by DNA substrates were acquired using a 100x oil immersion objective and a CCD camera operating at 120 Hz. Real-time image analysis allows the spatial coordinates of the beads to be determined with nm accuracy in the x, y, and z directions. We controlled the stretching force of the DNA by using a step motor coupled to a pair of magnets located above the sample. The applied force is quantified from the Brownian motion of the bead and the extension of the tether, obtained by direct comparison of images taken at different focal planes (53, 54).

Magnetic tweezers experiments were performed as follows. First, a double PARAFILM (Sigma)-layer flow chamber was incubated at 4°C overnight with 25ng/μL Digoxigenin Antibody (Bio-Rad) that adsorbed onto the polystyrene-covered lower surface. During the experiment 8μL of a DNA containing 5x *parS* (1.4nM) was diluted 1:300 in TE (10mM Tris pH 8, 1 mM EDTA) and mixed with 20 μl of 1 μm-diameter magnetic beads (Dynabeads Myone Streptavidin T1, Thermo Fisher Scientific) diluted 1:10 in PBS-BSA (0.4 mg/mL BSA (NewEngland Biolab)). After 10 minutes the excess of DNA in solution was removed by precipitating the beads with a magnet and discarding all the supernatant volume. Beads were then washed three times with fresh PBS-BSA and resuspended in 80 μl before flushing into the flow cell. The DNA-beads were then incubated in the chamber for 15 minutes to promote the interaction of the digoxigenin handle with the anti-digoxigenin surface. The excess beads were washed away by flushing around 1 ml PBS. Torsionally constrained molecules and beads containing more than a single DNA molecule were identified from their distinct rotation-extension curves and discarded for further analysis. Force-extension curves were generated by measuring the extension of the tethers at decreasing forces from 5.5 pN to 0.006 pN. The curves were first measured on naked DNA molecules and then the experiment was repeated using 1 μM of *Bacillus subtilis* ParB or 1 μM ParT in the reaction buffer (100 mM Tris pH 8, 100 mM NaCl, 1 mM MgCl_2_, 1 mM DTT and 0.1 mg/mL BSA) supplemented with 2 mM CTP. Data were analyzed and plotted using Origin software.

## DATA AND MATERIALS AVAILABILITY

All relevant data are within the manuscript and supplementary files. ChIP-seq data are available at GEO (accession code: GSE263222). Plasmids and strains generated in this study are available upon request.

## ACKNOWLEDGEMENTS

We thank Dr Wieland Steinchen for help with preliminary experiments, and Jovana Kaljević for helpful comments on the manuscript. This work is supported by the Lister Institute fellowship (T.B.K.L), the Wellcome Trust Investigator grant 221776/Z/2/Z to T.B.K.L that funds K.V.S, and the BBSRC-funded Institute Strategic Program Harnessing Biosynthesis for Sustainable Food and Health (HBio) (BB/X01097X/1). F.M.H acknowledges support from grant PID2020-112998GB-I00, funded by MICIU/AEI/10.13039/501100011033. F.B.P. acknowledges support from grant FJC2020-044824-I, funded by MICIU/AEI /10.13039/501100011033, and European Union Next Generation EU/PRTR.

---

**Fig. S1.**
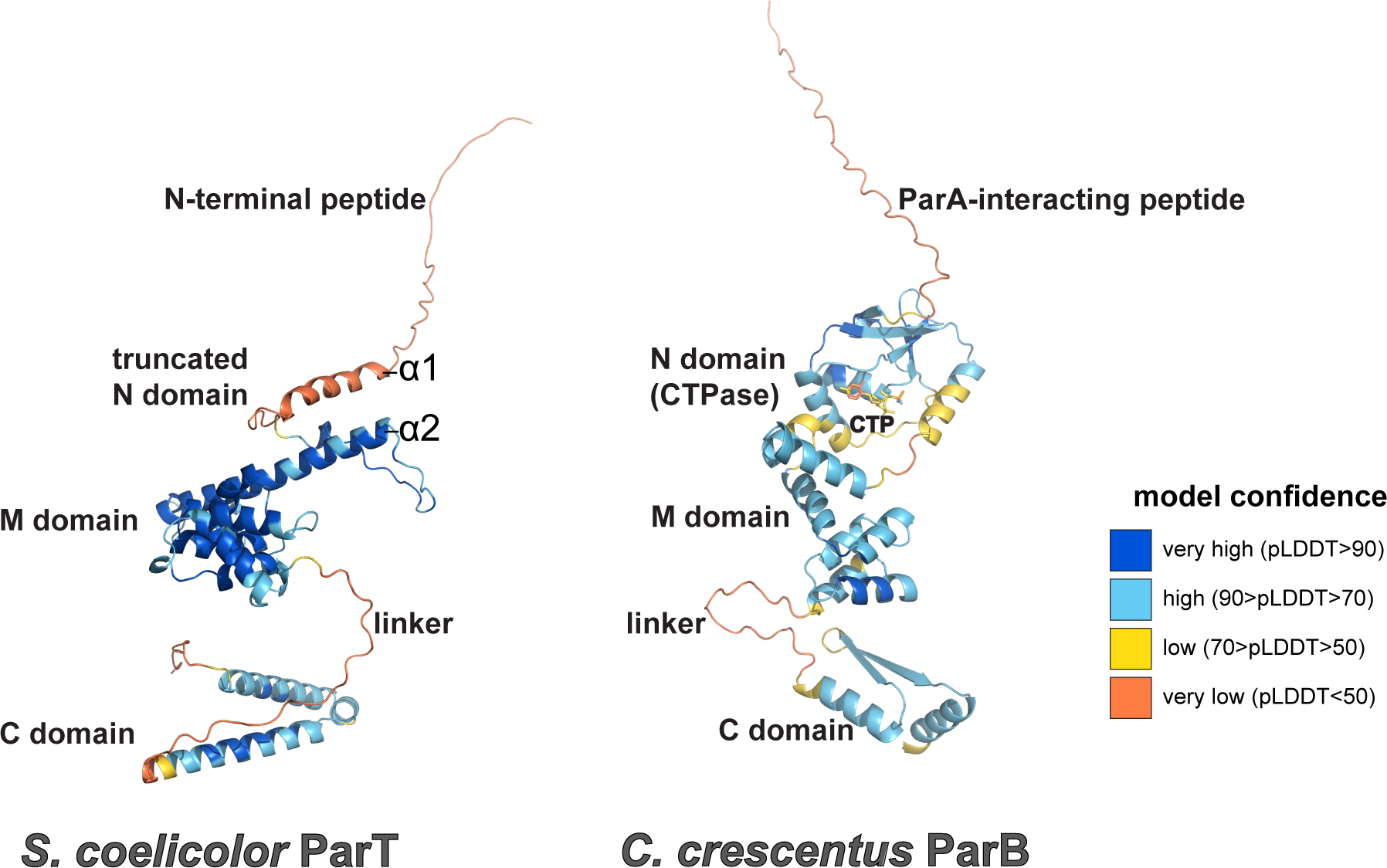
AlphaFold2-predicted structures of *S. coelicolor* ParT and *C. crescentus* chromosomal ParB. Structures were colored according to the model confidence score (pLDDT).

**Fig. S2.**
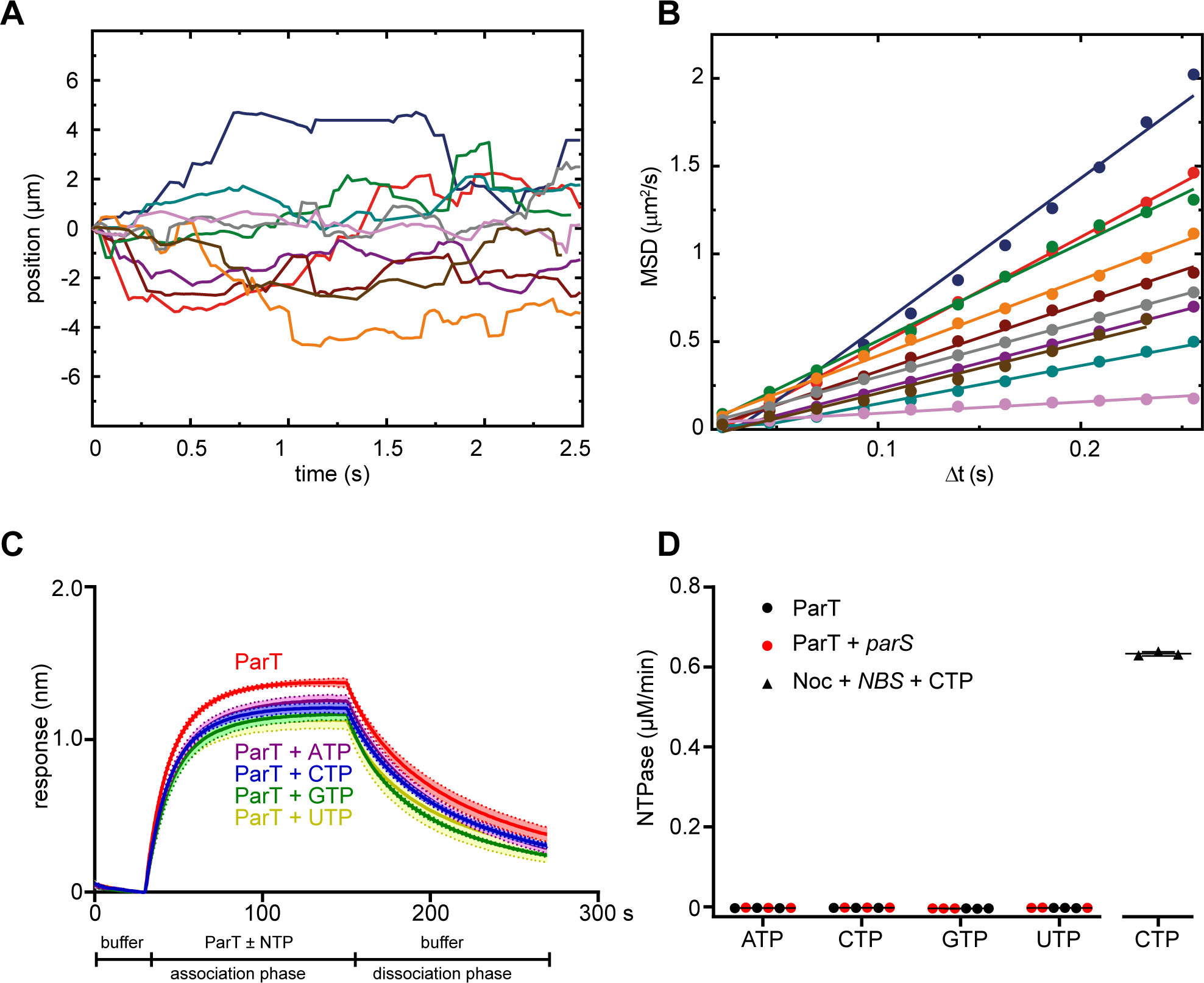
ParT diffuses on *parS* DNA independently of NTP. (**A**) Representative AF488-HALO-ParT trajectories were measured on the DNA (n=154). **(B)** Mean squared displacement (MSD) of AF488-HALO-ParT trajectories for different time intervals (Δt). The diffusion constant of ParT (1.11 ± 0.05 μm^2^/s, mean ± SEM) was calculated as half of the slope of the linear fit of MSD versus Δt. **(C)** Accumulation of ParT on a 180-bp *parS* closed DNA loop did not change significantly in the presence of NTP. BLI analysis of the interaction between a premix of 1 µM ParT ± 1 mM NTP with a 180-bp dual biotin-labeled DNA that contains either a *parS* or a scrambled *parS* site. Mean and standard deviation (shading) from three replicates are shown. **(D)** ParT does not have detectable NTPase activity compared to a ParB-like CTPase positive control Noc (55). NTPase activities were measured at 1 μM of ParT/Noc and 1mM NTP, in the presence or absence of 1 μM *parS* DNA or Noc-binding site (*NBS*) DNA. Experiments were triplicated.

**Fig. S3.**
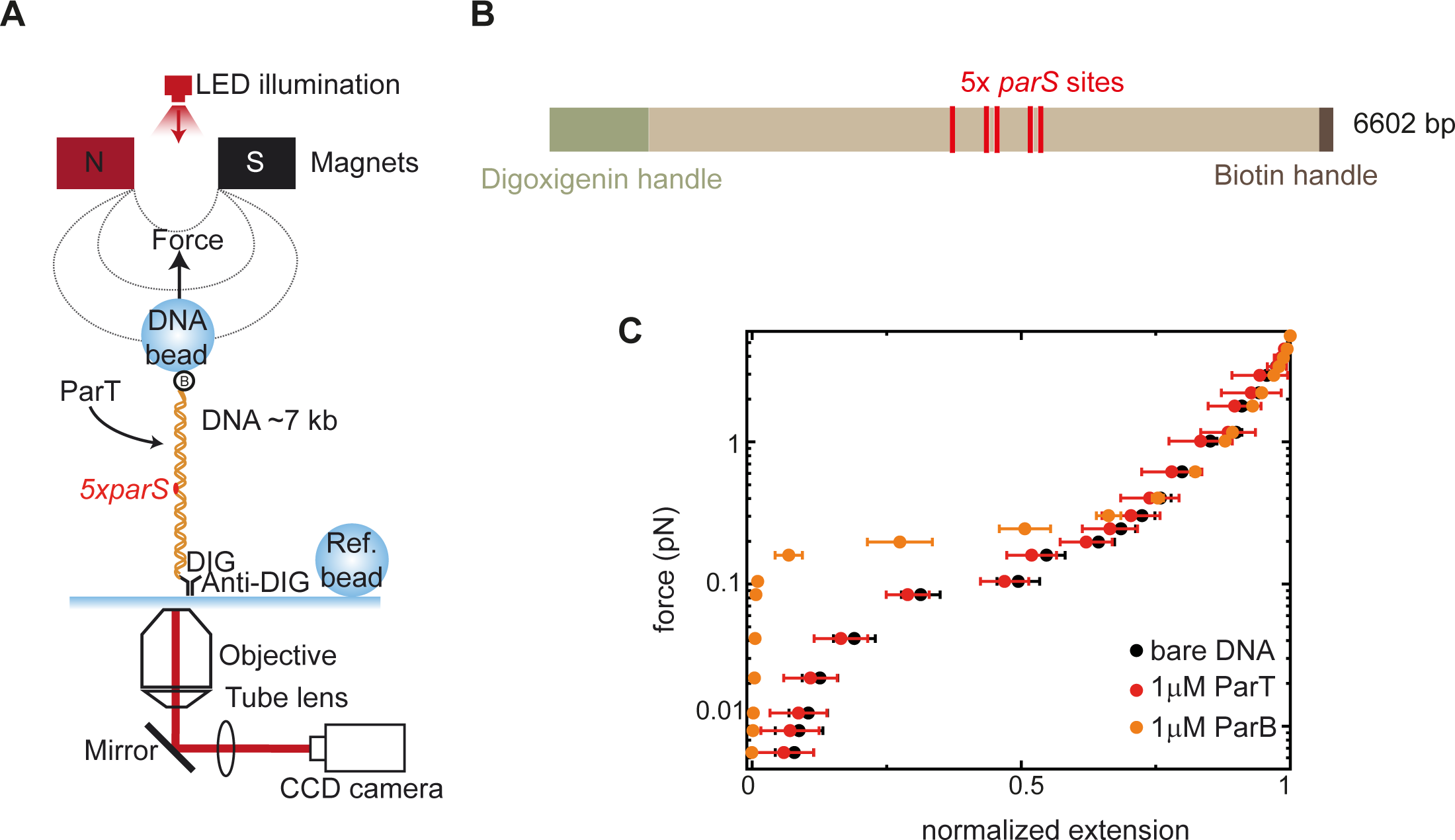
ParT did not condense a DNA containing 5x *parS* sites. (**A**) Schematic diagram of the basic magnetic tweezers (MT) components and the layout of the experiment. **(B)** A schematic representation of a 5x *parS* DNA, the position of the *parS* sites is represented to scale. **(C)** Average force-extension curves (mean ± SEM) of bare 5x *parS* DNA molecules (n = 29) and in the presence of 1μM ParT + 2 mM CTP (n = 29) or 1μM *Bacillus subtilis* ParB + 2mM CTP (n= 11).

**Fig. S4.**
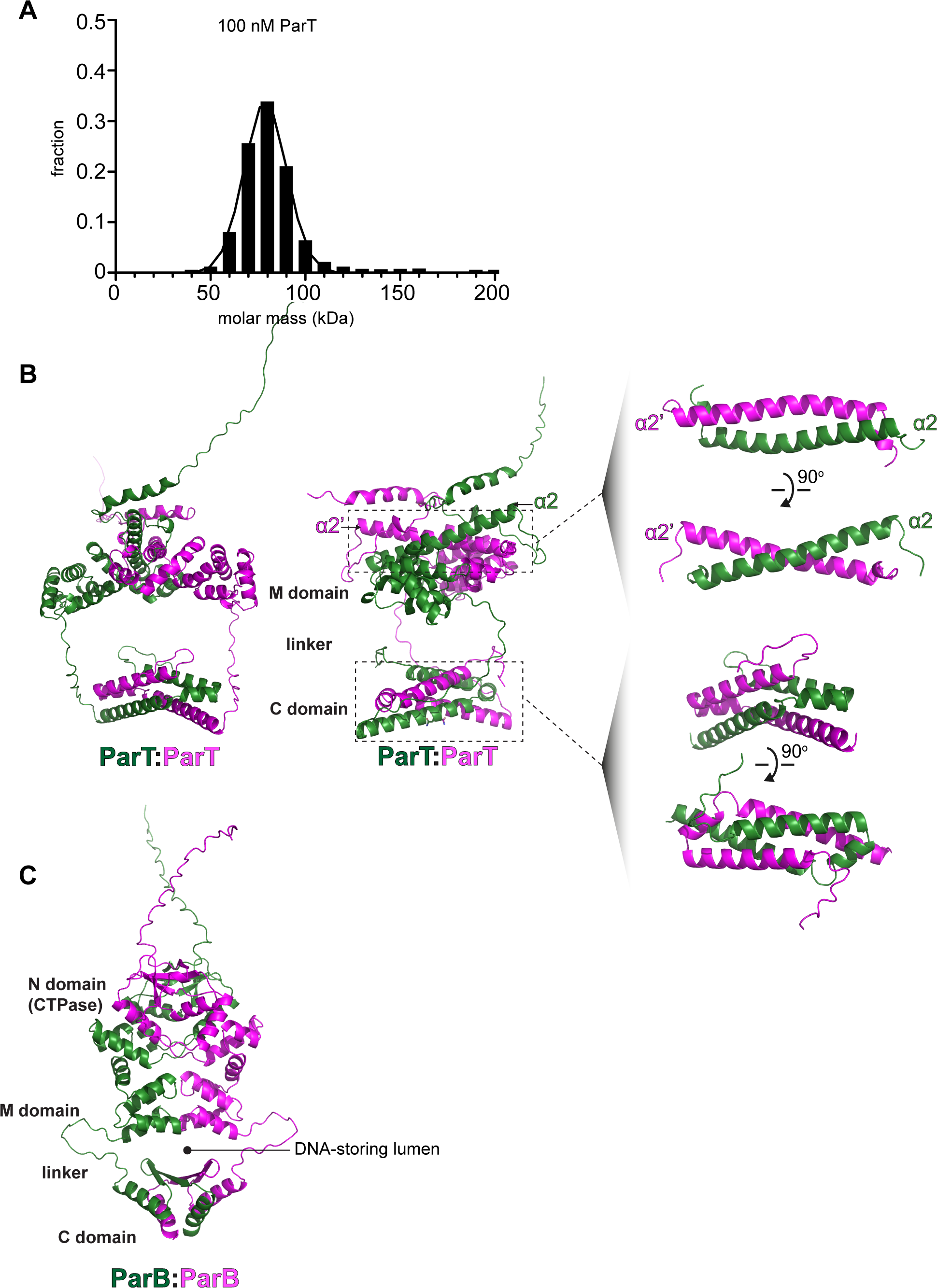
ParT can form a dimer in solution, as indicated by mass photometry. (**A**) Mass photometry measurements were carried out at room temperature at 100 nM ParT. The mass distribution curve was merged from triplicated mass-photometry experiments. **(B)** An AlphaFold2 Multimer-predicted structure of a ParT dimer and its dimerization interfaces at the N domain and C domain. **(C)** An AlphaFold2 Multimer-predicted structure of *C. crescentus* ParB dimer that shows the CTPase N domain, the *parS*-binding M domain, the dimerization C domain, and a DNA-storing lumen that is constituted by a flexible linker that connects M and C domains. During ParB sliding, DNA is entrapped within this DNA-storing lumen (22, 33).

**Fig. S5.**
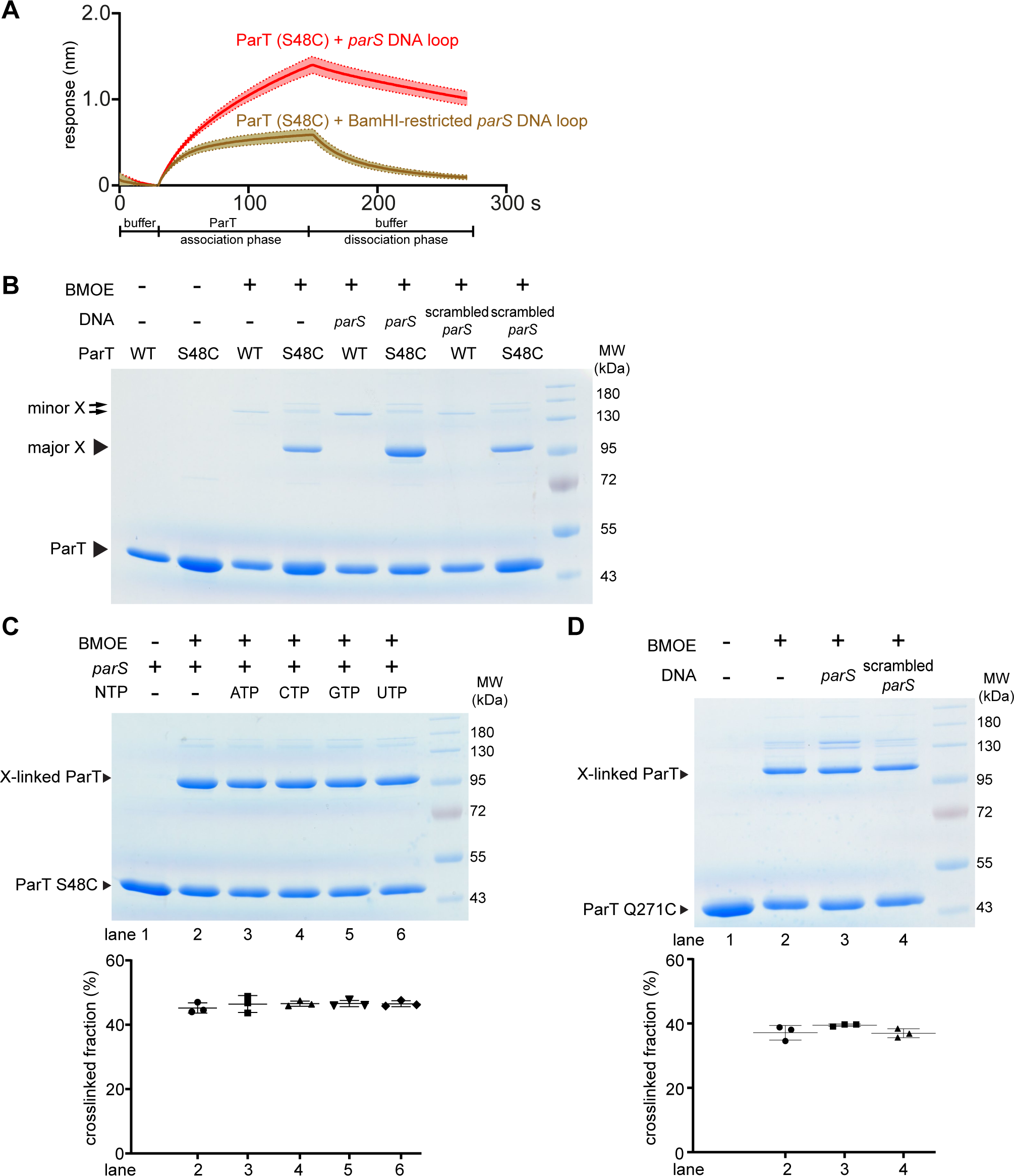
The crosslinking of ParT (S48C) is not affected by NTP. (**A**) BLI analysis of the interaction between 1 µM ParT (S48C) with a 180-bp dual biotin-labeled *parS* closed DNA loop (red) or a BamHI-restricted loop (brown) to generate a free DNA end. BLI analysis indicated that ParT (S48C) could accumulate on a closed DNA loop but the accumulation was much reduced on a DNA with an open end. Mean and standard deviation (shading) from three replicates are shown. **(B)** SDS-PAGE analysis of BMOE crosslinking products of ParT (WT) and ParT (S48C) with or without 40-bp *parS* DNA duplex or scrambled *parS* DNA duplex. ParT (WT) with a native cysteine residue C81 crosslinked minimally in tested conditions (double arrows, minor X species). Most crosslinked products (single arrow, major X) appeared when a ParT (S48C) variant was employed instead. **(C)** SDS-PAGE analysis of BMOE crosslinking products of 4 µM ParT (S48C) in the presence of 4 µM 40-bp *parS* DNA and 1 mM NTP. Experiments were triplicated, and a quantification (mean ± SD) of the major crosslinked fraction is shown below the representative image. **(D)** SDS-PAGE analysis of BMOE crosslinking products of 4 µM ParT (Q271C) in the presence of 4 µM 40-bp *parS* DNA or scrambled *parS* DNA. Experiments were triplicated, and a quantification (mean ± SD) of the major crosslinked fraction is shown below the representative image.

