## Supplementary Tables for "Assembly of a segrosome by a CTP-independent ParB-like protein"

**Supplementary Table 1. DNA oligonucleotides used in this study**

| **Name** | **Sequence** |
| --- | --- |
| *parS_Fw_bio* | 5’BIO/GGCCGACCTCGTGTCTCCAATTGGAGACATCAACGAGGGC |
| *parS_Rv* | GCCCTCGTTGATGTCTCCAATTGGAGACACGAGGTCGGCC |
| *scram parS_Fw* | 5’BIO/ATCGGGCACCTTAGGGCAATCGTAAGTTCGGCGCACGACC |
| *scram parS_Rv* | GGTCGTGCGCCGAACTTACGATTGCCCTAAGGTGCCCGAT |
| *M13F* | 5’BIO/CGCCAGGGTTTTCCCAGTCACGAC |
| *M13R* | 5’BIO/AGGAAACAGCTATGACCAT |
| *ParT_CFLAG_Fw* | GATCGTCTAGAACAGGAGGCCCCATATGAGCCGCCGCTCCCTCGCCCTC |
| *ParT_CFLAG_Rv* | TCCGCTCATGAGAACCTAGGATCCATTACTTGTCGTCATCGTCCTTGTAGTCGCCCGCGTCGTCGGCCACCTGCTCACCG |
| *ParT_Fw* | GTGAGCCGCCGCTCCCTCGCCCTCCCGTCG |
| *ParT_Rv* | TTACGCGTCGTCGGCCACCTGCTCACCGGAG |

| Annealed oligonucleotides with 1x *parS* site | *342.R-1xparS-ParT* | GCGTCGCCCTCGTTGATGTCTCCAATTGGAGACACGAGGTCGGCCGGTAC |
| --- | --- | --- |
|  | *347.F-1xparS-ParT ctrap* | CGGCCGACCTCGTGTCTCCAATTGGAGACATCAACGAGGGCGACGCGTAC |
| Annealed oligonucleotides with 2x *parS* site | *343.F-2parS dianas* | TGAGCTCGAGGCGCCCGTGTCTCCAATTGGAGACATCAAGTATAGAGCACCTGTTACGTACCTAGTCAAGGACTCAGCGTGTCTCCAATTGGAGACATCGTTAACGTCGACGC |
|  | *344.R-2parS dianas* | TCAGCGTCGACGTTAACGATGTCTCCAATTGGAGACACGCTGAGTCCTTGACTAGGTACGTAACAGGTGCTCTATACTTGATGTCTCCAATTGGAGACACGGGCGCCTCGAGC |
| MT 1x *parS* PCR fragment | *329.PCR_template_T7pCDNA3_FW* | ATCGAAATTAATACGACTCACTATAGG |
|  | *325.R req lambda 4* | GCCCGGATTCAAATGCTGCAG |
| MT DIG handle | *57.FMH_F2_BamHI-ApaI* | GCGTAAGTGGATCCGGGCCCCGACTCACTATAGGGAGAC  CGGC |
|  | *JOE_R1* | AGTAAGCGCCGTCAGACCAG |
| MT BIO handle | *57.FMH_F2_BamHI-ApaI* | GCGTAAGTGGATCCGGGCCCCGACTCACTATAGGGAGAC  CGGC |
|  | *209.BsrGI 71short handle* | CGATAACCAACTGGCGATG |
| C-trap BIO handle | *57.FMH_F2_BamHI-ApaI* | GCGTAAGTGGATCCGGGCCCCGACTCACTATAGGGAGACCGGC |
|  | *JOE_R1* | AGTAAGCGCCGTCAGACCAG |

**Supplementary Table 2. Plasmids used in this study**

| **Vector** | **Description** | **Use** | **Source** |
| --- | --- | --- | --- |
| *pET21b* | Empty expression vector for use in *E .coli* | Cloning | Lab stock |
| *pIJ10257* | Plasmid cloning vector for the conjugal transfer of DNA (under control of the *ermE** constitutive promoter) from *E. coli* to *Streptomyces* spp. Integrates specifically at the ΦBT1 phage integration site (HygromycinR) | Cloning | (1) |
| *pUZ8002* | A helper plasmid for conjugative transfer of pIJ10257 from *E. coli* to *S. coelicolor* | Conjugation to transfer DNA from *E. coli* to *S. coelicolor* | (2) |
| *pET21b::parT-(his)_6_* | Over-expression vector of ParT WT, 6xHis-tagged at C‑terminus | Protein over-expression | This study |
| *pET21b::parA-(his)_6_* | Over-expression vector of ParA WT, 6xHis-tagged at C‑terminus | Protein over-expression | This study |
| *pET21b::parT S48C-(his)_6_* | Over-expression vector of ParT S48C, 6xHis-tagged at C‑terminus | Protein over-expression | This study |
| *pET21b::parT Q271C-(his)_6_* | Over-expression vector of ParT Q271C, 6xHis-tagged at C‑terminus | Protein over-expression | This study |
| *pIJ10257::parT-flag* | Integration vector haboring a *parT* variant with a FLAG tag at C-terminus | Conjugation to transfer *pIJ10257::parT-flag* from *E. coli* to *S. coelicolor* | This study |
| *pIJ10257::parT* | Integration vector harboring a WT *parT* | Conjugation to transfer *pIJ10257::parT* from *E. coli* to *S. coelicolor* | This study |

**Supplementary Table 3. Strains used in this study**

| **Strain** | **Use** | **Source** |
| --- | --- | --- |
| *E. coli DH5α* | Cloning strain for the plasmids constructed in this study | Lab stock |
| *E. coli Rosetta (DE3) pLys* | Host strains for protein over-expression plasmids | Lab stock |
| *S. coelicolor A3(2)* | WT *S. coelicolor* A3(2) used as a recipient for *parT* variants, SCP1^+^ SCP2^+^ | Lab stock |
| *E. coli Rosetta (DE3) pLys pET21b::parT-(his)_6_* | Over-expression of ParT WT 6xHis | This study |
| *E. coli Rosetta (DE3) pLys pET21b::parA-(his)_6_* | Over-expression of ParA WT 6xHis | This study |
| *E. coli Rosetta (DE3) pLys pET21b::parT S48C-(his)_6_* | Over-expression of ParT S48C 6xHis | This study |
| *E. coli Rosetta (DE3) pLys pET21b::parT Q271C-(his)_6_* | Over-expression of ParT Q271C 6xHis | This study |
| *E. coli ET12567* + *pUZ8002* | A conjugative *E. coli* strain carrying a helper plasmid pUZ8002 | (2) |
| *E. coli ET12567 pIJ10257::parT-flag* + *pUZ8002* | Conjugative transfer from *E. coli* to *S. coelicolor* A3(2) to introduce a FLAG-tagged *parT* variant at the ΦBT1 phage integration site | This study |
| *E. coli ET12567 pIJ10257::parT* + *pUZ8002* | Conjugative transfer from *E. coli* to *S. coelicolor* A3(2) to introduce a WT *parT* variant at the ΦBT1 phage integration site | This study |
| *S. coelicolor A3(2)* ΦBT1*::parT-flag* | ChIP-seq experiment using an anti-FLAG antibody | This study |
| *S. coelicolor A3(2)* ΦBT1*::parT* | ChIP-seq experiment (negative control) using an anti-FLAG antibody | This study |

**Supplementary Table 4.** List of ChIP-seq data used in this study

| **ChIP-seq samples** | **Genetic backgrounds** | **antibody** | **replicates** | **numbers of mapped reads** | **Source** |
| --- | --- | --- | --- | --- | --- |
| sample 1 | *S. coelicolor A3(2)* ΦBT1*::parT* | anti-FLAG | replicate 1 | 3776638 | This study |
| sample 2 | *S. coelicolor A3(2)* ΦBT1*::parT-flag* | anti-FLAG | replicate 2 | 5429409 | This study |
| sample 4 | *S. coelicolor A3(2)* ΦBT1*::parT* | anti-FLAG | replicate 1 | 4838459 | This study |
| sample 5 | *S. coelicolor A3(2)* ΦBT1*::parT-flag* | anti-FLAG | replicate 2 | 4718368 | This study |

**Supplementary Table 5. dsDNA constructs used in this work for MT and C-trap experiments.**

*parS* sites in direct or inverted orientation are highlighted in green.

| **Fragment** | **Sequence** |
| --- | --- |
| MT central fragment with 5x *parS* sites  (6602 bp) | GGCCGCCCGCCATCGGGATTTTCACCACATCAATGGGGTAACGGTTTTTCCCAGCCACACGCTGCATGACATGCCACCGGCCATTTTTCAGTTGCTGAATAAACGCGCCGGGAATACGACGGTTACCCACCACAAGCACGCTGCCGCCACCTTTCAGGGATGAACGCTGCCCCTTTTTACGACGCCTGCGGCGCGAAAGGACAACCCGCGCATTACCCAGCTTGATTACGGGCAAATCCCCCCGGTTAACTTTGATTCTGGCCTGCGGATTTTTGACCGTGGCCCTTTTCAGCCTGGCCCTTTCCTTTACCAGTTTCCGGCGTACCTTTGTCTCACGGGCAACCTGTGACGCCGACTGCGATATCGCGGATGAAGCAACGCGGTTAATGGCCATTGCGGCGGCACCAGGCACCGCCGTTTTGCTGATACGGCTGAGGTTTTCAACGGCCTGCTCAAGACCTTTTATGGCCATACATCCCCCTTTCAGCGGCGACGGTTAACGGCAGGCGGTACGCCCCGTCCAAGCCAGAGATGACAACTTCCGCCATCATCCGGCGAAACCCGATCTACCCAGAAATTTTCCTCACCGATGGTCAGCGTGTCTCCACGCCGCAGCTGCCGCACCTCATCAGTCCGGACAAACAGGGACGGGCTGGAGCCTTCAACGCGCACGCCCTGTCCGGCATAGCTGATATTTTCAGGGTCATCAAAAACACCACGTATCACCGCACCTGACTGCTCACCGGATGTAATGGTGGCTGACGTTCCCATGTACCCGCGTATCGTTTCATCGGCGCGGGCAATGGCAGCATCGAACAGGTTATCGAAATCAGCCACAGCGCCTCCCGTTATTGCATTCTGGCCAGGCCGCGCTCTGTCATTTCGGCTGCCACACCGGCAGAGACACGAAACGCCGTTCCCGGCAGCACAAATGCCACAGGTTCATCCCGCGTGGCGTGAAGTGCATCAGTATGCAGCTTCACCAGTGCCACGACCGTGACCAGTTCAGACGTATCCAGAATCACGGTATCCGGCTGCGCTGATCCCACCTCATTTTCATGTCCGGTCAGCACATTTTCCCGGCTGAGAGGGGTGTCCTGACCGGCAGTTTCATCCGTGTCATCAAGCTCCTCTTTCAGCTCTGCCACACGGAGCGCCAGTTCTTCTTTCGTCCCCGTCAGGCTGACATCACGGTTCAGTTGTTCACCCAGCGAGCGGAGACGGGCAATCAGTTCATCTTTCGTCATGGACTCCTCCACAGAGAAACAATGGCCCCGAAGGGCCATGATTACGCCAGTTGTACGGACACGAACTCATCAGGGTCAGCCAGCAGCATCAGCGGTGCTGACTGAATCATGGTGAACTCACGCGCCGGATCGCCGGTGGTCACCCAGTTTTTCGGGTAACGGGCAGAGGCGTTAATGCCTTCGCGCTGTGCGTCCGCATCCTGAATGCAGCCATAGGTGCGCAGACCGCGTGCCTGAGTGTTCCCCAGCACCATCGTGTTGTCCGGCAGGAAGTTCTTTTTGACGCCGTTTTCCACGTACTGTCCGGAATACACGACGATGGCCACATCGCCATACATCCCCTTATAGGACACCGCTTTGCCCAGGTCTTTCACCGCTGTCTCCAGCTCGGAATTAGAGCCACGACGGGTATCCAGCTTCTCCTTGACGGCTTTGAAGGAACGGAACAGCGCCCAGCCTTTCGGATCGAACACGATGATATTCACCACACCGCTGGCGTTCAGCGCGTAGGCTTCGATATCGTCGGTCGGGTCATACGTGGACTTGTCACGCTTGCTCCACTCCGTGCCGCCGGACTGCGTGATGTTATTCTCCTCACTGCGGCCCATATCCACCTCAACCGGATCGAAGGCTTCACCGGTCATGGTGTATTTGCCCTTAAGCACGGCAGAAACTGCCTGCATCTCTTCGACCTGAGCAATGGCCAGCTCTTCGTCACGCATGTTCTGCATGATGATGCGACGGCGGCGGTAAGCCGGGTCCGCCAGATTCTGCGGATCTTCATCCGGCAGGCGACGCAGGGTCATCTGCGGATTCACTTCATGCTTCGGCTTGACATATCCCGGCGTAAATTCAGAGGTGGAGCCGCCACGGGAACGGATAACCTCACCGGAAACAATCGGCGAAACGTACAGCGCCATGTTTACCAGTCCCGGAATTTGTGAGAGATAGACTTTCTCCGTGGTGAAGGGATAGCTCTCACGGAAAAAGAGACGCAGAAACAGCGGATCAAACTTAAATTTCTGCTCATTTGCCGCCAGCAGTTGGGCGGTTGTGTACATCGACATAAAAAAATCCCGTAAAAAAAGCCGCACAGGCGGCCTTTAGTGATGAAGGGTAAAGTTAAACGATGCTGATTGCCGTTCCGGCAAACGCGGTCCGTTTTTTCGTCTCGTCGCTGGCAGCCTCCGGCCAGAGCACATCCTCATAACGGAACGTGCCGGACTTGTAGAACGTCAGCGTGGTGCTGGTCTGGTCAGCAGCAACCGCAAGAATGCCAACGGCAGCACCGTCGGTGGTGCCATCCCACGCAACCAGCTTACGGCTGGAGGTGTCCAGCATCAGCGGGGTCATTGCAGGCGCTTTCGCACTCAATCCGCCGGGCGCGGTTGCGGTATGAGCCGGGTCACTGTTGCCCTGCGGCTGGTAATGGGTAAAGGTTTCTTTGCTCGTCATAAACATCCCTTACACTGGTGTGTTCAGCAAATCGTTAACGGCATCAGATGCCGGGTTACCTGCAGCCAGCGGTGCCGGTGCCCCCTGCATCAGACGATCCAGCGCAGTGTCACTGCGCGCCTGTGCACTCTGTGGTGCTGCGGCCAGAATGCGGCGGGCCGTTTTCACGGTCATACCGGGGGTTTCTGCCAGCACGCGTGCCTGTTCTTCGCGTCCGTGAGCCTCCTCACAGTTGAGAATTCGTCGTAGAATTCAACGTGGAATTCCTATCGGAATTCTCGGATGAATTCGGATCCTTATGATTCTCGTTCAGGGTACCGGCCGACCTCGTGTCTCCAATTGGAGACATCAACGAGGGCGACGCGTACCAGTTCAGGAAGCGGTGATGCTGATAGAAGCCGGACTGAGTACCTACGAGAAAGAGTGCGCAAAACGCGGTGACGACTATCAGGAAATTTTTGCCCAGCAGGTCCGTGAAACGATGGAGCGCCGTGCAGCCGGTCTTAAACCGCCCGCCTGGGCGGCTGCAGCATTTGAATCCGGGCTCGTCATGCCCTCGTGGTCAGGTCTGGACGACGAGCCGTTCGATCCTGCCACGTCGCCCGTTACACCGGACCTTGGAGTTGTCTCTGACACATTCTGGCAACGATGTCTCCAATTGGAGACACGCTGAGTCCTTGACTAGGTACGTAACAGGTGCTCTATACTTGATGTCTCCAATTGGAGACACGGGCGCCTGCCAAATGTAAAGCGCAGCGCCCATCCATTTGCCTTTGCGGCAGCGGGGCCACAGGCAGAGCAGATCATCTCTGATCCATTGCCCCTGCCACCTCACTCGCCTGCAAGCCCGGTCGCCCGTGTCCATGAACTCGATGGGCAGGTACTTCTCCTCGGCGTGGGACACGATGCCAACACGACGCTGCATCTTGCCGAGTTGATGGCAAAGGTTCCCTATGGGGTGCCGAGACACTGCACCATTCTTCAGGATGGCAAGTTGGTACGCGTCGATTATCTCGACGTTAACGATGTCTCCAATTGGAGACACGCTGAGTCCTTGACTAGGTACGTAACAGGTGCTCTATACTTGATGTCTCCAATTGGAGACACGGGCGCCTCGAGCTCGAGAATGACCACTGCTGTGAGCGCTTTGCCTTGGCGGACAGGTGGCTCAAGGAGAAGAGCCTTCAGAAGGAAGGTCCAGTCGGTCATGCCTTTGCTCGGTTGATCCGCTCCCGCGACATTGTGGCGACAGCCCTGGGTCAACTGGGCCGAGATCCGTTGATCTTCCTGCATCCGCCAGAGGCGGGATGCGAAGAATGCGATGCCGCTCGCCAGTCGATTGGCTGAGCTCATAAGTGAGGAAGGCCGGGCGGGAAACTGCCCGGCCTGAACATACCTGAATGGTTATCCCCGCTGACGCGGGGAACATAAGTTCGGGGATCCTCTAGAGTCGACCTGCAGGCATGCAAGCTTGGCGTAATCATGGTCATAGCTGTTTCCTGTGTGAAATTGTTATCCGCTCACAATTCCACACAACATACGAGCCGGAAGCATAAAGTGTAAAGCCTGGGGTGCCTAATGAGTGAGCTAACTCACATTAATTGCGTTGCGCTCACTGCCCGCTTTCCAGTCGGGAAACCTGTCGTGCCAGCTGCATTAATGAATCGGCCAACGCGCGGGGAGAGGCGGTTTGCGTATTGGGCGCTCTTCCGCTTCCTCGCTCACTGACTCGCTGCGCTCGGTCGTTCGGCTGCGGCGAGCGGTATCAGCTCACTCAAAGGCGGTAATACGGTTATCCACAGAATCAGGGGATAACGCAGGAAAGAACATGTGAGCAAAAGGCCAGCAAAAGGCCAGGAACCGTAAAAAGGCCGCGTTGCTGGCGTTTTTCCATAGGCTCCGCCCCCCTGACGAGCATCACAAAAATCGACGCTCAAGTCAGAGGTGGCGAAACCCGACAGGACTATAAAGATACCAGGCGTTTCCCCCTGGAAGCTCCCTCGTGCGCTCTCCTGTTCCGACCCTGCCGCTTACCGGATACCTGTCCGCCTTTCTCCCTTCGGGAAGCGTGGCGCTTTCTCATAGCTCACGCTGTAGGTATCTCAGTTCGGTGTAGGTCGTTCGCTCCAAGCTGGGCTGTGTGCACGAACCCCCCGTTCAGCCCGACCGCTGCGCCTTATCCGGTAACTATCGTCTTGAGTCCAACCCGGTAAGACACGACTTATCGCCACTGGCAGCAGCCACTGGTAACAGGATTAGCAGAGCGAGGTATGTAGGCGGTGCTACAGAGTTCTTGAAGTGGTGGCCTAACTACGGCTACACTAGAAGGACAGTATTTGGTATCTGCGCTCTGCTGAAGCCAGTTACCTTCGGAAAAAGAGTTGGTAGCTCTTGATCCGGCAAACAAACCACCGCTGGTAGCGGTGGTTTTTTTGTTTGCAAGCAGCAGATTACGCGCAGAAAAAAAGGATCTCAAGAAGATCCTTTGATCTTTTCTACGGGGTCTGACGCTCAGTGGAACGAAAACTCACGTTAAGGGATTTTGGTCATGAGATTATCAAAAAGGATCTTCACCTAGATCCTTTTAAATTAAAAATGAAGTTTTAAATCAATCTAAAGTATATATGAGTAAACTTGGTCTGACAGTTACCAATGCTTAATCAGTGAGGCACCTATCTCAGCGATCTGTCTATTTCGTTCATCCATAGTTGCCTGACTCCCCGTCGTGTAGATAACTACGATACGGGAGGGCTTACCATCTGGCCCCAGTGCTGCAATGATACCGCGAGACCCACGCTCACCGGCTCCAGATTTATCAGCAATAAACCAGCCAGCCGGAAGGGCCGAGCGCAGAAGTGGTCCTGCAACTTTATCCGCCTCCATCCAGTCTATTAATTGTTGCCGGGAAGCTAGAGTAAGTAGTTCGCCAGTTAATAGTTTGCGCAACGTTGTTGCCATTGCTACAGGCATCGTGGTGTCACGCTCGTCGTTTGGTATGGCTTCATTCAGCTCCGGTTCCCAACGATCAAGGCGAGTTACATGATCCCCCATGTTGTGCAAAAAAGCGGTTAGCTCCTTCGGTCCTCCGATCGTTGTCAGAAGTAAGTTGGCCGCAGTGTTATCACTCATGGTTATGGCAGCACTGCATAATTCTCTTACTGTCATGCCATCCGTAAGATGCTTTTCTGTGACTGGTGAGTACTCAACCAAGTCATTCTGAGAATAGTGTATGCGGCGACCGAGTTGCTCTTGCCCGGCGTCAATACGGGATAATACCGCGCCACATAGCAGAACTTTAAAAGTGCTCATCATTGGAAAACGTTCTTCGGGGCGAAAACTCTCAAGGATCTTACCGCTGTTGAGATCCAGTTCGATGTAACCCACTCGTGCACCCAACTGATCTTCAGCATCTTTTACTTTCACCAGCGTTTCTGGGTGAGCAAAAACAGGAAGGCAAAATGCCGCAAAAAAGGGAATAAGGGCGACACGGAAATGTTGAATACTCATACTCTTCCTTTTTCAATATTATTGAAGCATTTATCAGGGTTATTGTCTCATGAGCGGATACATATTTGAATGTATTTAGAAAAATAAACAAATAGGGGTTCCGCGCACATTTCCCCGAAAAGTGCCACCTGACGTCTAAGAAACCATTATTATCATGACATTAACCTATAAAAATAGGCGTATCACGAGGCCCTTTCGTCTCGCGCGTTTCGGTGATGACGGTGAAAACCTCTGACACATGCAGCTCCCGGAGACGGTCACAGCTTGTCTGTAAGCGGATGCCGGGAGCAGACAAGCCCGTCAGGGCGCGTCAGCGGGTGTTGGCGGGTGTCGGGGCTGGCTTAACTATGCGGCATCAGAGCAGATTGTACTGAGAGTGCACCATATGGGGCC |
| C-trap central fragment with 3x *parS* sites  (20622 bp) | GGCCGCGGTGTGCTCCTTATTTATACATAACGAAAAACGCCTCGAGTGAAGCGTTATTGGTATGCGGTAAAACCGCACTCAGGCGGCCTTGATAGTCATATCATCTGAATCAAATATTCCTGATGTATCGATATCGGTAATTCTTATTCCTTCGCTACCATCCATTGGAGGCCATCCTTCCTGACCATTTCCATCATTCCAGTCGAACTCACACACAACACCATATGCATTTAAGTCGCTTGAAATTGCTATAAGCAGAGCATGTTGCGCCAGCATGATTAATACAGCATTTAATACAGAGCCGTGTTTATTGAGTCGGTATTCAGAGTCTGACCAGAAATTATTAATCTGGTGAAGTTTTTCCTCTGTCATTACGTCATGGTCGATTTCAATTTCTATTGATGCTTTCCAGTCGTAATCAATGATGTATTTTTTGATGTTTGACATCTGTTCATATCCTCACAGATAAAAAATCGCCCTCACACTGGAGGGCAAAGAAGATTTCCAATAATCAGAACAAGTCGGCTCCTGTTTAGTTACGAGCGACATTGCTCCGTGTATTCACTCGTTGGAATGAATACACAGTGCAGTGTTTATTCTGTTATTTATGCCAAAAATAAAGGCCACTATCAGGCAGCTTTGTTGTTCTGTTTACCAAGTTCTCTGGCAATCATTGCCGTCGTTCGTATTGCCCATTTATCGACATATTTCCCATCTTCCATTACAGGAAACATTTCTTCAGGCTTAACCATGCATTCCGATTGCAGCTTGCATCCATTGCATCGCTTGAATTGTCCACACCATTGATTTTTATCAATAGTCGTAGTCATACGGATAGTCCTGGTATTGTTCCATCACATCCTGAGGATGCTCTTCGAACTCTTCAAATTCTTCTTCCATATATCACCTTAAATAGTGGATTGCGGTAGTAAAGATTGTGCCTGTCTTTTAACCACATCAGGCTCGGTGGTTCTCGTGTACCCCTACAGCGAGAAATCGGATAAACTATTACAACCCCTACAGTTTGATGAGTATAGAAATGGATCCACTCGTTATTCTCGGACGAGTGTTCAGTAATGAACCTCTGGAGAGAACCATGTATATGATCGTTATCTGGGTTGGACTTCTGCTTTTAAGCCCAGATAACTGGCCTGAATATGTTAATGAGAGAATCGGTATTCCTCATGTGTGGCATGTTTTCGTCTTTGCTCTTGCATTTTCGCTAGCAATTAATGTGCATCGATTATCAGCTATTGCCAGCGCCAGATATAAGCGATTTAAGCTAAGAAAACGCATTAAGATGCAAAACGATAAAGTGCGATCAGTAATTCAAAACCTTACAGAAGAGCAATCTATGGTTTTGTGCGCAGCCCTTAATGAAGGCAGGAAGTATGTGGTTACATCAAAACAATTCCCATACATTAGTGAGTTGATTGAGCTTGGTGTGTTGAACAAAACTTTTTCCCGATGGAATGGAAAGCATATATTATTCCCTATTGAGGATATTTACTGGACTGAATTAGTTGCCAGCTATGATCCATATAATATTGAGATAAAGCCAAGGCCAATATCTAAGTAACTAGATAAGAGGAATCGATTTTCCCTTAATTTTCTGGCGTCCACTGCATGTTATGCCGCGTTCGCCAGGCTTGCTGTACCATGTGCGCTGATTCTTGCGCTCAATACGTTGCAGGTTGCTTTCAATCTGTTTGTGGTATTCAGCCAGCACTGTAAGGTCTATCGGATTTAGTGCGCTTTCTACTCGTGATTTCGGTTTGCGATTCAGCGAGAGAATAGGGCGGTTAACTGGTTTTGCGCTTACCCCAACCAACAGGGGATTTGCTGCTTTCCATTGAGCCTGTTTCTCTGCGCGACGTTCGCGGCGGCGTGTTTGTGCATCCATCTGGATTCTCCTGTCAGTTAGCTTTGGTGGTGTGTGGCAGTTGTAGTCCTGAACGAAAACCCCCCGCGATTGGCACATTGGCAGCTAATCCGGAATCGCACTTACGGCCAATGCTTCGTTTCGTATCACACACCCCAAAGCCTTCTGCTTTGAATGCTGCCCTTCTTCAGGGCTTAATTTTTAAGAGCGTCACCTTCATGGTGGTCAGTGCGTCCTGCTGATGTGCTCAGTATCACCGCCAGTGGTATTTATGTCAACACCGCCAGAGATAATTTATCACCGCAGATGGTTATCTGTATGTTTTTTATATGAATTTATTTTTTGCAGGGGGGCATTGTTTGGTAGGTGAGAGATCTGAATTGCTATGTTTAGTGAGTTGTATCTATTTATTTTTCAATAAATACAATTGGTTATGTGTTTTGGGGGCGATCGTGAGGCAAAGAAAACCCGGCGCTGAGGCCGGGTTATTCTTGTTCTCTGGTCAAATTATATAGTTGGAAAACAAGGATGCATATATGAATGAACGATGCAGAGGCAATGCCGATGGCGATAGTGGGTATCATGTAGCCGCTTATGCTGGAAAGAAGCAATAACCCGCAGAAAAACAAAGCTCCAAGCTCAACAAAACTAAGGGCATAGACAATAACTACCGATGTCATATACCCATACTCTCTAATCTTGGCCAGTCGGCGCGTTCTGCTTCCGATTAGAAACGTCAAGGCAGCAATCAGGATTGCAATCATGGTTCCTGCATATGATGACAATGTCGCCCCAAGACCATCTCTATGAGCTGAAAAAGAAACACCAGGAATGTAGTGGCGGAAAAGGAGATAGCAAATGCTTACGATAACGTAAGGAATTATTACTATGTAAACACCAGGCATGATTCTGTTCCGCATAATTACTCCTGATAATTAATCCTTAACTTTGCCCACCTGCCTTTTAAAACATTCCAGTATATCACTTTTCATTCTTGCGTAGCAATATGCCATCTCTTCAGCTATCTCAGCATTGGTGACCTTGTTCAGAGGCGCTGAGAGATGGCCTTTTTCTGATAGATAATGTTCTGTTAAAATATCTCCGGCCTCATCTTTTGCCCGCAGGCTAATGTCTGAAAATTGAGGTGACGGGTTAAAAATAATATCCTTGGCAACCTTTTTTATATCCCTTTTAAATTTTGGCTTAATGACTATATCCAATGAGTCAAAAAGCTCCCCTTCAATATCTGTTGCCCCTAAGACCTTTAATATATCGCCAAATACAGGTAGCTTGGCTTCTACCTTCACCGTTGTTCGGCCGATGAAATGCATATGCATAACATCGTCTTTGGTGGTTCCCCTCATCAGTGGCTCTATCTGAACGCGCTCTCCACTGCTTAATGACATTCCTTTCCCGATTAAAAAATCTGTCAGATCGGATGTGGTCGGCCCGAAAACAGTTCTGGCAAAACCAATGGTGTCGCCTTCAACAAACAAAAAAGATGGGAATCCCAATGATTCGTCATCTGCGAGGCTGTTCTTAATATCTTCAACTGAAGCTTTAGAGCGATTTATCTTCTGAACCAGACTCTTGTCATTTGTTTTGGTAAAGAGAAAAGTTTTTCCATCGATTTTATGAATATACAAATAATTGGAGCCAACCTGCAGGTGATGATTATCAGCCAGCAGAGAATTAAGGAAAACAGACAGGTTTATTGAGCGCTTATCTTTCCCTTTATTTTTGCTGCGGTAAGTCGCATAAAAACCATTCTTCATAATTCAATCCATTTACTATGTTATGTTCTGAGGGGAGTGAAAATTCCCCTAATTCGATGAAGATTCTTGCTCAATTGTTATCAGCTATGCGCCGACCAGAACACCTTGCCGATCAGCCAAACGTCTCTTCAGGCCACTGACTAGCGATAACTTTCCCCACAACGGAACAACTCTCATTGCATGGGATCATTGGGTACTGTGGGTTTAGTGGTTGTAAAAACACCTGACCGCTATCCCTGATCAGTTTCTTGAAGGTAAACTCATCACCCCCAAGTCTGGCTATGCAGAAATCACCTGGCTCAACAGCCTGCTCAGGGTCAACGAGAATTAACATTCCGTCAGGAAAGCTTGGCTTGGAGCCTGTTGGTGCGGTCATGGAATTACCTTCAACCTCAAGCCAGAATGCAGAATCACTGGCTTTTTTGGTTGTGCTTACCCATCTCTCCGCATCACCTTTGGTAAAGGTTCTAAGCTTAGGTGAGAACATCCCTGCCTGAACATGAGAAAAAACAGGGTACTCATACTCACTTCTAAGTGACGGCTGCATACTAACCGCTTCATACATCTCGTAGATTTCTCTGGCGATTGAAGGGCTAAATTCTTCAACGCTAACTTTGAGAATTTTTGTAAGCAATGCGGCGTTATAAGCATTTAATGCATTGATGCCATTAAATAAAGCACCAACGCCTGACTGCCCCATCCCCATCTTGTCTGCGACAGATTCCTGGGATAAGCCAAGTTCATTTTTCTTTTTTTCATAAATTGCTTTAAGGCGACGTGCGTCCTCAAGCTGCTCTTGTGTTAATGGTTTCTTTTTTGTGCTCATACGTTAAATCTATCACCGCAAGGGATAAATATCTAACACCGTGCGTGTTGACTATTTTACCTCTGGCGGTGATAATGGTTGCATGTACTAAGGAGGTTGTATGGAACAACGCATAACCCTGAAAGATTATGCAATGCGCTTTGGGCAAACCAAGACAGCTAAAGATCTCGGCGTATATCAAAGCGCGATCAACAAGGCCATTCATGCAGGCCGAAAGATTTTTTTAACTATAAACGCTGATGGAAGCGTTTATGCGGAAGAGGTAAAGCCCTTCCCGAGTAACAAAAAAACAACAGCATAAATAACCCCGCTCTTACACATTCCAGCCCTGAAAAAGGGCATCAAATTAAACCACACCTATGGTGTATGCATTTATTTGCATACATTCAATCAATTGTTATCTAAGGAAATACTTACATATGGTTCGTGCAAACAAACGCAACGAGGCTCTACGAATCGAGAGTGCGTTGCTTAACAAAATCGCAATGCTTGGAACTGAGAAGACAGCGGAAGCTGTGGGCGTTGATAAGTCGCGTCGAGCCTCAGCTCATGTCATCCTCAGCACACTTGACCCTCAGCTCAGCTAGCCTCAGCCTACAATCACCTCAGCGAATTCGGTGACCCTTACGCGAATCCGCTTTCAGACGTTGACTGGTCGCGTCTGGCAAAAGTTAAAGACCTGACGCCCGGCGAACTGACCGCTGAGTCCTATGACGACAGCTATCTCGATGATGAAGATGCAGACTGGACTGCGACCGGGCAGGGGCAGAAATCTGCCGGAGATACCAGCTTCACGCTGGCGTGGATGCCCGGAGAGCAGGGGCAGCAGGCGCTGCTGGCGTGGTTTAATGAAGGCGATACCCGTGCCTATAAAATCCGCTTCCCGAACGGCACGGTCGATGTGTTCCGTGGCTGGGTCAGCAGTATCGGTAAGGCGGTGACGGCGAAGGAAGTGATCACCCGCACGGTGAAAGTCACCAATGTGGGACGTCCGTCGATGGCAGAAGATCGCAGCACGGTAACAGCGGCAACCGGCATGACCGTGACGCCTGCCAGCACCTCGGTGGTGAAAGGGCAGAGCACCACGCTGACCGTGGCCTTCCAGCCGGAGGGCGTAACCGACAAGAGCTTTCGTGCGGTGTCTGCGGATAAAACAAAAGCCACCGTGTCGGTCAGTGGTATGACCATCACCGTGAACGGCGTTGCTGCAGGCAAGGTCAACATTCCGGTTGTATCCGGTAATGGTGAGTTTGCTGCGGTTGCAGAAATTACCGTCACCGCCAGTTAATCCGGAGAGTCAGCGATGTTCCTGAAAACCGAATCATTTGAACATAACGGTGTGACCGTCACGCTTTCTGAACTGTCAGCCCTGCAGCGCATTGAGCATCTCGCCCTGATGAAACGGCAGGCAGAACAGGCGGAGTCAGACAGCAACCGGAAGTTTACTGTGGAAGACGCCATCAGAACCGGCGCGTTTCTGGTGGCGATGTCCCTGTGGCATAACCATCCGCAGAAGACGCAGATGCCGTCCATGAATGAAGCCGTTAAACAGATTGAGCAGGAAGTGCTTACCACCTGGCCCACGGAGGCAATTTCTCATGCTGAAAACGTGGTGTACCGGCTGTCTGGTATGTATGAGTTTGTGGTGAATAATGCCCCTGAACAGACAGAGGACGCCGGGCCCGCAGAGCCTGTTTCTGCGGGAAAGTGTTCGACGGTGAGCTGAGTTTTGCCCTGAAACTGGCGCGTGAGATGGGGCGACCCGACTGGCGTGCCATGCTTGCCGGGATGTCATCCACGGAGTATGCCGACTGGCACCGCTTTTACAGTACCCATTATTTTCATGATGTTCTGCTGGATATGCACTTTTCCGGGCTGACGTACACCGTGCTCAGCCTGTTTTTCAGCGATCCGGATATGCATCCGCTGGATTTCAGTCTGCTGAACCGGCGCGAGGCTGACGAAGAGCCTGAAGATGATGTGCTGATGCAGAAAGCGGCAGGGCTTGCCGGAGGTGTCCGCTTTGGCCCGGACGGGAATGAAGTTATCCCCGCTTCCCCGGATGTGGCGGACATGACGGAGGATGACGTAATGCTGATGACAGTATCAGAAGGGATCGCAGGAGGAGTCCGGTATGGCTGAACCGGTAGGCGATCTGGTCGTTGATTTGAGTCTGGATGCGGCCAGATTTGACGAGCAGATGGCCAGAGTCAGGCGTCATTTTTCTGGTACGGAAAGTGATGCGAAAAAAACAGCGGCAGTCGTTGAACAGTCGCTGAGCCGACAGGCGCTGGCTGCACAGAAAGCGGGGATTTCCGTCGGGCAGTATAAAGCCGCCATGCGTATGCTGCCTGCACAGTTCACCGACGTGGCCACGCAGCTTGCAGGCGGGCAAAGTCCGTGGCTGATCCTGCTGCAACAGGGGGGGCAGGTGAAGGACTCCTTCGGCGGGATGATCCCCATGTTCAGGGGGCTTGCCGGTGCGATCACCCTGCCGATGGTGGGGGCCACCTCGCTGGCGGTGGCGACCGGTGCGCTGGCGTATGCCTGGTATCAGGGCAACTCAACCCTGTCCGATTTCAACAAAACGCTGGTCCTTTCCGGCAATCAGGCGGGACTGACGGCAGATCGTATGCTGGTCCTGTCCAGAGCCGGGCAGGCGGCAGGGCTGACGTTTAACCAGACCAGCGAGTCACTCAGCGCACTGGTTAAGGCGGGGGTAAGCGGTGAGGCTCAGATTGCGTCCATCAGCCAGAGTGTGGCGCGTTTCTCCTCTGCATCCGGCGTGGAGGTGGACAAGGTCGCTGAAGCCTCTAGAGAATGCTACGTACCTGATGAGCTCCAGCTTTTGTTCCCTTTAGTGAGGGTTAATTGCGCGCTTGGCGTAATCATGGTCATAGCTGTTTCCTGTGTGAAATTGTTATCCGCTCACAATTCCACACAACATACGAGCCGGAAGCATAAAGTGTAAAGCCTGGGGTGCCTAATGAGTGAGCTAACTCACATTAATTGCGTTGCGCTCACTGCCCGCTTTCCAGTCGGGAAACCTGTCGTGCCAGCTGCATTAATGAATCGGCCAACGCGCGGGGAGAGGCGGTTTGCGTATTGGGCGCTCTTCCGCTTCCTCGCTCACTGACTCGCTGCGCTCGGTCGTTCGGCTGCGGCGAGCGGTATCAGCTCACTCAAAGGCGGTAATACGGTTATCCACAGAATCAGGGGATAACGCAGGAAAGAACATGTGAGCAAAAGGCCAGCAAAAGGCCAGGAACCGTAAAAAGGCCGCGTTGCTGGCGTTTTTCCATAGGCTCCGCCCCCCTGACGAGCATCACAAAAATCGACGCTCAAGTCAGAGGTGGCGAAACCCGACAGGACTATAAAGATACCAGGCGTTTCCCCCTGGAAGCTCCCTCGTGCGCTCTCCTGTTCCGACCCTGCCGCTTACCGGATACCTGTCCGCCTTTCTCCCTTCGGGAAGCGTGGCGCTTTCTCATAGCTCACGCTGTAGGTATCTCAGTTCGGTGTAGGTCGTTCGCTCCAAGCTGGGCTGTGTGCACGAACCCCCCGTTCAGCCCGACCGCTGCGCCTTATCCGGTAACTATCGTCTTGAGTCCAACCCGGTAAGACACGACTTATCGCCACTGGCAGCAGCCACTGGTAACAGGATTAGCAGAGCGAGGTATGTAGGCGGTGCTACAGAGTTCTTGAAGTGGTGGCCTAACTACGGCTACACTAGAAGGACAGTATTTGGTATCTGCGCTCTGCTGAAGCCAGTTACCTTCGGAAAAAGAGTTGGTAGCTCTTGATCCGGCAAACAAACCACCGCTGGTAGCGGTGGTTTTTTTGTTTGCAAGCAGCAGATTACGCGCAGAAAAAAAGGATCTCAAGAAGATCCTTTGATCTTTTCTACGGGGTCTGACGCTCAGTGGAACGAAAACTCACGTTAAGGGATTTTGGTCATGAGATTATCAAAAAGGATCTTCACCTAGATCCTTTTAAATTAAAAATGAAGTTTTAAATCAATCTAAAGTATATATGAGTAAACTTGGTCTGACAGTTACCAATGCTTAATCAGTGAGGCACCTATCTCAGCGATCTGTCTATTTCGTTCATCCATAGTTGCCTGACTCCCCGTCGTGTAGATAACTACGATACGGGAGGGCTTACCATCTGGCCCCAGTGCTGCAATGATACCGCGAGACCCACGCTCACCGGCTCCAGATTTATCAGCAATAAACCAGCCAGCCGGAAGGGCCGAGCGCAGAAGTGGTCCTGCAACTTTATCCGCCTCCATCCAGTCTATTAATTGTTGCCGGGAAGCTAGAGTAAGTAGTTCGCCAGTTAATAGTTTGCGCAACGTTGTTGCCATTGCTACAGGCATCGTGGTGTCACGCTCGTCGTTTGGTATGGCTTCATTCAGCTCCGGTTCCCAACGATCAAGGCGAGTTACATGATCCCCCATGTTGTGCAAAAAAGCGGTTAGCTCCTTCGGTCCTCCGATCGTTGTCAGAAGTAAGTTGGCCGCAGTGTTATCACTCATGGTTATGGCAGCACTGCATAATTCTCTTACTGTCATGCCATCCGTAAGATGCTTTTCTGTGACTGGTGAGTACTCAACCAAGTCATTCTGAGAATAGTGTATGCGGCGACCGAGTTGCTCTTGCCCGGCGTCAATACGGGATAATACCGCGCCACATAGCAGAACTTTAAAAGTGCTCATCATTGGAAAACGTTCTTCGGGGCGAAAACTCTCAAGGATCTTACCGCTGTTGAGATCCAGTTCGATGTAACCCACTCGTGCACCCAACTGATCTTCAGCATCTTTTACTTTCACCAGCGTTTCTGGGTGAGCAAAAACAGGAAGGCAAAATGCCGCAAAAAAGGGAATAAGGGCGACACGGAAATGTTGAATACTCATACTCTTCCTTTTTCAATATTATTGAAGCATTTATCAGGGTTATTGTCTCATGAGCGGATACATATTTGAATGTATTTAGAAAAATAAACAAATAGGGGTTCCGCGCACATTTCCCCGAAAAGTGCCACCTAAATTGTAAGCGTTAATATTTTGTTAAAATTCGCGTTAAATTTTTGTTAAATCAGCTCATTTTTTAACCAATAGGCCGAAATCGGCAAAATCCCTTATAAATCAAAAGAATAGACCGAGATAGGGTTGAGTGTTGTTCCAGTTTGGAACAAGAGTCCACTATTAAAGAACGTGGACTCCAACGTCAAAGGGCGAAAAACCGTCTATCAGGGCGATGGCCCACTACGTGAACCATCACCCTAATCAAGTTTTTTGGGGTCGAGGTGCCGTAAAGCACTAAATCGGAACCCTAAAGGGAGCCCCCGATTTAGAGCTTGACGGGGAAAGCCGGCGAACGTGGCGAGAAAGGAAGGGAAGAAAGCGAAAGGAGCGGGCGCTAGGGCGCTGGCAAGTGTAGCGGTCACGCTGCGCGTAACCACCACACCCGCCGCGCTTAATGCGCCGCTACAGGGCGCGTCCCATTCGCCATTCAGGCTGCGCAACTGTTGGGAAGGGCGATCGGTGCGGGCCTCTTCGCTATTACGCCAGCTGGCGAAAGGGGGATGTGCTGCAAGGCGATTAAGTTGGGTAACGCCAGGGTTTTCCCAGTCACGACGTTGTAAAACGACGGCCAGTGAGCGCGCGTAATACGACTCACTATAGGGCGAATTGGGTACCGGCCGACCTCGTGTCTCCAATTGGAGACATCAACGAGGGCGACGCGTACCAGTTCAGGAAGCGGTGATGCTGATAGAAGCCGGACTGAGTACCTACGAGAAAGAGTGCGCAAAACGCGGTGACGACTATCAGGAAATTTTTGCCCAGCAGGTCCGTGAAACGATGGAGCGCCGTGCAGCCGGTCTTAAACCGCCCGCCTGGGCGGCTGCAGCATTTGAATCCGGGCTGCGACAATCAACAGAGGAGGAGAAGAGTGACAGCAGAGCTGCGTAATCTCCCGCATATTGCCAGCATGGCCTTTAATGAGCCGCTGATGCTTGAACCCGCCTATGCGCGGGTTTTCTTTTGTGCGCTTGCAGGCCAGCTTGGGATCAGCAGCCTGGCGGATGCGGTGTCCGGCGACAGCCTGACTGCCCAGGAGGCACTCGGCCCGTGTCTCCAATTGGAGACATCAAGTATAGAGCACCTGTTACGTACCTAGTCAAGGACTCAGCGTGTCTCCAATTGGAGACATCGTTCGACGCTGGCATTATCCGGTGATGATGACGGACCACGACAGGCCCGCAGTTATCAGGTCATGAACGGCATCGCCGTGCTGCCGGTGTCCGGCACGCTGGTCAGCCGGACGCGGGCGCTGCAGCCGTACTCGGGGATGACCGGTTACAACGGCATTATCGCCCGTCTGCAACAGGCTGCCAGCGATCCGATGGTGGACGGCATTCTGCTCGATATGGACACGCCCGGCGGGATGGTGGCGGGGGCATTTGACTGCGCTGACATCATCGCCCGTGTGCGTGACATAAAACCGGTATGGGCGCTTGCCAACGACATGAACTGCAGTGCAGGTCAGTTGCTTGCCAGTGCCGCCTCCCGGCGTCTGGTCACGCAGACCGCCCGGACAGGCTCCATCGGCGTCATGATGGCTCACAGTAATTACGGTGCTGCGCTGGAGAAACAGGGTGTGGAAATCACGCTGATTTACAGCGGCAGCCATAAGGTGGATGGCAACCCCTACAGCCATCTTCCGGATGACGTCCGGGAGACACTGCAGTCCCGGATGGACGCAACCCGCCAGATGTTTGCGCAGAAGGTGTCGGCATATACCGGCCTGTCCGTGCAGGTTGTGCTGGATACCGAGGCTGCAGTGTACAGCGGTCAGGAGGCCATTGATGCCGGACTGGCTGATGAACTTGTTAACAGCACCGATGCGATCACCGTCATGCGTGATGCACTGGATGCACGTAAATCCCGTCTCTCAGGAGGGCGAATGACCAAAGAGACTCAATCAACAACTGTTTCAGCCACTGCTTCGCAGGCTGACGTTACTGACGTGGTGCCAGCGACGGAGGGCGAGAACGCCAGCGCGGCGCAGCCGGACGTGAACGCGCAGATCACCGCAGCGGTTGCGGCAGAAAACAGCCGCATTATGGGGATCCTCAACTGTGAGGAGGCTCACGGACGCGAAGAACAGGCACGCGTGCTGGCAGAAACCCCCGGTATGACCGTGAAAACGGCCCGCCGCATTCTGGCCGCAGCACCACAGAGTGCACAGGCGCGCAGTGACACTGCGCTGGATCGTCTGATGCAGGGGGCACCGGCACCGCTGGCTGCAGGTAACCCGGCATCTGATGCCGTTAACGATTTGCTGAACACACCAGTGTAAGGGATGTTTATGACGAGCAAAGAAACCTTTACCCATTACCAGCCGCAGGGCAACAGTGACCCGGCTCATACCGCAACCGCGCCCGGCGGATTGAGTGCGAAAGCGCCTGCAATGACCCCGCTGATGCTGGACACCTCCAGCCGTAAGCTGGTTGCGTGGGATGGCACCACCGACGGTGCTGCCGTTGGCATTCTTGCGGTTGCTGCTGACCAGACCAGCACCACGCTGACGTTCTACAAGTCCGGCACGTTCCGTTATGAGGATGTGCTCTGGCCGGAGGCTGCCAGCGACGAGACGAAAAAACGGACCGCGTTTGCCGGAACGGCAATCAGCATCGTTTAACTTTACCCTTCATCACTAAAGGCCGCCTGTGCGGCTTTTTTTACGGGATTTTTTTATGTCGATGTACACAACCGCCCAACTGCTGGCGGCAAATGAGCAGAAATTTAAGTTTGATCCGCTGTTTCTGCGTCTCTTTTTCCGTGAGAGCTATCCCTTCACCACGGAGAAAGTCTATCTCTCACAAATTCCGGGACTGGTAAACATGGCGCTGTACGTTTCGCCGATTGTTTCCGGTGAGGTTATCCGTTCCCGTGGCGGCTCCACCTCTGAATTTACGCCGGGATATGTCAAGCCGAAGCATGAAGTGAATCCGCAGATGACCCTGCGTCGCCTGCCGGATGAAGATCCGCAGAATCTGGCGGACCCGGCTTACCGCCGCCGTCGCATCATCATGCAGAACATGCGTGACGAAGAGCTGGCCATTGCTCAGGTCGAAGAGATGCAGGCAGTTTCTGCCGTGCTTAAGGGCAAATACACCATGACCGGTGAAGCCTTCGATCCGGTTGAGGTGGATATGGGCCGCAGTGAGGAGAATAACATCACGCAGTCCGGCGGCACGGAGTGGAGCAAGCGTGACAAGTCCACGTATGACCCGACCGACGATATCGAAGCCTACGCGCTGAACGCCAGCGGTGTGGTGAATATCATCGTGTTCGATCCGAAAGGCTGGGCGCTGTTCCGTTCCTTCAAAGCCGTCAAGGAGAAGCTGGATACCCGTCGTGGCTCTAATTCCGAGCTGGAGACAGCGGTGAAAGACCTGGGCAAAGCGGTGTCCTATAAGGGGATGTATGGCGATGTGGCCATCGTCGTGTATTCCGGACAGTACGTGGAAAACGGCGTCAAAAAGAACTTCCTGCCGGACAACACGATGGTGCTGGGGAACACTCAGGCACGCGGTCTGCGCACCTATGGCTGCATTCAGGATGCGGACGCACAGCGCGAAGGCATTAACGCCTCTGCCCGTTACCCGAAAAACTGGGTGACCACCGGCGATCCGGCGCGTGAGTTCACCATGATTCAGTCAGCACCGCTGATGCTGCTGGCTGACCCTGATGAGTTCGTGTCCGTACAACTGGCGTAATCATGGCCCTTCGGGGCCATTGTTTCTCTGTGGAGGAGTCCATGACGAAAGATGAACTGATTGCCCGTCTCCGCTCGCTGGGTGAACAACTGAACCGTGATGTCAGCCTGACGGGGACGAAAGAAGAACTGGCGCTCCGTGTGGCAGAGCTGAAAGAGGAGCTTGATGACACGGATGAAACTGCCGGTCAGGACACCCCTCTCAGCCGGGAAAATGTGCTGACCGGACATGAAAATGAGGTGGGATCAGCGCAGCCGGATACCGTGATTCTGGATACGTCTGAACTGGTCACGGTCGTGGCACTGGTGAAGCTGCATACTGATGCACTTCACGCCACGCGGGATGAACCTGTGGCATTTGTGCTGCCGGGAACGGCGTTTCGTGTCTCTGCCGGTGTGGCAGCCGAAATGACAGAGCGCGGCCTGGCCAGAATGCAATAACGGGAGGCGCTGTGGCTGATTTCGATAACCTGTTCGATGCTGCCATTGCCCGCGCCGATGAAACGATACGCGGGTACATGGGAACGTCAGCCACCATTACATCCGGTGAGCAGTCAGGTGCGGTGATACGTGGTGTTTTTGATGACCCTGAAAATATCAGCTATGCCGGACAGGGCGTGCGCGTTGAAGGCTCCAGCCCGTCCCTGTTTGTCCGGACTGATGAGGTGCGGCAGCTGCGGCGTGGAGACACGCTGACCATCGGTGAGGAAAATTTCTGGGTAGATCGGGTTTCGCCGGATGATGGCGGAAGTTGTCATCTCTGGCTTGGACGGGGCGTACCGCCTGCCGTTAACCGTCGCCGCTGAAAGGGGGATGTATGGCCATAAAAGGTCTTGAGCAGGCCGTTGAAAACCTCAGCCGTATCAGCAAAACGGCGGTGCCTGGTGCCGCCGCAATGGCCATTAACCGCGTTGCTTCATCCGCGATATCGCAGTCGGCGTCACAGGTTGCCCGTGAGACAAAGGTACGCCGGAAACTGGTAAAGGAAAGGGCCAGGCTGAAAAGGGCCACGGTCAAAAATCCGCAGGCCAGAATCAAAGTTAACCGGGGGGATTTGCCCGTAATCAAGCTGGGTAATGCGCGGGTTGTCCTTTCGCGCCGCAGGCGTCGTAAAAAGGGGCAGCGTTCATCCCTGAAAGGTGGCGGCAGCGTGCTTGTGGTGGGTAACCGTCGTATTCCCGGCGCGTTTATTCAGCAACTGAAAAATGGCCGGTGGCATGTCATGCAGCGTGTGGCTGGGAAAAACCGTTACCCCATTGATGTGGTGAAAATCCCGATGGCGGTGCCGCTGACCACGGCGTTTAAACAAAATATTGAGCGGATACGGCGTGAACGTCTTCCGAAAGAGCTGGGCTATGCGCTGCAGCATCAACTGAGGATGGTAATAAAGCGATGAAACATACTGAACTCCGTGCAGCCGTACTGGATGCACTGGAGAAGCATGACACCGGGGCGACGTTTTTTGATGGTCGCCCCGCTGTTTTTGATGAGGCGGATTTTCCGGCAGTTGCCGTTTATCTCACCGGCGCTGAATACACGGGCGAAGAGCTGGACAGCGATACCTGGCAGGCGGAGCTGCATATCGAAGTTTTCCTGCCTGCTCAGGTGCCGGATTCAGAGCTGGATGCGTGGATGGAGTCCCGGATTTATCCGGTGATGAGCGATATCCCGGCACTGTCAGATTTGATCACCAGTATGGTGGCCAGCGGCTATGACTACCGGCGCGACGATGATGCGGGCTTGTGGAGTTCAGCCGATCTGACTTATGTCATTACCTATGAAATGTGAGGACGCTATGCCTGTACCAAATCCTACAATGCCGGTGAAAGGTGCCGGGACCACCCTGTGGGTTTATAAGGGGAGCGGTGACCCTTACGCGAATCCGCTTTCAGACGTTGACTGGTCGCGTCTGGCAAAAGTTAAAGACCTGACGCCCGGCGAACTGACCGCTGAGTCCTATGACGACAGCTATCTCGATGATGAAGATGCAGACTGGACTGCGACCGGGCAGGGGCAGAAATCTGCCGGAGATACCAGCTTCACGCTGGCGTGGATGCCCGGAGAGCAGGGGCAGCAGGCGCTGCTGGCGTGGTTTAATGAAGGCGATACCCGTGCCTATAAAATCCGCTTCCCGAACGGCACGGTCGATGTGTTCCGTGGCTGGGTCAGCAGTATCGGTAAGGCGGTGACGGCGAAGGAAGTGATCACCCGCACGGTGAAAGTCACCAATGTGGGACGTCCGTCGATGGCAGAAGATCGCAGCACGGTAACAGCGGCAACCGGCATGACCGTGACGCCTGCCAGCACCTCGGTGGTGAAAGGGCAGAGCACCACGCTGACCGTGGCCTTCCAGCCGGAGGGCGTAACCGACAAGAGCTTTCGTGCGGTGTCTGCGGATAAAACAAAAGCCACCGTGTCGGTCAGTGGTATGACCATCACCGTGAACGGCGTTGCTGCAGGCAAGGTCAACATTCCGGTTGTATCCGGTAATGGTGAGTTTGCTGCGGTTGCAGAAATTACCGTCACCGCCAGTTAATCCGGAGAGTCAGCGATGTTCCTGAAAACCGAATCATTTGAACATAACGGTGTGACCGTCACGCTTTCTGAACTGTCAGCCCTGCAGCGCATTGAGCATCTCGCCCTGATGAAACGGCAGGCAGAACAGGCGGAGTCAGACAGCAACCGGAAGTTTACTGTGGAAGACGCCATCAGAACCGGCGCGTTTCTGGTGGCGATGTCCCTGTGGCATAACCATCCGCAGAAGACGCAGATGCCGTCCATGAATGAAGCCGTTAAACAGATTGAGCAGGAAGTGCTTACCACCTGGCCCACGGAGGCAATTTCTCATGCTGAAAACGTGGTGTACCGGCTGTCTGGTATGTATGAGTTTGTGGTGAATAATGCCCCTGAACAGACAGAGGACGCCGGGCCCGCAGAGCCTGTTTCTGCGGGAAAGTGTTCGACGGTGAGCTGAGTTTTGCCCTGAAACTGGCGCGTGAGATGGGGCGACCCGACTGGCGTGCCATGCTTGCCGGGATGTCATCCACGGAGTATGCCGACTGGCACCGCTTTTACAGTACCCATTATTTTCATGATGTTCTGCTGGATATGCACTTTTCCGGGCTGACGTACACCGTGCTCAGCCTGTTTTTCAGCGATCCGGATATGCATCCGCTGGATTTCAGTCTGCTGAACCGGCGCGAGGCTGACGAAGAGCCTGAAGATGATGTGCTGATGCAGAAAGCGGCAGGGCTTGCCGGAGGTGTCCGCTTTGGCCCGGACGGGAATGAAGTTATCCCCGCTTCCCCGGATGTGGCGGACATGACGGAGGATGACGTAATGCTGATGACAGTATCAGAAGGGATCGCAGGAGGAGTCCGGTATGGCTGAACCGGTAGGCGATCTGGTCGTTGATTTGAGTCTGGATGCGGCCAGATTTGACGAGCAGATGGCCAGAGTCAGGCGTCATTTTTCTGGTACGGAAAGTGATGCGAAAAAAACAGCGGCAGTCGTTGAACAGTCGCTGAGCCGACAGGCGCTGGCTGCACAGAAAGCGGGGATTTCCGTCGGGCAGTATAAAGCCGCCATGCGTATGCTGCCTGCACAGTTCACCGACGTGGCCACGCAGCTTGCAGGCGGGCAAAGTCCGTGGCTGATCCTGCTGCAACAGGGGGGGCAGGTGAAGGACTCCTTCGGCGGGATGATCCCCATGTTCAGGGGGCTTGCCGGTGCGATCACCCTGCCGATGGTGGGGGCCACCTCGCTGGCGGTGGCGACCGGTGCGCTGGCGTATGCCTGGTATCAGGGCAACTCAACCCTGTCCGATTTCAACAAAACGCTGGTCCTTTCCGGCAATCAGGCGGGACTGACGGCAGATCGTATGCTGGTCCTGTCCAGAGCCGGGCAGGCGGCAGGGCTGACGTTTAACCAGACCAGCGAGTCACTCAGCGCACTGGTTAAGGCGGGGGTAAGCGGTGAGGCTCAGATTGCGTCCATCAGCCAGAGTGTGGCGCGTTTCTCCTCTGCATCCGGCGTGGAGGTGGACAAGGTCGCTGAAGCCTTCGGGAAGCTGACCACAGACCCGACGTCGGGGCTGACGGCGATGGCTCGCCAGTTCCATAACGTGTCGGCGGAGCAGATTGCGTATGTTGCTCAGTTGCAGCGTTCCGGCGATGAAGCCGGGGCATTGCAGGCGGCGAACGAGGCCGCAACGAAAGGGTTTGATGACCAGACCCGCCGCCTGAAAGAGAACATGGGCACGCTGGAGACCTGGGCAGACAGGACTGCGCGGGCATTCAAATCCATGTGGGATGCGGTGCTGGATATTGGTCGTCCTGATACCGCGCAGGAGATGCTGATTAAGGCAGAGGCTGCGTATAAGAAAGCAGACGACATCTGGAATCTGCGCAAGGATGATTATTTTGTTAACGATGAAGCGCGGGCGCGTTACTGGGATGATCGTGAAAAGGCCCGTCTTGCGCTTGAAGCCGCCCGAAAGAAGGCTGAGCAGCAGACTCAACAGGACAAAAATGCGCAGCAGCAGAGCGATACCGAAGCGTCACGGCTGAAATATACCGAAGAGGCGCAGAAGGCTTACGAACGGCTGCAGACGCCGCTGGAGAAATATACCGCCCGTCAGGAAGAACTGAACAAGGCACTGAAAGACGGGAAAATCCTGCAGGCGGATTACAACACGCTGATGGCGGCGGCGAAAAAGGATTATGAAGCGACGCTGAAAAAGCCGAAACAGTCCAGCGTGAAGGTGTCTGCGGGCGATCGTCAGGAAGACAGTGCTCATGCTGCCCTGCTGACGCTTCAGGCAGAACTCCGGACGCTGGAGAAGCATGCCGGAGCAAATGAGAAAATCAGCCAGCAGCGCCGGGATTTGTGGAAGGCGGAGAGTCAGTTCGCGGTACTGGAGGAGGCGGCGCAACGTCGCCAGCTGTCTGCACAGGAGAAATCCCTGCTGGCGCATAAAGATGAGACGCTGGAGTACAAACGCCAGCTGGCTGCACTTGGCGACAAGGTTACGTATCAGGAGCGCCTGAACGCGCTGGCGCAGCAGGCGGATAAATTCGCACAGCAGCAACGGGCAAAACGGGCCGCCATTGATGCGAAAAGCCGGGGGCTGACTGACCGGCAGGCAGAACGGGAAGCCACGGAACAGCGCCTGAAGGAACAGTATGGCGATAATCCGCTGGCGCTGAATAACGTCATGTCAGAGCAGAAAAAGACCTGGGCGGCTGAAGACCAGCTTCGCGGGAACTGGATGGCAGGCCTGAAGTCCGGCTGGAGTGAGTGGGAAGAGAGCGCCACGGACAGTATGTCGCAGGTAAAAAGTGCAGCCACGCAGACCTTTGATGGTATTGCACAGAATATGGCGGCGATGCTGACCGGCAGTGAGCAGAACTGGCGCAGCTTCACCCGTTCCGTGCTGTCCATGATGACAGAAATTCTGCTTAAGCAGGCAATGGTGGGGATTGTCGGGAGTATCGGCAGCGCCATTGGCGGGGCTGTTGGTGGCGGCGCATCCGCGTCAGGCGGTACAGCCATTCAGGCCGCTGCGGCGAAATTCCATTTTGCAACCGGAGGATTTACGGGAACCGGCGGCAAATATGAGCCAGCGGGGATTGTTCACCGTGGTGAGTTTGTCTTCACGAAGGAGGCAACCAGCCGGATTGGCGTGGGGAATCTTTACCGGCTGATGCGCGGCTATGCCACCGGCGGTTATGTCGGTACACCGGGCAGCATGGCAGACAGCCGGTCGCAGGCGTCCGGGACGTTTGAGCAGAATAACCATGTGGTGATTAACAACGACGGCACGAACGGGCAGATAGGTCCGGCTGCTCTGAAGGCGGTGTATGACATGGCCCGCAAGGGTGCCCGTGATGAAATTCAGACACAGATGCGTGATGGTGGCCTGTTCTCCGGAGGTGGACGATGAAGACCTTCCGCTGGAAAGTGAAACCCGGTATGGATGTGGCTTCGGTCCCTTCTGTAAGAAAGGTGCGCTTTGGTGATGGCTATTCTCAGCGAGCGCCTGCCGGGCTGAATGCCAACCTGAAAACGTACAGCGTGACGCTTTCTGTCCCCCGTGAGGAGGCCACGGTACTGGAGTCGTTTCTGGAAGAGCACGGGGGCTGGAAATCCTTTCTGTGGACGCCGCCTTATGAGTGGCGGCAGATAAAGGTGACCTGCGCAAAATGGTCGTCGCGGGTCAGTATGCTGCGTGTTGAGTTCAGCGCAGAGTTTGAACAGGTGGTGAACTGATGCAGGATATCCGGCAGGAAACACTGAATGAATGCACCCGTGCGGAGCAGTCGGCCAGCGTGGTGCTCTGGGAAATCGACCTGACAGAGGTCGGTGGAGAACGTTATTTTTTCTGTAATGAGCAGAACGAAAAAGGTGAGCCGGTCACCTGGCAGGGGCGACAGTATCAGCCGTATCCCATTCAGGGGAGCGGTTTTGAACTGAATGGCAAAGGCACCAGTACGCGCCCCACGCTGACGGTTTCTAACCTGTACGGTATGGTCACCGGGATGGCGGAAGATATGCAGAGTCTGGTCGGCGGAACGGTGGTCCGGCGTAAGGTTTACGCCCGTTTTCTGGATGCGGTGAACTTCGTCAACGGAAACAGTTACGCCGATCCGGAGCAGGAGGTGATCAGCCGCTGGCGCATTGAGCAGTGCAGCGAACTGAGCGCGGTGAGTGCCTCCTTTGTACTGTCCACGCCGACGGAAACGGATGGCGCTGTTTTTCCGGGACGTATCATGCTGGCCAACACCTGCACCTGGACCTATCGCGGTGACGAGTGCGGTTATAGCGGTCCGGCTGTCGCGGATGAATATGACCAGCCAACGTCCGATATCACGAAGGATAAATGCAGCAAATGCCTGAGCGGTTGTAAGTTCCGCAATAACGTCGGCAACTTTGGCGGCTTCCTTTCCATTAACAAACTTTCGCAGTAAATCCCATGACACAGACAGAATCAGCGATTCTGGCGCACGCCCGGCGATGTGCGCCAGCGGAGTCGTGCGGCTTCGTGGTAAGCACGCCGGAGGGGGAAAGATATTTCCCCTGCGTGAATATCTCCGGTGAGCCGGAGGCGTATTTCCGTATGTCGCCGGAAGACTGGCTGCAGGCAGAAATGCAGGGTGAGATTGTGGCGCTGGTCCACAGCCACCCCGGTGGTCTGCCCTGGCTGAGTGAGGCCGACCGGCGGCTGCAGGTGCAGAGTGATTTGCCGTGGTGGCTGGTCTGCCGGGGGACGATTCATAAGTTCCGCTGTGTGCCGCATCTCACCGGGCGGCGCTTTGAGCACGGTGTGACGGACTGTTACACACTGTTCCGGGC |
| C-trap central fragment without *parS* sites  (20482 bp) | GGCCGCGGTGTGCTCCTTATTTATACATAACGAAAAACGCCTCGAGTGAAGCGTTATTGGTATGCGGTAAAACCGCACTCAGGCGGCCTTGATAGTCATATCATCTGAATCAAATATTCCTGATGTATCGATATCGGTAATTCTTATTCCTTCGCTACCATCCATTGGAGGCCATCCTTCCTGACCATTTCCATCATTCCAGTCGAACTCACACACAACACCATATGCATTTAAGTCGCTTGAAATTGCTATAAGCAGAGCATGTTGCGCCAGCATGATTAATACAGCATTTAATACAGAGCCGTGTTTATTGAGTCGGTATTCAGAGTCTGACCAGAAATTATTAATCTGGTGAAGTTTTTCCTCTGTCATTACGTCATGGTCGATTTCAATTTCTATTGATGCTTTCCAGTCGTAATCAATGATGTATTTTTTGATGTTTGACATCTGTTCATATCCTCACAGATAAAAAATCGCCCTCACACTGGAGGGCAAAGAAGATTTCCAATAATCAGAACAAGTCGGCTCCTGTTTAGTTACGAGCGACATTGCTCCGTGTATTCACTCGTTGGAATGAATACACAGTGCAGTGTTTATTCTGTTATTTATGCCAAAAATAAAGGCCACTATCAGGCAGCTTTGTTGTTCTGTTTACCAAGTTCTCTGGCAATCATTGCCGTCGTTCGTATTGCCCATTTATCGACATATTTCCCATCTTCCATTACAGGAAACATTTCTTCAGGCTTAACCATGCATTCCGATTGCAGCTTGCATCCATTGCATCGCTTGAATTGTCCACACCATTGATTTTTATCAATAGTCGTAGTCATACGGATAGTCCTGGTATTGTTCCATCACATCCTGAGGATGCTCTTCGAACTCTTCAAATTCTTCTTCCATATATCACCTTAAATAGTGGATTGCGGTAGTAAAGATTGTGCCTGTCTTTTAACCACATCAGGCTCGGTGGTTCTCGTGTACCCCTACAGCGAGAAATCGGATAAACTATTACAACCCCTACAGTTTGATGAGTATAGAAATGGATCCACTCGTTATTCTCGGACGAGTGTTCAGTAATGAACCTCTGGAGAGAACCATGTATATGATCGTTATCTGGGTTGGACTTCTGCTTTTAAGCCCAGATAACTGGCCTGAATATGTTAATGAGAGAATCGGTATTCCTCATGTGTGGCATGTTTTCGTCTTTGCTCTTGCATTTTCGCTAGCAATTAATGTGCATCGATTATCAGCTATTGCCAGCGCCAGATATAAGCGATTTAAGCTAAGAAAACGCATTAAGATGCAAAACGATAAAGTGCGATCAGTAATTCAAAACCTTACAGAAGAGCAATCTATGGTTTTGTGCGCAGCCCTTAATGAAGGCAGGAAGTATGTGGTTACATCAAAACAATTCCCATACATTAGTGAGTTGATTGAGCTTGGTGTGTTGAACAAAACTTTTTCCCGATGGAATGGAAAGCATATATTATTCCCTATTGAGGATATTTACTGGACTGAATTAGTTGCCAGCTATGATCCATATAATATTGAGATAAAGCCAAGGCCAATATCTAAGTAACTAGATAAGAGGAATCGATTTTCCCTTAATTTTCTGGCGTCCACTGCATGTTATGCCGCGTTCGCCAGGCTTGCTGTACCATGTGCGCTGATTCTTGCGCTCAATACGTTGCAGGTTGCTTTCAATCTGTTTGTGGTATTCAGCCAGCACTGTAAGGTCTATCGGATTTAGTGCGCTTTCTACTCGTGATTTCGGTTTGCGATTCAGCGAGAGAATAGGGCGGTTAACTGGTTTTGCGCTTACCCCAACCAACAGGGGATTTGCTGCTTTCCATTGAGCCTGTTTCTCTGCGCGACGTTCGCGGCGGCGTGTTTGTGCATCCATCTGGATTCTCCTGTCAGTTAGCTTTGGTGGTGTGTGGCAGTTGTAGTCCTGAACGAAAACCCCCCGCGATTGGCACATTGGCAGCTAATCCGGAATCGCACTTACGGCCAATGCTTCGTTTCGTATCACACACCCCAAAGCCTTCTGCTTTGAATGCTGCCCTTCTTCAGGGCTTAATTTTTAAGAGCGTCACCTTCATGGTGGTCAGTGCGTCCTGCTGATGTGCTCAGTATCACCGCCAGTGGTATTTATGTCAACACCGCCAGAGATAATTTATCACCGCAGATGGTTATCTGTATGTTTTTTATATGAATTTATTTTTTGCAGGGGGGCATTGTTTGGTAGGTGAGAGATCTGAATTGCTATGTTTAGTGAGTTGTATCTATTTATTTTTCAATAAATACAATTGGTTATGTGTTTTGGGGGCGATCGTGAGGCAAAGAAAACCCGGCGCTGAGGCCGGGTTATTCTTGTTCTCTGGTCAAATTATATAGTTGGAAAACAAGGATGCATATATGAATGAACGATGCAGAGGCAATGCCGATGGCGATAGTGGGTATCATGTAGCCGCTTATGCTGGAAAGAAGCAATAACCCGCAGAAAAACAAAGCTCCAAGCTCAACAAAACTAAGGGCATAGACAATAACTACCGATGTCATATACCCATACTCTCTAATCTTGGCCAGTCGGCGCGTTCTGCTTCCGATTAGAAACGTCAAGGCAGCAATCAGGATTGCAATCATGGTTCCTGCATATGATGACAATGTCGCCCCAAGACCATCTCTATGAGCTGAAAAAGAAACACCAGGAATGTAGTGGCGGAAAAGGAGATAGCAAATGCTTACGATAACGTAAGGAATTATTACTATGTAAACACCAGGCATGATTCTGTTCCGCATAATTACTCCTGATAATTAATCCTTAACTTTGCCCACCTGCCTTTTAAAACATTCCAGTATATCACTTTTCATTCTTGCGTAGCAATATGCCATCTCTTCAGCTATCTCAGCATTGGTGACCTTGTTCAGAGGCGCTGAGAGATGGCCTTTTTCTGATAGATAATGTTCTGTTAAAATATCTCCGGCCTCATCTTTTGCCCGCAGGCTAATGTCTGAAAATTGAGGTGACGGGTTAAAAATAATATCCTTGGCAACCTTTTTTATATCCCTTTTAAATTTTGGCTTAATGACTATATCCAATGAGTCAAAAAGCTCCCCTTCAATATCTGTTGCCCCTAAGACCTTTAATATATCGCCAAATACAGGTAGCTTGGCTTCTACCTTCACCGTTGTTCGGCCGATGAAATGCATATGCATAACATCGTCTTTGGTGGTTCCCCTCATCAGTGGCTCTATCTGAACGCGCTCTCCACTGCTTAATGACATTCCTTTCCCGATTAAAAAATCTGTCAGATCGGATGTGGTCGGCCCGAAAACAGTTCTGGCAAAACCAATGGTGTCGCCTTCAACAAACAAAAAAGATGGGAATCCCAATGATTCGTCATCTGCGAGGCTGTTCTTAATATCTTCAACTGAAGCTTTAGAGCGATTTATCTTCTGAACCAGACTCTTGTCATTTGTTTTGGTAAAGAGAAAAGTTTTTCCATCGATTTTATGAATATACAAATAATTGGAGCCAACCTGCAGGTGATGATTATCAGCCAGCAGAGAATTAAGGAAAACAGACAGGTTTATTGAGCGCTTATCTTTCCCTTTATTTTTGCTGCGGTAAGTCGCATAAAAACCATTCTTCATAATTCAATCCATTTACTATGTTATGTTCTGAGGGGAGTGAAAATTCCCCTAATTCGATGAAGATTCTTGCTCAATTGTTATCAGCTATGCGCCGACCAGAACACCTTGCCGATCAGCCAAACGTCTCTTCAGGCCACTGACTAGCGATAACTTTCCCCACAACGGAACAACTCTCATTGCATGGGATCATTGGGTACTGTGGGTTTAGTGGTTGTAAAAACACCTGACCGCTATCCCTGATCAGTTTCTTGAAGGTAAACTCATCACCCCCAAGTCTGGCTATGCAGAAATCACCTGGCTCAACAGCCTGCTCAGGGTCAACGAGAATTAACATTCCGTCAGGAAAGCTTGGCTTGGAGCCTGTTGGTGCGGTCATGGAATTACCTTCAACCTCAAGCCAGAATGCAGAATCACTGGCTTTTTTGGTTGTGCTTACCCATCTCTCCGCATCACCTTTGGTAAAGGTTCTAAGCTTAGGTGAGAACATCCCTGCCTGAACATGAGAAAAAACAGGGTACTCATACTCACTTCTAAGTGACGGCTGCATACTAACCGCTTCATACATCTCGTAGATTTCTCTGGCGATTGAAGGGCTAAATTCTTCAACGCTAACTTTGAGAATTTTTGTAAGCAATGCGGCGTTATAAGCATTTAATGCATTGATGCCATTAAATAAAGCACCAACGCCTGACTGCCCCATCCCCATCTTGTCTGCGACAGATTCCTGGGATAAGCCAAGTTCATTTTTCTTTTTTTCATAAATTGCTTTAAGGCGACGTGCGTCCTCAAGCTGCTCTTGTGTTAATGGTTTCTTTTTTGTGCTCATACGTTAAATCTATCACCGCAAGGGATAAATATCTAACACCGTGCGTGTTGACTATTTTACCTCTGGCGGTGATAATGGTTGCATGTACTAAGGAGGTTGTATGGAACAACGCATAACCCTGAAAGATTATGCAATGCGCTTTGGGCAAACCAAGACAGCTAAAGATCTCGGCGTATATCAAAGCGCGATCAACAAGGCCATTCATGCAGGCCGAAAGATTTTTTTAACTATAAACGCTGATGGAAGCGTTTATGCGGAAGAGGTAAAGCCCTTCCCGAGTAACAAAAAAACAACAGCATAAATAACCCCGCTCTTACACATTCCAGCCCTGAAAAAGGGCATCAAATTAAACCACACCTATGGTGTATGCATTTATTTGCATACATTCAATCAATTGTTATCTAAGGAAATACTTACATATGGTTCGTGCAAACAAACGCAACGAGGCTCTACGAATCGAGAGTGCGTTGCTTAACAAAATCGCAATGCTTGGAACTGAGAAGACAGCGGAAGCTGTGGGCGTTGATAAGTCGCGTCGAGCCTCAGCTCATGTCATCCTCAGCACACTTGACCCTCAGCTCAGCTAGCCTCAGCCTACAATCACCTCAGCGAATTCGGTGACCCTTACGCGAATCCGCTTTCAGACGTTGACTGGTCGCGTCTGGCAAAAGTTAAAGACCTGACGCCCGGCGAACTGACCGCTGAGTCCTATGACGACAGCTATCTCGATGATGAAGATGCAGACTGGACTGCGACCGGGCAGGGGCAGAAATCTGCCGGAGATACCAGCTTCACGCTGGCGTGGATGCCCGGAGAGCAGGGGCAGCAGGCGCTGCTGGCGTGGTTTAATGAAGGCGATACCCGTGCCTATAAAATCCGCTTCCCGAACGGCACGGTCGATGTGTTCCGTGGCTGGGTCAGCAGTATCGGTAAGGCGGTGACGGCGAAGGAAGTGATCACCCGCACGGTGAAAGTCACCAATGTGGGACGTCCGTCGATGGCAGAAGATCGCAGCACGGTAACAGCGGCAACCGGCATGACCGTGACGCCTGCCAGCACCTCGGTGGTGAAAGGGCAGAGCACCACGCTGACCGTGGCCTTCCAGCCGGAGGGCGTAACCGACAAGAGCTTTCGTGCGGTGTCTGCGGATAAAACAAAAGCCACCGTGTCGGTCAGTGGTATGACCATCACCGTGAACGGCGTTGCTGCAGGCAAGGTCAACATTCCGGTTGTATCCGGTAATGGTGAGTTTGCTGCGGTTGCAGAAATTACCGTCACCGCCAGTTAATCCGGAGAGTCAGCGATGTTCCTGAAAACCGAATCATTTGAACATAACGGTGTGACCGTCACGCTTTCTGAACTGTCAGCCCTGCAGCGCATTGAGCATCTCGCCCTGATGAAACGGCAGGCAGAACAGGCGGAGTCAGACAGCAACCGGAAGTTTACTGTGGAAGACGCCATCAGAACCGGCGCGTTTCTGGTGGCGATGTCCCTGTGGCATAACCATCCGCAGAAGACGCAGATGCCGTCCATGAATGAAGCCGTTAAACAGATTGAGCAGGAAGTGCTTACCACCTGGCCCACGGAGGCAATTTCTCATGCTGAAAACGTGGTGTACCGGCTGTCTGGTATGTATGAGTTTGTGGTGAATAATGCCCCTGAACAGACAGAGGACGCCGGGCCCGCAGAGCCTGTTTCTGCGGGAAAGTGTTCGACGGTGAGCTGAGTTTTGCCCTGAAACTGGCGCGTGAGATGGGGCGACCCGACTGGCGTGCCATGCTTGCCGGGATGTCATCCACGGAGTATGCCGACTGGCACCGCTTTTACAGTACCCATTATTTTCATGATGTTCTGCTGGATATGCACTTTTCCGGGCTGACGTACACCGTGCTCAGCCTGTTTTTCAGCGATCCGGATATGCATCCGCTGGATTTCAGTCTGCTGAACCGGCGCGAGGCTGACGAAGAGCCTGAAGATGATGTGCTGATGCAGAAAGCGGCAGGGCTTGCCGGAGGTGTCCGCTTTGGCCCGGACGGGAATGAAGTTATCCCCGCTTCCCCGGATGTGGCGGACATGACGGAGGATGACGTAATGCTGATGACAGTATCAGAAGGGATCGCAGGAGGAGTCCGGTATGGCTGAACCGGTAGGCGATCTGGTCGTTGATTTGAGTCTGGATGCGGCCAGATTTGACGAGCAGATGGCCAGAGTCAGGCGTCATTTTTCTGGTACGGAAAGTGATGCGAAAAAAACAGCGGCAGTCGTTGAACAGTCGCTGAGCCGACAGGCGCTGGCTGCACAGAAAGCGGGGATTTCCGTCGGGCAGTATAAAGCCGCCATGCGTATGCTGCCTGCACAGTTCACCGACGTGGCCACGCAGCTTGCAGGCGGGCAAAGTCCGTGGCTGATCCTGCTGCAACAGGGGGGGCAGGTGAAGGACTCCTTCGGCGGGATGATCCCCATGTTCAGGGGGCTTGCCGGTGCGATCACCCTGCCGATGGTGGGGGCCACCTCGCTGGCGGTGGCGACCGGTGCGCTGGCGTATGCCTGGTATCAGGGCAACTCAACCCTGTCCGATTTCAACAAAACGCTGGTCCTTTCCGGCAATCAGGCGGGACTGACGGCAGATCGTATGCTGGTCCTGTCCAGAGCCGGGCAGGCGGCAGGGCTGACGTTTAACCAGACCAGCGAGTCACTCAGCGCACTGGTTAAGGCGGGGGTAAGCGGTGAGGCTCAGATTGCGTCCATCAGCCAGAGTGTGGCGCGTTTCTCCTCTGCATCCGGCGTGGAGGTGGACAAGGTCGCTGAAGCCTCTAGAGAATGCTACGTACCTGATGAGCTCCAGCTTTTGTTCCCTTTAGTGAGGGTTAATTGCGCGCTTGGCGTAATCATGGTCATAGCTGTTTCCTGTGTGAAATTGTTATCCGCTCACAATTCCACACAACATACGAGCCGGAAGCATAAAGTGTAAAGCCTGGGGTGCCTAATGAGTGAGCTAACTCACATTAATTGCGTTGCGCTCACTGCCCGCTTTCCAGTCGGGAAACCTGTCGTGCCAGCTGCATTAATGAATCGGCCAACGCGCGGGGAGAGGCGGTTTGCGTATTGGGCGCTCTTCCGCTTCCTCGCTCACTGACTCGCTGCGCTCGGTCGTTCGGCTGCGGCGAGCGGTATCAGCTCACTCAAAGGCGGTAATACGGTTATCCACAGAATCAGGGGATAACGCAGGAAAGAACATGTGAGCAAAAGGCCAGCAAAAGGCCAGGAACCGTAAAAAGGCCGCGTTGCTGGCGTTTTTCCATAGGCTCCGCCCCCCTGACGAGCATCACAAAAATCGACGCTCAAGTCAGAGGTGGCGAAACCCGACAGGACTATAAAGATACCAGGCGTTTCCCCCTGGAAGCTCCCTCGTGCGCTCTCCTGTTCCGACCCTGCCGCTTACCGGATACCTGTCCGCCTTTCTCCCTTCGGGAAGCGTGGCGCTTTCTCATAGCTCACGCTGTAGGTATCTCAGTTCGGTGTAGGTCGTTCGCTCCAAGCTGGGCTGTGTGCACGAACCCCCCGTTCAGCCCGACCGCTGCGCCTTATCCGGTAACTATCGTCTTGAGTCCAACCCGGTAAGACACGACTTATCGCCACTGGCAGCAGCCACTGGTAACAGGATTAGCAGAGCGAGGTATGTAGGCGGTGCTACAGAGTTCTTGAAGTGGTGGCCTAACTACGGCTACACTAGAAGGACAGTATTTGGTATCTGCGCTCTGCTGAAGCCAGTTACCTTCGGAAAAAGAGTTGGTAGCTCTTGATCCGGCAAACAAACCACCGCTGGTAGCGGTGGTTTTTTTGTTTGCAAGCAGCAGATTACGCGCAGAAAAAAAGGATCTCAAGAAGATCCTTTGATCTTTTCTACGGGGTCTGACGCTCAGTGGAACGAAAACTCACGTTAAGGGATTTTGGTCATGAGATTATCAAAAAGGATCTTCACCTAGATCCTTTTAAATTAAAAATGAAGTTTTAAATCAATCTAAAGTATATATGAGTAAACTTGGTCTGACAGTTACCAATGCTTAATCAGTGAGGCACCTATCTCAGCGATCTGTCTATTTCGTTCATCCATAGTTGCCTGACTCCCCGTCGTGTAGATAACTACGATACGGGAGGGCTTACCATCTGGCCCCAGTGCTGCAATGATACCGCGAGACCCACGCTCACCGGCTCCAGATTTATCAGCAATAAACCAGCCAGCCGGAAGGGCCGAGCGCAGAAGTGGTCCTGCAACTTTATCCGCCTCCATCCAGTCTATTAATTGTTGCCGGGAAGCTAGAGTAAGTAGTTCGCCAGTTAATAGTTTGCGCAACGTTGTTGCCATTGCTACAGGCATCGTGGTGTCACGCTCGTCGTTTGGTATGGCTTCATTCAGCTCCGGTTCCCAACGATCAAGGCGAGTTACATGATCCCCCATGTTGTGCAAAAAAGCGGTTAGCTCCTTCGGTCCTCCGATCGTTGTCAGAAGTAAGTTGGCCGCAGTGTTATCACTCATGGTTATGGCAGCACTGCATAATTCTCTTACTGTCATGCCATCCGTAAGATGCTTTTCTGTGACTGGTGAGTACTCAACCAAGTCATTCTGAGAATAGTGTATGCGGCGACCGAGTTGCTCTTGCCCGGCGTCAATACGGGATAATACCGCGCCACATAGCAGAACTTTAAAAGTGCTCATCATTGGAAAACGTTCTTCGGGGCGAAAACTCTCAAGGATCTTACCGCTGTTGAGATCCAGTTCGATGTAACCCACTCGTGCACCCAACTGATCTTCAGCATCTTTTACTTTCACCAGCGTTTCTGGGTGAGCAAAAACAGGAAGGCAAAATGCCGCAAAAAAGGGAATAAGGGCGACACGGAAATGTTGAATACTCATACTCTTCCTTTTTCAATATTATTGAAGCATTTATCAGGGTTATTGTCTCATGAGCGGATACATATTTGAATGTATTTAGAAAAATAAACAAATAGGGGTTCCGCGCACATTTCCCCGAAAAGTGCCACCTAAATTGTAAGCGTTAATATTTTGTTAAAATTCGCGTTAAATTTTTGTTAAATCAGCTCATTTTTTAACCAATAGGCCGAAATCGGCAAAATCCCTTATAAATCAAAAGAATAGACCGAGATAGGGTTGAGTGTTGTTCCAGTTTGGAACAAGAGTCCACTATTAAAGAACGTGGACTCCAACGTCAAAGGGCGAAAAACCGTCTATCAGGGCGATGGCCCACTACGTGAACCATCACCCTAATCAAGTTTTTTGGGGTCGAGGTGCCGTAAAGCACTAAATCGGAACCCTAAAGGGAGCCCCCGATTTAGAGCTTGACGGGGAAAGCCGGCGAACGTGGCGAGAAAGGAAGGGAAGAAAGCGAAAGGAGCGGGCGCTAGGGCGCTGGCAAGTGTAGCGGTCACGCTGCGCGTAACCACCACACCCGCCGCGCTTAATGCGCCGCTACAGGGCGCGTCCCATTCGCCATTCAGGCTGCGCAACTGTTGGGAAGGGCGATCGGTGCGGGCCTCTTCGCTATTACGCCAGCTGGCGAAAGGGGGATGTGCTGCAAGGCGATTAAGTTGGGTAACGCCAGGGTTTTCCCAGTCACGACGTTGTAAAACGACGGCCAGTGAGCGCGCGTAATACGACTCACTATAGGGCGAATTGGGTACCAGTTCAGGAAGCGGTGATGCTGATAGAAGCCGGACTGAGTACCTACGAGAAAGAGTGCGCAAAACGCGGTGACGACTATCAGGAAATTTTTGCCCAGCAGGTCCGTGAAACGATGGAGCGCCGTGCAGCCGGTCTTAAACCGCCCGCCTGGGCGGCTGCAGCATTTGAATCCGGGCTGCGACAATCAACAGAGGAGGAGAAGAGTGACAGCAGAGCTGCGTAATCTCCCGCATATTGCCAGCATGGCCTTTAATGAGCCGCTGATGCTTGAACCCGCCTATGCGCGGGTTTTCTTTTGTGCGCTTGCAGGCCAGCTTGGGATCAGCAGCCTGGCGGATGCGGTGTCCGGCGACAGCCTGACTGCCCAGGAGGCACTCGCGACGCTGGCATTATCCGGTGATGATGACGGACCACGACAGGCCCGCAGTTATCAGGTCATGAACGGCATCGCCGTGCTGCCGGTGTCCGGCACGCTGGTCAGCCGGACGCGGGCGCTGCAGCCGTACTCGGGGATGACCGGTTACAACGGCATTATCGCCCGTCTGCAACAGGCTGCCAGCGATCCGATGGTGGACGGCATTCTGCTCGATATGGACACGCCCGGCGGGATGGTGGCGGGGGCATTTGACTGCGCTGACATCATCGCCCGTGTGCGTGACATAAAACCGGTATGGGCGCTTGCCAACGACATGAACTGCAGTGCAGGTCAGTTGCTTGCCAGTGCCGCCTCCCGGCGTCTGGTCACGCAGACCGCCCGGACAGGCTCCATCGGCGTCATGATGGCTCACAGTAATTACGGTGCTGCGCTGGAGAAACAGGGTGTGGAAATCACGCTGATTTACAGCGGCAGCCATAAGGTGGATGGCAACCCCTACAGCCATCTTCCGGATGACGTCCGGGAGACACTGCAGTCCCGGATGGACGCAACCCGCCAGATGTTTGCGCAGAAGGTGTCGGCATATACCGGCCTGTCCGTGCAGGTTGTGCTGGATACCGAGGCTGCAGTGTACAGCGGTCAGGAGGCCATTGATGCCGGACTGGCTGATGAACTTGTTAACAGCACCGATGCGATCACCGTCATGCGTGATGCACTGGATGCACGTAAATCCCGTCTCTCAGGAGGGCGAATGACCAAAGAGACTCAATCAACAACTGTTTCAGCCACTGCTTCGCAGGCTGACGTTACTGACGTGGTGCCAGCGACGGAGGGCGAGAACGCCAGCGCGGCGCAGCCGGACGTGAACGCGCAGATCACCGCAGCGGTTGCGGCAGAAAACAGCCGCATTATGGGGATCCTCAACTGTGAGGAGGCTCACGGACGCGAAGAACAGGCACGCGTGCTGGCAGAAACCCCCGGTATGACCGTGAAAACGGCCCGCCGCATTCTGGCCGCAGCACCACAGAGTGCACAGGCGCGCAGTGACACTGCGCTGGATCGTCTGATGCAGGGGGCACCGGCACCGCTGGCTGCAGGTAACCCGGCATCTGATGCCGTTAACGATTTGCTGAACACACCAGTGTAAGGGATGTTTATGACGAGCAAAGAAACCTTTACCCATTACCAGCCGCAGGGCAACAGTGACCCGGCTCATACCGCAACCGCGCCCGGCGGATTGAGTGCGAAAGCGCCTGCAATGACCCCGCTGATGCTGGACACCTCCAGCCGTAAGCTGGTTGCGTGGGATGGCACCACCGACGGTGCTGCCGTTGGCATTCTTGCGGTTGCTGCTGACCAGACCAGCACCACGCTGACGTTCTACAAGTCCGGCACGTTCCGTTATGAGGATGTGCTCTGGCCGGAGGCTGCCAGCGACGAGACGAAAAAACGGACCGCGTTTGCCGGAACGGCAATCAGCATCGTTTAACTTTACCCTTCATCACTAAAGGCCGCCTGTGCGGCTTTTTTTACGGGATTTTTTTATGTCGATGTACACAACCGCCCAACTGCTGGCGGCAAATGAGCAGAAATTTAAGTTTGATCCGCTGTTTCTGCGTCTCTTTTTCCGTGAGAGCTATCCCTTCACCACGGAGAAAGTCTATCTCTCACAAATTCCGGGACTGGTAAACATGGCGCTGTACGTTTCGCCGATTGTTTCCGGTGAGGTTATCCGTTCCCGTGGCGGCTCCACCTCTGAATTTACGCCGGGATATGTCAAGCCGAAGCATGAAGTGAATCCGCAGATGACCCTGCGTCGCCTGCCGGATGAAGATCCGCAGAATCTGGCGGACCCGGCTTACCGCCGCCGTCGCATCATCATGCAGAACATGCGTGACGAAGAGCTGGCCATTGCTCAGGTCGAAGAGATGCAGGCAGTTTCTGCCGTGCTTAAGGGCAAATACACCATGACCGGTGAAGCCTTCGATCCGGTTGAGGTGGATATGGGCCGCAGTGAGGAGAATAACATCACGCAGTCCGGCGGCACGGAGTGGAGCAAGCGTGACAAGTCCACGTATGACCCGACCGACGATATCGAAGCCTACGCGCTGAACGCCAGCGGTGTGGTGAATATCATCGTGTTCGATCCGAAAGGCTGGGCGCTGTTCCGTTCCTTCAAAGCCGTCAAGGAGAAGCTGGATACCCGTCGTGGCTCTAATTCCGAGCTGGAGACAGCGGTGAAAGACCTGGGCAAAGCGGTGTCCTATAAGGGGATGTATGGCGATGTGGCCATCGTCGTGTATTCCGGACAGTACGTGGAAAACGGCGTCAAAAAGAACTTCCTGCCGGACAACACGATGGTGCTGGGGAACACTCAGGCACGCGGTCTGCGCACCTATGGCTGCATTCAGGATGCGGACGCACAGCGCGAAGGCATTAACGCCTCTGCCCGTTACCCGAAAAACTGGGTGACCACCGGCGATCCGGCGCGTGAGTTCACCATGATTCAGTCAGCACCGCTGATGCTGCTGGCTGACCCTGATGAGTTCGTGTCCGTACAACTGGCGTAATCATGGCCCTTCGGGGCCATTGTTTCTCTGTGGAGGAGTCCATGACGAAAGATGAACTGATTGCCCGTCTCCGCTCGCTGGGTGAACAACTGAACCGTGATGTCAGCCTGACGGGGACGAAAGAAGAACTGGCGCTCCGTGTGGCAGAGCTGAAAGAGGAGCTTGATGACACGGATGAAACTGCCGGTCAGGACACCCCTCTCAGCCGGGAAAATGTGCTGACCGGACATGAAAATGAGGTGGGATCAGCGCAGCCGGATACCGTGATTCTGGATACGTCTGAACTGGTCACGGTCGTGGCACTGGTGAAGCTGCATACTGATGCACTTCACGCCACGCGGGATGAACCTGTGGCATTTGTGCTGCCGGGAACGGCGTTTCGTGTCTCTGCCGGTGTGGCAGCCGAAATGACAGAGCGCGGCCTGGCCAGAATGCAATAACGGGAGGCGCTGTGGCTGATTTCGATAACCTGTTCGATGCTGCCATTGCCCGCGCCGATGAAACGATACGCGGGTACATGGGAACGTCAGCCACCATTACATCCGGTGAGCAGTCAGGTGCGGTGATACGTGGTGTTTTTGATGACCCTGAAAATATCAGCTATGCCGGACAGGGCGTGCGCGTTGAAGGCTCCAGCCCGTCCCTGTTTGTCCGGACTGATGAGGTGCGGCAGCTGCGGCGTGGAGACACGCTGACCATCGGTGAGGAAAATTTCTGGGTAGATCGGGTTTCGCCGGATGATGGCGGAAGTTGTCATCTCTGGCTTGGACGGGGCGTACCGCCTGCCGTTAACCGTCGCCGCTGAAAGGGGGATGTATGGCCATAAAAGGTCTTGAGCAGGCCGTTGAAAACCTCAGCCGTATCAGCAAAACGGCGGTGCCTGGTGCCGCCGCAATGGCCATTAACCGCGTTGCTTCATCCGCGATATCGCAGTCGGCGTCACAGGTTGCCCGTGAGACAAAGGTACGCCGGAAACTGGTAAAGGAAAGGGCCAGGCTGAAAAGGGCCACGGTCAAAAATCCGCAGGCCAGAATCAAAGTTAACCGGGGGGATTTGCCCGTAATCAAGCTGGGTAATGCGCGGGTTGTCCTTTCGCGCCGCAGGCGTCGTAAAAAGGGGCAGCGTTCATCCCTGAAAGGTGGCGGCAGCGTGCTTGTGGTGGGTAACCGTCGTATTCCCGGCGCGTTTATTCAGCAACTGAAAAATGGCCGGTGGCATGTCATGCAGCGTGTGGCTGGGAAAAACCGTTACCCCATTGATGTGGTGAAAATCCCGATGGCGGTGCCGCTGACCACGGCGTTTAAACAAAATATTGAGCGGATACGGCGTGAACGTCTTCCGAAAGAGCTGGGCTATGCGCTGCAGCATCAACTGAGGATGGTAATAAAGCGATGAAACATACTGAACTCCGTGCAGCCGTACTGGATGCACTGGAGAAGCATGACACCGGGGCGACGTTTTTTGATGGTCGCCCCGCTGTTTTTGATGAGGCGGATTTTCCGGCAGTTGCCGTTTATCTCACCGGCGCTGAATACACGGGCGAAGAGCTGGACAGCGATACCTGGCAGGCGGAGCTGCATATCGAAGTTTTCCTGCCTGCTCAGGTGCCGGATTCAGAGCTGGATGCGTGGATGGAGTCCCGGATTTATCCGGTGATGAGCGATATCCCGGCACTGTCAGATTTGATCACCAGTATGGTGGCCAGCGGCTATGACTACCGGCGCGACGATGATGCGGGCTTGTGGAGTTCAGCCGATCTGACTTATGTCATTACCTATGAAATGTGAGGACGCTATGCCTGTACCAAATCCTACAATGCCGGTGAAAGGTGCCGGGACCACCCTGTGGGTTTATAAGGGGAGCGGTGACCCTTACGCGAATCCGCTTTCAGACGTTGACTGGTCGCGTCTGGCAAAAGTTAAAGACCTGACGCCCGGCGAACTGACCGCTGAGTCCTATGACGACAGCTATCTCGATGATGAAGATGCAGACTGGACTGCGACCGGGCAGGGGCAGAAATCTGCCGGAGATACCAGCTTCACGCTGGCGTGGATGCCCGGAGAGCAGGGGCAGCAGGCGCTGCTGGCGTGGTTTAATGAAGGCGATACCCGTGCCTATAAAATCCGCTTCCCGAACGGCACGGTCGATGTGTTCCGTGGCTGGGTCAGCAGTATCGGTAAGGCGGTGACGGCGAAGGAAGTGATCACCCGCACGGTGAAAGTCACCAATGTGGGACGTCCGTCGATGGCAGAAGATCGCAGCACGGTAACAGCGGCAACCGGCATGACCGTGACGCCTGCCAGCACCTCGGTGGTGAAAGGGCAGAGCACCACGCTGACCGTGGCCTTCCAGCCGGAGGGCGTAACCGACAAGAGCTTTCGTGCGGTGTCTGCGGATAAAACAAAAGCCACCGTGTCGGTCAGTGGTATGACCATCACCGTGAACGGCGTTGCTGCAGGCAAGGTCAACATTCCGGTTGTATCCGGTAATGGTGAGTTTGCTGCGGTTGCAGAAATTACCGTCACCGCCAGTTAATCCGGAGAGTCAGCGATGTTCCTGAAAACCGAATCATTTGAACATAACGGTGTGACCGTCACGCTTTCTGAACTGTCAGCCCTGCAGCGCATTGAGCATCTCGCCCTGATGAAACGGCAGGCAGAACAGGCGGAGTCAGACAGCAACCGGAAGTTTACTGTGGAAGACGCCATCAGAACCGGCGCGTTTCTGGTGGCGATGTCCCTGTGGCATAACCATCCGCAGAAGACGCAGATGCCGTCCATGAATGAAGCCGTTAAACAGATTGAGCAGGAAGTGCTTACCACCTGGCCCACGGAGGCAATTTCTCATGCTGAAAACGTGGTGTACCGGCTGTCTGGTATGTATGAGTTTGTGGTGAATAATGCCCCTGAACAGACAGAGGACGCCGGGCCCGCAGAGCCTGTTTCTGCGGGAAAGTGTTCGACGGTGAGCTGAGTTTTGCCCTGAAACTGGCGCGTGAGATGGGGCGACCCGACTGGCGTGCCATGCTTGCCGGGATGTCATCCACGGAGTATGCCGACTGGCACCGCTTTTACAGTACCCATTATTTTCATGATGTTCTGCTGGATATGCACTTTTCCGGGCTGACGTACACCGTGCTCAGCCTGTTTTTCAGCGATCCGGATATGCATCCGCTGGATTTCAGTCTGCTGAACCGGCGCGAGGCTGACGAAGAGCCTGAAGATGATGTGCTGATGCAGAAAGCGGCAGGGCTTGCCGGAGGTGTCCGCTTTGGCCCGGACGGGAATGAAGTTATCCCCGCTTCCCCGGATGTGGCGGACATGACGGAGGATGACGTAATGCTGATGACAGTATCAGAAGGGATCGCAGGAGGAGTCCGGTATGGCTGAACCGGTAGGCGATCTGGTCGTTGATTTGAGTCTGGATGCGGCCAGATTTGACGAGCAGATGGCCAGAGTCAGGCGTCATTTTTCTGGTACGGAAAGTGATGCGAAAAAAACAGCGGCAGTCGTTGAACAGTCGCTGAGCCGACAGGCGCTGGCTGCACAGAAAGCGGGGATTTCCGTCGGGCAGTATAAAGCCGCCATGCGTATGCTGCCTGCACAGTTCACCGACGTGGCCACGCAGCTTGCAGGCGGGCAAAGTCCGTGGCTGATCCTGCTGCAACAGGGGGGGCAGGTGAAGGACTCCTTCGGCGGGATGATCCCCATGTTCAGGGGGCTTGCCGGTGCGATCACCCTGCCGATGGTGGGGGCCACCTCGCTGGCGGTGGCGACCGGTGCGCTGGCGTATGCCTGGTATCAGGGCAACTCAACCCTGTCCGATTTCAACAAAACGCTGGTCCTTTCCGGCAATCAGGCGGGACTGACGGCAGATCGTATGCTGGTCCTGTCCAGAGCCGGGCAGGCGGCAGGGCTGACGTTTAACCAGACCAGCGAGTCACTCAGCGCACTGGTTAAGGCGGGGGTAAGCGGTGAGGCTCAGATTGCGTCCATCAGCCAGAGTGTGGCGCGTTTCTCCTCTGCATCCGGCGTGGAGGTGGACAAGGTCGCTGAAGCCTTCGGGAAGCTGACCACAGACCCGACGTCGGGGCTGACGGCGATGGCTCGCCAGTTCCATAACGTGTCGGCGGAGCAGATTGCGTATGTTGCTCAGTTGCAGCGTTCCGGCGATGAAGCCGGGGCATTGCAGGCGGCGAACGAGGCCGCAACGAAAGGGTTTGATGACCAGACCCGCCGCCTGAAAGAGAACATGGGCACGCTGGAGACCTGGGCAGACAGGACTGCGCGGGCATTCAAATCCATGTGGGATGCGGTGCTGGATATTGGTCGTCCTGATACCGCGCAGGAGATGCTGATTAAGGCAGAGGCTGCGTATAAGAAAGCAGACGACATCTGGAATCTGCGCAAGGATGATTATTTTGTTAACGATGAAGCGCGGGCGCGTTACTGGGATGATCGTGAAAAGGCCCGTCTTGCGCTTGAAGCCGCCCGAAAGAAGGCTGAGCAGCAGACTCAACAGGACAAAAATGCGCAGCAGCAGAGCGATACCGAAGCGTCACGGCTGAAATATACCGAAGAGGCGCAGAAGGCTTACGAACGGCTGCAGACGCCGCTGGAGAAATATACCGCCCGTCAGGAAGAACTGAACAAGGCACTGAAAGACGGGAAAATCCTGCAGGCGGATTACAACACGCTGATGGCGGCGGCGAAAAAGGATTATGAAGCGACGCTGAAAAAGCCGAAACAGTCCAGCGTGAAGGTGTCTGCGGGCGATCGTCAGGAAGACAGTGCTCATGCTGCCCTGCTGACGCTTCAGGCAGAACTCCGGACGCTGGAGAAGCATGCCGGAGCAAATGAGAAAATCAGCCAGCAGCGCCGGGATTTGTGGAAGGCGGAGAGTCAGTTCGCGGTACTGGAGGAGGCGGCGCAACGTCGCCAGCTGTCTGCACAGGAGAAATCCCTGCTGGCGCATAAAGATGAGACGCTGGAGTACAAACGCCAGCTGGCTGCACTTGGCGACAAGGTTACGTATCAGGAGCGCCTGAACGCGCTGGCGCAGCAGGCGGATAAATTCGCACAGCAGCAACGGGCAAAACGGGCCGCCATTGATGCGAAAAGCCGGGGGCTGACTGACCGGCAGGCAGAACGGGAAGCCACGGAACAGCGCCTGAAGGAACAGTATGGCGATAATCCGCTGGCGCTGAATAACGTCATGTCAGAGCAGAAAAAGACCTGGGCGGCTGAAGACCAGCTTCGCGGGAACTGGATGGCAGGCCTGAAGTCCGGCTGGAGTGAGTGGGAAGAGAGCGCCACGGACAGTATGTCGCAGGTAAAAAGTGCAGCCACGCAGACCTTTGATGGTATTGCACAGAATATGGCGGCGATGCTGACCGGCAGTGAGCAGAACTGGCGCAGCTTCACCCGTTCCGTGCTGTCCATGATGACAGAAATTCTGCTTAAGCAGGCAATGGTGGGGATTGTCGGGAGTATCGGCAGCGCCATTGGCGGGGCTGTTGGTGGCGGCGCATCCGCGTCAGGCGGTACAGCCATTCAGGCCGCTGCGGCGAAATTCCATTTTGCAACCGGAGGATTTACGGGAACCGGCGGCAAATATGAGCCAGCGGGGATTGTTCACCGTGGTGAGTTTGTCTTCACGAAGGAGGCAACCAGCCGGATTGGCGTGGGGAATCTTTACCGGCTGATGCGCGGCTATGCCACCGGCGGTTATGTCGGTACACCGGGCAGCATGGCAGACAGCCGGTCGCAGGCGTCCGGGACGTTTGAGCAGAATAACCATGTGGTGATTAACAACGACGGCACGAACGGGCAGATAGGTCCGGCTGCTCTGAAGGCGGTGTATGACATGGCCCGCAAGGGTGCCCGTGATGAAATTCAGACACAGATGCGTGATGGTGGCCTGTTCTCCGGAGGTGGACGATGAAGACCTTCCGCTGGAAAGTGAAACCCGGTATGGATGTGGCTTCGGTCCCTTCTGTAAGAAAGGTGCGCTTTGGTGATGGCTATTCTCAGCGAGCGCCTGCCGGGCTGAATGCCAACCTGAAAACGTACAGCGTGACGCTTTCTGTCCCCCGTGAGGAGGCCACGGTACTGGAGTCGTTTCTGGAAGAGCACGGGGGCTGGAAATCCTTTCTGTGGACGCCGCCTTATGAGTGGCGGCAGATAAAGGTGACCTGCGCAAAATGGTCGTCGCGGGTCAGTATGCTGCGTGTTGAGTTCAGCGCAGAGTTTGAACAGGTGGTGAACTGATGCAGGATATCCGGCAGGAAACACTGAATGAATGCACCCGTGCGGAGCAGTCGGCCAGCGTGGTGCTCTGGGAAATCGACCTGACAGAGGTCGGTGGAGAACGTTATTTTTTCTGTAATGAGCAGAACGAAAAAGGTGAGCCGGTCACCTGGCAGGGGCGACAGTATCAGCCGTATCCCATTCAGGGGAGCGGTTTTGAACTGAATGGCAAAGGCACCAGTACGCGCCCCACGCTGACGGTTTCTAACCTGTACGGTATGGTCACCGGGATGGCGGAAGATATGCAGAGTCTGGTCGGCGGAACGGTGGTCCGGCGTAAGGTTTACGCCCGTTTTCTGGATGCGGTGAACTTCGTCAACGGAAACAGTTACGCCGATCCGGAGCAGGAGGTGATCAGCCGCTGGCGCATTGAGCAGTGCAGCGAACTGAGCGCGGTGAGTGCCTCCTTTGTACTGTCCACGCCGACGGAAACGGATGGCGCTGTTTTTCCGGGACGTATCATGCTGGCCAACACCTGCACCTGGACCTATCGCGGTGACGAGTGCGGTTATAGCGGTCCGGCTGTCGCGGATGAATATGACCAGCCAACGTCCGATATCACGAAGGATAAATGCAGCAAATGCCTGAGCGGTTGTAAGTTCCGCAATAACGTCGGCAACTTTGGCGGCTTCCTTTCCATTAACAAACTTTCGCAGTAAATCCCATGACACAGACAGAATCAGCGATTCTGGCGCACGCCCGGCGATGTGCGCCAGCGGAGTCGTGCGGCTTCGTGGTAAGCACGCCGGAGGGGGAAAGATATTTCCCCTGCGTGAATATCTCCGGTGAGCCGGAGGCGTATTTCCGTATGTCGCCGGAAGACTGGCTGCAGGCAGAAATGCAGGGTGAGATTGTGGCGCTGGTCCACAGCCACCCCGGTGGTCTGCCCTGGCTGAGTGAGGCCGACCGGCGGCTGCAGGTGCAGAGTGATTTGCCGTGGTGGCTGGTCTGCCGGGGGACGATTCATAAGTTCCGCTGTGTGCCGCATCTCACCGGGCGGCGCTTTGAGCACGGTGTGACGGACTGTTACACACTGTTCCGGGC |
